# Exploiting TGF-β-mediated Stromal Programming in Homologous Recombination-Deficient Pancreatic Cancer

**DOI:** 10.1101/2025.09.16.676458

**Authors:** Elodie Roger, Hannah M. Mummey, Eleni Zimmer, Dharini Srinivasan, Anna Härle, Loïc Moubri, Alica K. Beutel, Rima Singh, Menar Ekizce, Michael K. Melzer, Yun Lee, Austin Silva, Luisa Härle, Thomas Engleitner, Frank Arnold, Mareen Morawe, Benjamin Naggay, Juliane Schneider, Leonard Gilberg, Julia P. Mosler, Christina Ludwig, Chen Meng, Maximilian Hirschenberger, Victoria Hunszinger, Klaus Kluck, Martina Kirchner, Anna-Lena Volckmar, Mareike Wirth, Mohamed S.S. Alhamdani, Jörg D. Hoheisel, J.-Matthias Löhr, Thomas Seufferlein, Alireza Abaei, Ralf Kemkemer, Roland Rad, Jan Budczies, Medhanie A. Mulaw, Patrick C. Hermann, Mark Hänle, Konstantin M. J. Sparrer, Christopher J. Halbrook, Kyle J. Gaulton, Konrad Steinestel, Albrecht Stenzinger, Nelson Dusetti, Lukas Perkhofer, Johann Gout, Alexander Kleger

## Abstract

The tumor microenvironment (TME) actively contributes to pancreatic ductal adenocarcinoma (PDAC) pathogenesis through dynamic bidirectional tumor–stroma interactions. Here, we demonstrate that homologous recombination-defective (HRD) tumor epithelium reprograms the TME in a genotype-specific manner to enhance cancer aggressiveness. Using genetically engineered mouse models, pancreatic stellate cell (PSC) and cancer-associated fibroblast (CAF) co-culture systems, single-nucleus multiomics, and human PDAC models, we show that tumoral loss of ATM serine/threonine kinase drives CAFs toward αSMA^+^ myofibroblastic differentiation, independently of P53 status. These myCAFs, in turn, promote cancer aggressiveness and chemoresistance. Mechanistically, ATM deficiency increases reactive oxygen species and contractility signaling, enhancing TGF-β1 secretion. Pharmacological TGF-β inhibition reverses myCAF differentiation, sensitizes tumors to chemotherapy, and impairs tumor progression in both murine and human ATM-null models. Our findings reveal that ATM-deficient tumors shape a cancer-promoting niche via TGF-β signaling and identify dual targeting of intrinsic and extrinsic vulnerabilities as a promising precision oncology strategy.

**SIGNIFICANCE:** HRD pancreatic cancers reprogram the tumor microenvironment in a genotype-specific manner through TGF-β-driven myCAF-enrichment. Targeting this stromal axis alongside platinum-based chemotherapy improves therapeutic efficacy in ATM-deficient models. These findings highlight the need to integrate epithelial genotype and stromal context for truly personalized treatment strategies in PDAC.

## INTRODUCTION

Pancreatic ductal adenocarcinoma (PDAC) remains one of the deadliest solid malignancies, with a dismal five-year overall survival rate that just recently reached 13% among all stages (1). In advanced disease stage, most treatments are not employed based on molecular tumor characteristics but instead on patients’ performance (2,3). The only exception is the PARP inhibitor olaparib, a DNA-damaging drug (4,5), which operates synthetically lethal with germline *BRCA1/2* mutations and in a set of other DNA damage response genes (DDR) (4,6–10). Apart from this, targeted therapies have mostly failed underpinning PDAC’s complex, hard-to-treat cancer biology (11). These overall poor prognosis and treatment outcomes are largely attributed to the high heterogeneity of both the neoplastic epithelium and its tumor microenvironment (TME) (12). The TME is composed of cancer-associated fibroblasts (CAFs), immune, endothelial, neuronal cells, and matrix proteins. The distinct TME components, their heterogeneity, and their spatial distribution impact behavior and plasticity of individual PDACs (13,14). CAFs, the most prominent cellular TME component, exhibit diverse phenotypes and functional features which define co-existing distinct subpopulations (15–21). αSMA^+^ myofibroblastic CAFs (myCAFs) reside mainly in close vicinity of cancer cells and are associated with extracellular matrix (ECM) production and fibrosis development, whereas distant inflammatory CAFs (iCAFs) are associated with a cytokine secretory phenotype (16). A third subset, named antigen-presenting CAFs (apCAFs), show high expression of MHC class II family proteins and participate to tumor immunosuppression by inducing regulatory T cell formation (17,18). However, the precise functions of each subtype in promoting or restricting PDAC progression and response to therapy remains a vibrant research field. ECM-producing αSMA^+^ myCAFs restrain cancer progression by constituting a physical barrier limiting metastatic dissemination and tumor angiogenesis (19,20,22). Conversely, CAF-secreted dense ECM network prevents drug delivery (23). Beyond this mechanical constraint, myCAF-derived factors support tumor progression notably through fostering cell proliferation and invasion (21,24). Nevertheless, underlying molecular mechanisms and the impact of the neoplastic epithelial genotype remain poorly understood. Consequently, previous global and non-selective stroma-targeting approaches not only failed to improve but even worsened clinical outcome (19,23,25). *In vivo* studies revealed that chemical inhibition of the JAK/STAT pathway, a key driver of the iCAF phenotype, promoted myCAF enrichment. This stromal shift ultimately led to reduced tumor growth in PDAC-bearing mice (26). These observations highlight the need for more selective strategies that distinguish between tumor-promoting and tumor-restraining stromal components. Although an extensive desmoplastic reaction systematically accompanies PDAC development, the contribution of a particular PDAC genotype in orchestrating stromal compartment programming remains poorly understood and does not translate into efficient therapeutic approaches. Deciphering mutation-specific epithelium–stroma molecular communications would thus further guide the elaboration of novel personalized therapies interfering into tumor-promoting crosstalks. A particularly aggressive PDAC subtype, defined as genomically unstable with a higher mutational burden, frequently harbors DDR gene mutations and is associated with sensitivity toward platinum-based chemotherapies (27,28). Pathogenic germline or somatic DDR mutations occur in 17-25% of patients, most commonly affecting *ATM*, *BRCA1*/2, or *PALB2* (10,29). This molecularly defined subgroup comprises clinically relevant subset of patients who may benefit from stratified therapeutic strategies targeting both epithelial vulnerabilities and genotype-specific stromal features. The *ATM* gene, encoding a key serine/threonine kinase operating in homologous recombination (HR) repair, is the most frequently mutated DDR gene in PDAC (4). We previously showed that its loss generates HR deficiency (HRD) with impaired DNA repair capacity and high genomic instability and causes highly fibrotic and metastatic oncogenic features (4,5,8). Although this highly aggressive subtype significantly limits PDAC outcome, ATM loss-mediated HRD goes alongside with druggable vulnerabilities toward platin derivatives or PARP, ATR, and DNA-PK inhibitors (4,8,9). Here, we show that ATM loss-of-function in neoplastic epithelium drives reprogramming of the surrounding niche to feature cancer aggressiveness through a TGF-β-mediated tumor–stroma crosstalk, constituting an additional layer of druggable vulnerability in HRD PDAC. Thus, epithelial genotypes not only mirror cancer diversity *per se*, but additionally drive TME heterogeneity and need to be considered to design truly personalized therapeutics.

## RESULTS

### ATM loss-of-function accelerates pancreatic cancer tumorigenesis in the absence of P53

P53 plays critical roles to maintain genetic stability (30). In line with the genomically unstable phenotype reported in mouse ATM-deficient PDAC (referred as AKC; (5,8,10)), P53 levels were higher in AKC cells suggesting a significant activation of the P53 checkpoint in these cancers (**Fig. 1A–C**). To test whether P53 restrains the effects of ATM loss-of-function during pancreatic tumorigenesis, we crossed into our *Atm^fl/fl^* (A)*; LSL-Kras^G12D/+^* (K)*; Ptf1a^Cre/+^* (C) mice, a floxed *Trp53* allele (P) to generate the AKPC mouse model (**Fig. 1D**). While the *Trp53* status determined the survival corridor, the additional *Atm* deletion led to a significantly reduced survival in either genotype (AKC vs. KC: 296.0 vs. 550.5 days; AKPC vs. KPC: 55.0 vs. 66.0 days; **Fig. 1E**). Notably, even heterozygous *Atm* loss shortened survival in KPC mice (59.0 days vs. 66.0 days), indicating a hypomorphic effect of the *Atm* allele (**Supplementary Fig. S1A**). To dissect the tumor features arising specifically from the loss of ATM, we conducted a comprehensive immunohistological characterization. High-grade pancreatic intraepithelial neoplasia (PanIN) lesions were more frequent in 5-week-old AKPC mice (**Fig. 1F** and **G**). At later timepoint, ATM deletion led to invasive tumors with undifferentiated features, indicative of a mesenchymal and substantially aggressive phenotype (**Fig. 1H; Supplementary Fig. S1B**). Liver metastasis incidence tended to be higher in AKPC than in their KPC counterparts (**Fig. 1I**). Isolated tumor cells also revealed prominent mesenchymal features upon ATM loss-of-function without detectable differences in proliferation (**Fig. 1J** and **K; Supplementary Fig. S1C** and **S1D**) (5,8). Within the DDR machinery, ATM acts as a safeguard against genomic instability and thereby tumor progression (4). Accordingly, ATM-deficient malignant cells exhibited larger nucleus size, an indication of aneuploidy, and higher levels of DNA damage as highlighted by H2AX p-S139 staining, even in the absence of the P53 checkpoint (4,8) (**Supplementary Fig. S1E–S1G**). In sum, ATM deletion unleashes a mesenchymal, dedifferentiation program in both the P53-proficient KC (5,8) and the aggressive P53-deficient KPC cancers.

**Figure 1.**
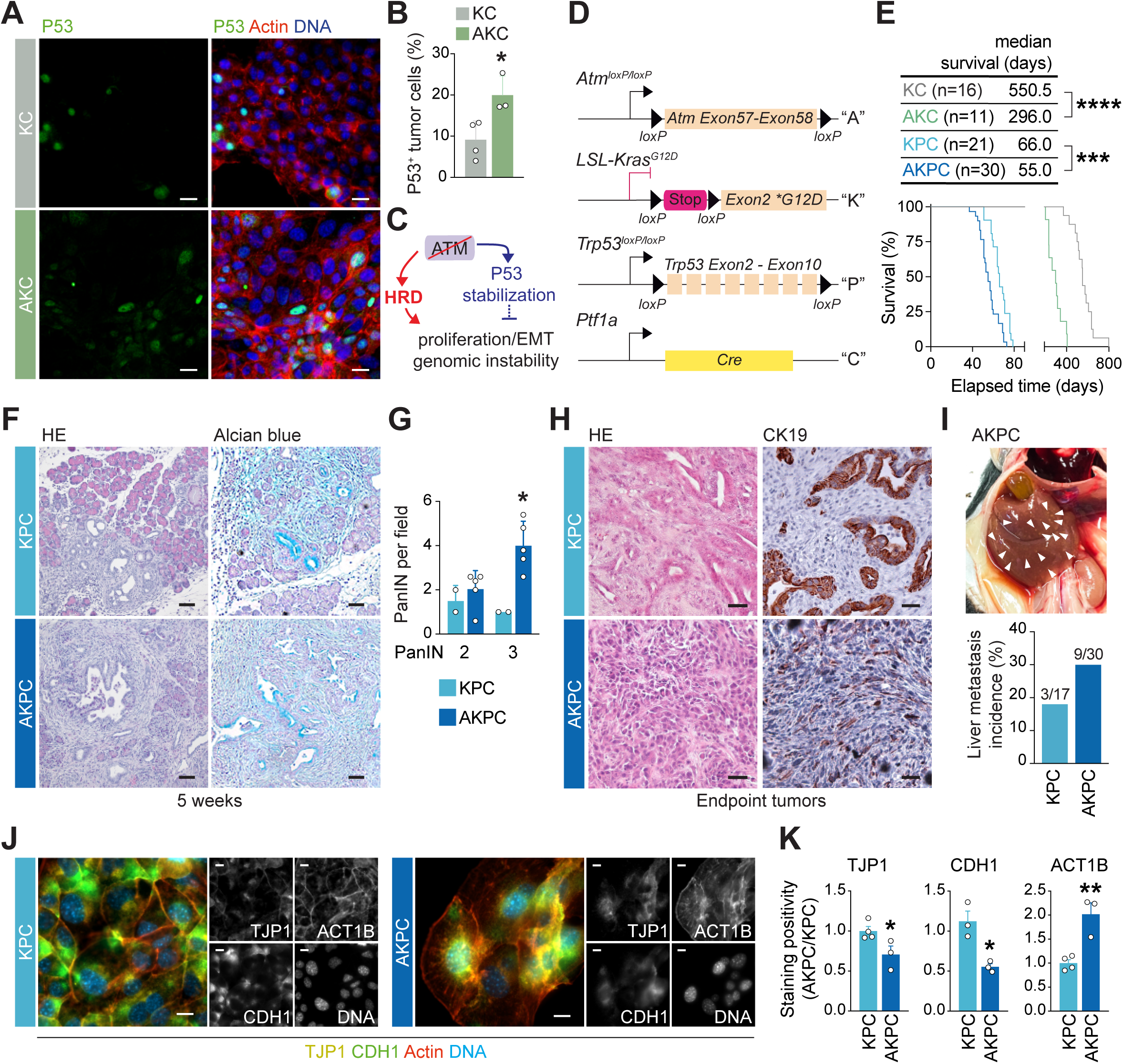
ATM loss-of-function accelerates pancreatic cancer tumorigenesis in P53-deficient context. **A** and **B,** Immunofluorescence staining for P53 (green) and direct fluorescence staining of cortical F-actin by phalloidin-Atto565 (red) (**A**) and quantification of P53^+^ cells in *Atm^+/+^*; *LSL-Kras^G12D/+^*; *Ptf1a^Cre/+^*(KC) and *Atm^fl/fl^*; *LSL-Kras^G12D/+^*; *Ptf1a^Cre/+^*(AKC) tumor cells (**B**). Cells were counterstained with DAPI (blue). Scale bars, 20 µm. Results show means ± SD of ≥ 3 cell lines used in at least 2 independent experiments. Each dot represents a cell line. *, *P* < 0.05, unpaired Student *t* test. **C,** Schematic model depicting the hypothetic impact of P53 on oncogenic effects mediated by ATM loss-of-function in AKC pancreas. **D,** Schematic representation of *Ptf1a^Cre^* (“C”), *LSL-Kras^G12D^* (“K”), floxed *Atm* (“A”), and floxed *Trp53* (“P”) alleles. **E,** Kaplan-Meier analysis of survival of *Atm^+/+^*; *Trp53^fl/fl^*; *LSL-Kras^G12D/+^*; *Ptf1a^Cre/+^* (KPC) (n=21), *Atm^fl/fl^*; *Trp53^fl/fl^*; *LSL-Kras^G12D/+^*; *Ptf1a^Cre/+^*(AKPC) (n=30) (top), *Atm^+/+^*; *LSL-Kras^G12D/+^*; *Ptf1a^Cre/+^* (KC) (n=16), and *Atm^fl/fl^*; *LSL-Kras^G12D/+^*; *Ptf1a^Cre/+^* (AKC) (n=11) mice (bottom). *** *P* < 0.001; *\*\*\*\*, P* < 0.0001, log-rank (Mantel-Cox) test. Median survival of KC, AKC, KPC, and AKPC mice are shown. **F** and **G,** Histologic sections stained by hematoxylin-eosin (HE) and alcian blue (**F**), and quantification of pancreatic intraepithelial neoplasia (PanIN) lesions (**G**) in pancreata resected from 5-week-old KPC (n=2) and AKPC (n=5) mice. Scale bars, 100 µm. Results show means ± SD. Each dot represents a mouse. *, *P* < 0.05, unpaired Student *t* test. **H,** Histologic sections stained by HE (left) and immunohistochemistry for CK19 (right) on KPC and AKPC endpoint pancreatic tumors. Scale bars, 50 µm. **I,** Representative image of PDAC liver metastases in an endpoint AKPC animal (top) and incidence of liver metastasis in endpoint AKPC and KPC mice (bottom). White arrows show liver metastasis. Fisher’s exact test, not significant. **J** and **K,** Immunofluorescence staining for the TJP1 (yellow) and CDH1 (green) epithelial markers and direct fluorescence staining of cortical F-actin by phalloidin-Atto565 (red) (**J**) and quantification of TJP1 (left), CDH1 (center), and F-actin^+^ (right) surface, normalized to nucleus count, in KPC and AKPC malignant cells (**K**). Cells were counterstained with DAPI (blue). Scale bars, 10 µm. Results show means ± SD ≥ 3 cell lines, each used in at least 2 independent experiments. Each dot represents a cell line. *, *P* < 0.05; **, *P* < 0.01, unpaired Student *t* test. EMT, epithelial-mesenchymal transition; HRD, homologous recombination deficiency.

### ATM-deficient tumor cells generate a myofibroblast-enriched tumor microenvironment

To further explore ATM loss-specific tumor biology, we extended our histopathological characterization to the stromal compartment. Immunostainings revealed modest changes in the immune landscape of ATM-deficient tumors, including reduced B220^+^ and B220^+^CD11b^+^ populations but preserved CD11b^+^, Ly6C^+^, and CD11b^+^Ly6C^+^ myeloid and monocytic compartments, indicating a selective loss of B-lineage cells and biphenotypic progenitors (**Supplementary Fig. S2A** and **S2B**). Additionally, a downward trend in CD3^+^ and CD8^+^ T cells and a marked reduction in M2-like CD206^+^ tumor-associated macrophages (TAMs) was observed, suggesting an immunologically cold ecosystem (**Supplementary Fig. S2A** and **S2B**). In contrast, picrosirius red staining performed to visualize collagen deposition revealed pronounced fibrosis upon ATM depletion occurring in both P53-depleted and non-depleted contexts, which may contribute to the observed restricted immune infiltration (**Fig. 2A** and **B; Supplementary Fig. S2C**). The significant collagen deposition comprised an increased collagen type I/III ratio (**Supplementary Fig. S2D** and **S2E**), ultimately denoting a possible enrichment in collagen-producing αSMA^+^ myCAFs. The AKPC collagen fibers appeared also more organized with increased directionality (**Supplementary Fig. S2D** and **S2E**), a negative prognostic feature promoting epithelial-to-mesenchymal transition (EMT) and dissemination (31,32), in line with the invasive phenotype of AKPC tumors (**Fig. 1H-K**). To examine the dominating CAF subtype in this collagen-enriched stroma, we analyzed cellular αSMA immunoreactivity and reported a significant increase of αSMA^+^ fibroblast subtype in AKPC compared to KPC PDAC counterparts (**Fig. 2A** and **B; Supplementary Fig. S2F**). Concomitantly, SMAD2 phosphorylation and nuclear translocation, mirroring TGF-β canonical axis activation, was observed in αSMA^+^ myofibroblasts supporting an ATM loss-dependent TME remodeling toward a myCAF fate (26) through cancer cell-derived factors. Again, this myofibroblastic-specific programming was P53-independent, highlighting the epithelial ATM status as a critical determinant of CAF differentiation (**Fig. 2A** and **B; Supplementary Fig. S2A**). To fully capture the intratumoral stromal landscape of ATM-deficient PDAC tumors, we next conducted a single-nucleus multiomics sequencing on KPC and AKPC tumors, combining gene expression and chromatin profiling (**Fig. 2C**). A total of 10,445 nuclei were sequenced (AKPC: 5,698, KPC: 4,747). After initial quality controls, we visualized the results using dimensionality reduction through uniform manifold approximation and projection and identified 13 cell clusters, annotated based on signature genes cross-referenced with known markers of specific cell populations (17,33) (**Fig. 2D; Supplementary Fig. S3A–S3F**). All cell types were present in both KPC and AKPC tumors but their relative proportions differed significantly (**Fig. 2E; Supplementary Fig. S3E; Supplementary Table S1**). Transcriptional signatures associated with hypoxia and tissue dissociation (34) remained low and did not differ between genotypes, indicating that tissue processing did not introduce stress-related artifacts (**Supplementary Fig. S3G**). We observed increased malignant ductal and EMT-like cell contents in ATM-deficient tumors, along with differences in stroma cellular composition (**Fig. 2E**). AKPC tumors exhibited a myeloid-enriched profile, dominated by an accumulation of M1-like TAMs, while KPC tumors showed higher proportions of B and T lymphocytes, reflecting distinct immune compositions shaped by ATM status (**Supplementary Fig. S3H** and **S3I**). Consistent with our histological examinations (**Fig. 2A** and **B**), AKPC tumors exhibited increased total number of fibroblasts with a particular enrichment in αSMA^+^ myCAFs, highly expressing collagen and tenascin ECM genes (**Fig. 2E** and **F; Supplementary Fig. S4A and S4B**). In contrast, ATM-proficient KPC tumors displayed increased iCAF and apCAF numbers (**Fig. 2E**). Flow cytometry analysis of CAFs confirmed a significantly increased ratio of αSMA^+^Ly6C^-^ myCAFs on αSMA^-^Ly6C^+^ iCAFs in AKPC tumors (**Fig. 2G**). Interestingly, fibroblasts and CAFs originating from KPC and AKPC tumors depicted distinct molecular signaling patterns (**Fig. 2H and I; Supplementary Fig. S4C**). Specifically, both AKPC myCAF and iCAF subtypes upregulated genes involved in ECM organization, cell contractility, and TGF-β signaling, usually described as myofibroblastic features (**Fig. 2H** and **I**). At the same time, single-cell trajectory analysis of KPC CAFs revealed distinct transition dynamics among the three segregated CAF subtypes. In contrast, the inferred trajectory for AKPC CAFs exhibited scattered transitions, with a clear overlap between myCAF and iCAF subpopulations (**Supplementary Fig. S4D**). This proximity between iCAF and myCAF states in ATM-deficient tumors, along with enriched myofibroblastic programing, suggested overall enhanced myCAF differentiation dynamics. Finally, no significant differences in chromatin profiles or predicted transcription factor activity were reported between KPC and AKPC CAFs, indicating that the aforementioned programing events were not regulated at an epigenetic level (**Supplementary Fig. S4E**).

**Figure 2.**
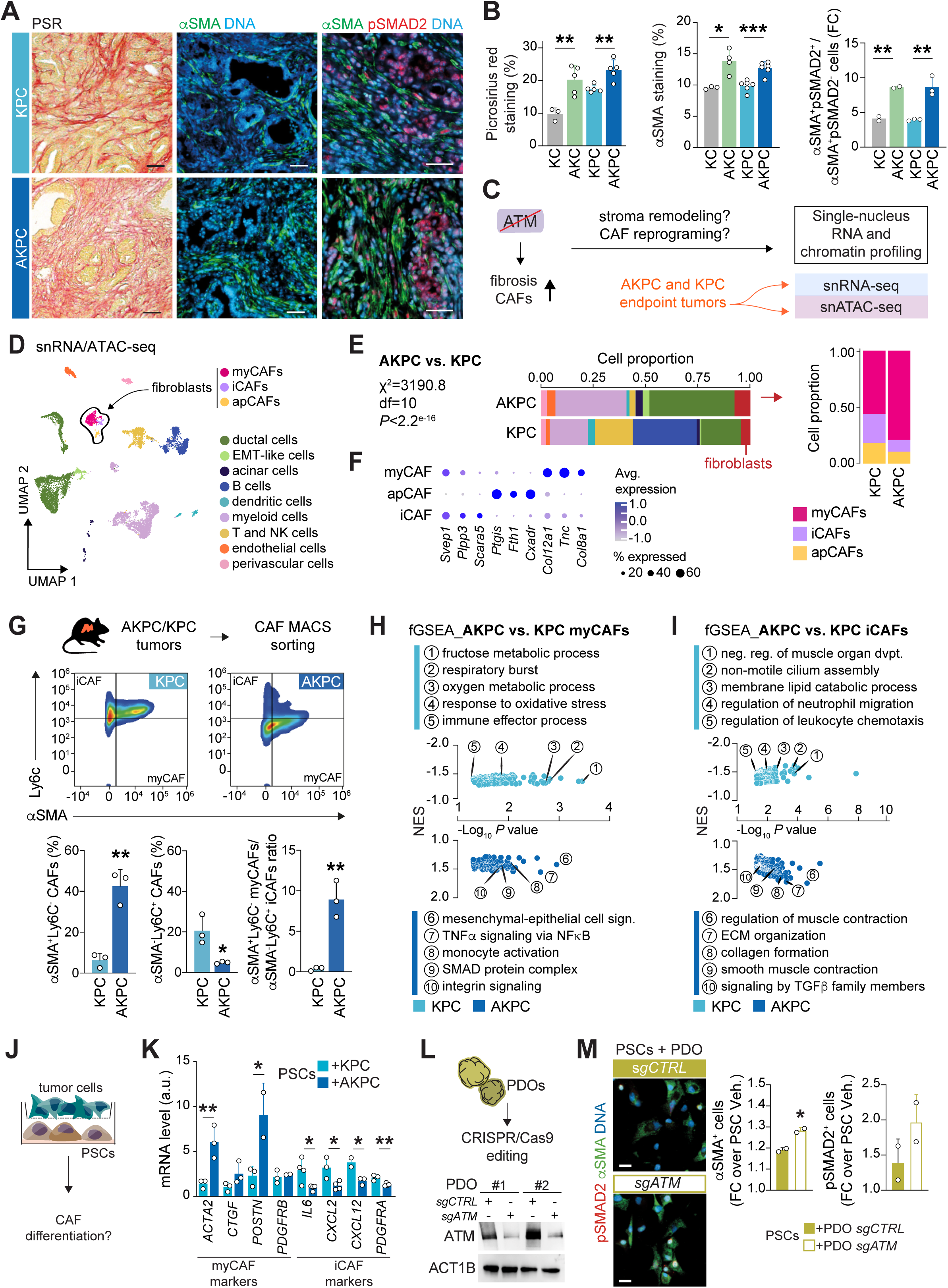
HRD ATM-deleted PDAC exhibits an enriched myCAF stroma. **A,** Representative histologic sections stained by picrosirius red (left) and immunofluorescence staining for αSMA (green) and SMAD2 p-S465/S467 (red) on *Atm^+/+^*; *Trp53^fl/fl^*; *LSL-Kras^G12D/+^*; *Ptf1a^Cre/+^* (KPC) and *Atm^fl/fl^*; *Trp53^fl/fl^*; *LSL-Kras^G12D/+^*; *Ptf1a^Cre/+^* (AKPC) endpoint pancreatic tumors. Cells were counterstained with DAPI (blue). Scale bars, 100 µm. **B,** Quantification of picrosirius red-positive (left), αSMA^+^ surface (center) and αSMA^+^SMAD2 p-S465/S467^+^ cells vs. αSMA^+^SMAD2 p-S465/S467^-^ cells (right) in *Atm^+/+^*; *LSL-Kras^G12D/+^*; *Ptf1a^Cre/+^*(KC), *Atm^fl/fl^*; *LSL-Kras^G12D/+^*; *Ptf1a^Cre/+^* (AKC) (shown in (**Supplementary Fig. S2A**)), KPC, and AKPC (shown in (**A**)) endpoint pancreatic tumors. Results show means ± SD of ≥ 3 KPC and AKPC mice and ≥ 2 KC and AKC mice. Each dot represents a mouse. *, *P* < 0.05; **, *P* < 0.01; ***, *P* < 0.001, unpaired Student *t* test. **C,** Schematic illustration of the single cell-resolved workflow used to simultaneously capture transcriptome and epigenome. KPC (n=2) and AKPC (n=2) tumors were dissociated into single cells and subjected to single nucleus capture using the 10X Genomics platform. **D,** UMAP embedding visualizing integrated snRNA and ATAC-seq clusters of cell populations identified in KPC and AKPC tumors. **E,** Proportion of cells from KPC and AKPC tumors (left). The right panel depicts the proportion of myofibroblastic (myCAFs), inflammatory (iCAFs) and antigen-presenting (apCAFs) cancer-associated fibroblast subpopulations identified from the fibroblast cluster (right). **F,** Dot plot showing the three highest expressed genes segregating the myCAF, iCAF and apCAF subpopulations. Color intensity indicates the average expression for the corresponding gene. Dot size corresponds to the proportion of cells with non-zero expression of the gene in each cell cluster. **G,** Representative flow-cytometric analysis (upper panels) and quantification by flow cytometry (lower panels) of αSMA^+^Ly6C^-^ and αSMA^-^Ly6C^+^ CAFs in KPC and AKPC endpoint pancreatic tumors. Results show means ± SD of 3 KPC and AKPC mice. Each dot represents a mouse. *, *P* < 0.05; **, *P* < 0.01, unpaired Student *t* test. **H** and **I,** GO, REACTOME and Hallmark-based gene-set enrichment analysis of the transcriptomics data showing upregulated and downregulated pathways with *P* ≤ 0.05 in AKPC vs. KPC myCAFs (**H**) and iCAFs (**I**) populations. **J,** Schematic representation of the Transwell-based co-culture assay (**K** and **M**). **K,** qPCR analysis of myCAF and iCAF marker gene expression in PSCs co-cultured for 72 hours with KPC or AKPC tumor cells as in (**J**). Results show means ± SD of ≥ 2 cell lines, each used in at least 2 independent experiments. Each dot represents a cell line. *, *P* < 0.05; **, *P* < 0.01, unpaired Student *t* test. **L,** Western blot analysis of ATM in *sgATM* or *sgCTRL* CRISPR-edited PDAC patient-derived organoids (PDOs). **M,** Immunofluorescence staining for αSMA (green) and SMAD2 p-S465/S467 (red) and quantification of αSMA^+^ and SMAD2 p-S465/S467^+^ PSCs vs. in inactivated PSCs after 72 hours of co-culture with *sgATM* or *sgCTRL* CRISPR-edited PDOs as in (**J**). Results show means ± SD of two PDO lines used in 3 independent experiments. Each dot represents a PDO line. *, *P* < 0.05, unpaired Student *t* test. GO, gene ontology; MACS, magnetic cell separation.

To further substantiate the functional ATM-specific tumor–CAF interplay, we established a transwell-based, cell-cell contact-free co-culture system for tumor cells and pancreatic stellate cells (PSCs) as CAF precursors (35), which allowed tracing the acquisition of distinct myCAF and iCAF traits (**Fig. 2J; Supplementary Fig. S5A–S5D**). Indeed, the ATM-deficient cancer cells (AKC or AKPC) led to αSMA^+^ myCAF differentiation of the PSCs, independently of their P53 genetic status (**Fig. 2J** and **K; Supplementary Fig. S5E** and **S5F**). A conditioned medium-based approach confirmed these observations and further pinpoints ATM-null cancer cell secretomes to comprise signals initiating myCAF reprogramming (**Supplementary Fig. S5G** and **S5H**). Interestingly, the evaluation of CAF marker expression revealed distinct transcriptional programing driving myCAF differentiation, dependent on P53 status (**Fig. 2K; Supplementary Fig. S5E**). In the P53-deficient context, ATM loss markedly diminished the expression of iCAF genes while simultaneously upregulating myCAF markers *ACTA2* and *POSTN*. Conversely, in the presence of P53, ATM deficiency resulted in the upregulation of myCAF genes without significantly affecting iCAF marker expression (**Fig. 2K; Supplementary Fig. S5E**). Immunostainings of endpoint tumors confirmed these findings, showing increased expression of myCAF markers CTGF and POSTN, along with reduced CXCL2 and Ly6C iCAF markers in ATM-deficient tumors (**Supplementary Fig. S5I** and **S5J**).

Next, we employed our recently developed and validated porcine urinary bladder (PUB) model as a decellularized scaffold to track CAF differentiation dynamics in a versatile and flexible co-culture format (**Supplementary Fig. S5K** and **S5L**) (36,37). Indeed, the PUB allowed tumor growth *ex vivo* and accurately recapitulated *in vivo* tumor features with a more spindle-shaped neoplastic epithelium and an activated myCAF enriched surrounding niche (**Supplementary Fig. S5K**) in ATM-deficient grafts. These data licensed the PUB assay for further readouts and confirmed our previous findings in an additional 3D model allowing cell–cell interactions. Finally, both *ATM^+/Δ^* human MIA PaCa-2 cells (4) and CRISPR/Cas9-edited *ATM*-knockout patient-derived organoids (PDOs), employed in the same co-culture settings translated our findings on myCAF programming capacity to human PDAC derived-systems (**Fig. 2L** and **M; Supplementary Fig. S5M** and **S5N; Supplementary Table S3**).

### Global profiling of ATM-deficient tumor cells illuminates an actin cytoskeleton remodeling and secretory program

After confirming that epithelial ATM loss-of-function promotes myCAF differentiation, we investigated the underlying tumor cell molecular programs using our single nucleus-resolved multiomics analysis. EMT-like cells from AKPC tumors showed enhanced transcriptional programs for cell locomotion and invasion compared to KPC counterparts (**Fig. 3A** and **B**). Both malignant ductal and EMT-like cells additionally showed higher expression of mesenchymal genes, corroborating that ATM loss is a driver of the mesenchymal phenotype, in line with the increased EMT-like cells proportion in AKPC tumors (**Fig. 2E** and **Fig. 3A–3C**). Bulk RNA sequencing on AKPC and KPC cell lines validated these findings, revealing significant downregulation of gene sets involved in epithelial cell organization such as adherens junction, gap junction, or cell junction organization (**Fig. 3D** and **E**). Intriguingly, both single-cell and bulk RNA sequencing analyses demonstrated a positive correlation between ATM-deficiency and programming related to mitochondrial and respiratory functions (**Fig. 3B** and **E**). Conversely, ATM-proficient KPC cells exhibited upregulation of gene programs related to metabolism and immune cell regulation (**Fig. 3E**). Consistently, we also inferred distinct gene regulatory networks between KPC and AKPC ductal cells, the ATM-proficient cells exhibiting enhanced predicted *Stat3*, *Stat5*, *Jun*, and *Nfkb2* transcription factor activity, along with increased expression of target genes involved in inflammatory response and interleukin signaling. In contrast, predicted transcription factor signaling network in AKPC ductal cells was more closely associated with cell stress response (**Supplementary Fig. S6A** and **S6B**).

**Figure 3.**
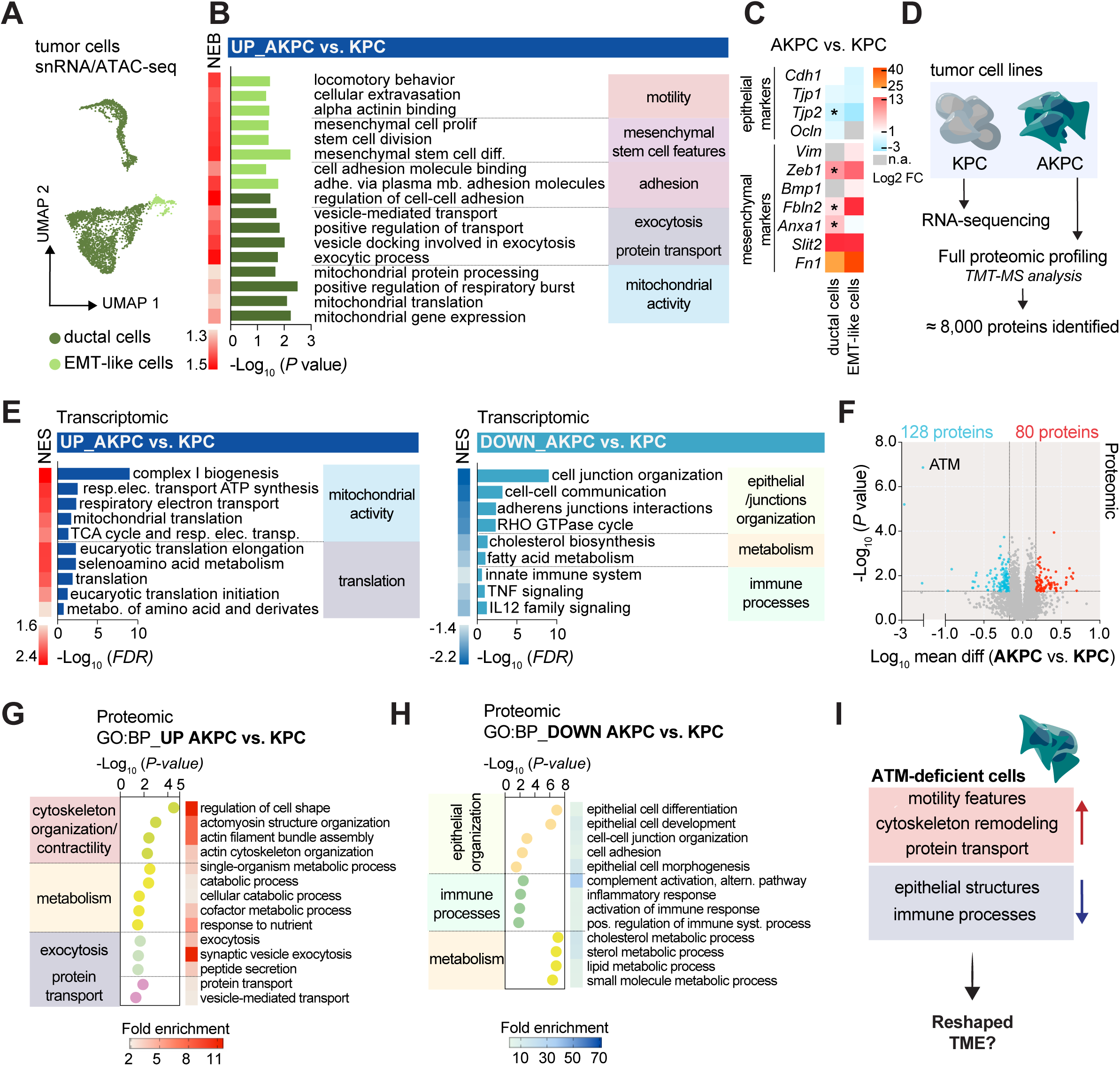
Transcriptomic and proteomic profiling of ATM-deficient tumor cells illuminate an actin cytoskeleton remodeling and secretory program. **A** and **B,** UMAP of snRNA/ATAC-seq data of *Atm^+/+^*; *Trp53^fl/fl^*; *LSL-Kras^G12D/+^*; *Ptf1a^Cre/+^* (KPC) and *Atm^fl/fl^*; *Trp53^fl/fl^*; *LSL-Kras^G12D/+^*; *Ptf1a^Cre/+^* (AKPC) tumors illustrating the EMT-like and ductal cell populations (**A**) and GO, REACTOME and Hallmark-based gene-set enrichment analysis of the transcriptomics data showing upregulated pathways with *P* ≤ 0.05 in AKPC vs. KPC EMT-like (light green) and ductal (dark green) cells (**B**). **C,** Heatmap depicting the expression levels of epithelial (*Cdh1*, *Tjp1*, *Tjp2*, and *Ocln*) and mesenchymal (*Vim*, *Zeb1*, *Bmp1*, *Fbln2*, *Anxa1*, *Slit2*, and *Fn1*) genes in AKPC vs. KPC ductal and EMT-like cells. *, *P* < 0.05. **D,** Schematic representation of the transcriptomics and proteomics approaches conducted on KPC and AKPC tumor cell lines. **E,** REACTOME and KEGG-based gene set enrichment analysis of the transcriptomics data showing downregulated (left panel) and upregulated (right panel) pathways with FDR ≤ 0.05 in AKPC (n=4) vs. KPC (n=5) cell lines. **F,** Volcano plot displaying differential protein abundance between AKPC (n=5) and KPC (n=5) cells. Red dots represent the upregulated proteins and blue dots represent the downregulated proteins with *P* ≤ 0.05 and fold change ≥ 1.5 in AKPC vs. KPC cells. **G** and **H,** Representative GO:BP functional annotations for proteins upregulated (**G**) and downregulated (**H**) with *P* ≤ 0.05 in AKPC vs. KPC cells. **I,** Schematic summary illustrating the main features of KPC and AKPC cells. BP, biological process; GO, gene ontology; KEGG, Kyoto encyclopedia of genes and genomes; TME, tumor microenvironment.

To add another comprehensive layer to our molecular understanding, and test whether these transcriptomic programs indeed translate into effective signaling network, we then performed a full proteomic analysis which revealed 208 differentially regulated proteins according to ATM genetic status (**Fig. 3D** and **F**). Interestingly, proteome profiling was able to distinguish minute changes in cellular organization, metabolism, and signaling pathways in AKPC cells. Corroborating our transcriptomic profiling, ATM loss was positively correlated with an enrichment in actin cytoskeleton organization and actomyosin machinery protein sets, suggesting enhanced contractility capacities (**Fig. 3G** and **H; Supplementary Fig. S6C**).

Additionally, analysis of epithelial (e.g., CDH1) and EMT-related (e.g., ZEB1) proteins, consistent with mRNA levels, confirmed the ATM deficiency-associated mesenchymal profile (**Fig. 3C; Supplementary Fig. S6D**). ATM loss also correlated with an enrichment in protein transport and exocytosis signaling, aligned with the upregulation of genes related to vesicle-mediated transport in ductal cells from AKPC tumors (**Fig. 3B** and **G; Supplementary Fig. S6C**). In contrast, ATM-proficient KPC cells showed protein expression patterns associated with epithelial differentiation and prominent immune tropism (**Fig. 3H; Supplementary Fig. S6C**). Altogether, these data underpinned that ATM loss-of-function in PDAC cells not only drives mesenchymal programing but might also affect their secretory phenotype, in contrast with the epithelial differentiation and immunoregulatory phenotype reported in ATM-proficient cells, suggesting the existence of distinct dialogs with their respective TME (**Fig. 3I**).

### Stromal myCAF programming driven by the ATM-null tumor epithelium is TGF-β dependent

To identify putative factors involved in ATM-specific tumor–stroma interplay, e.g., cytokines, low-abundant and small secreted factors hardly detectable by shotgun mass spectrometry (38), we analyzed the secretomes of our cancer cell lines using a targeted antibody microarray approach (39) (**Fig. 4A**). AKPC-derived factors profiling revealed a specific enrichment for various inflammatory cytokines and chemokines, integrins, and growth factors linked to actin crosslink formation, actin cytoskeleton, and chemotaxis processes (**Fig. 4B** and **C**).

**Figure 4.**
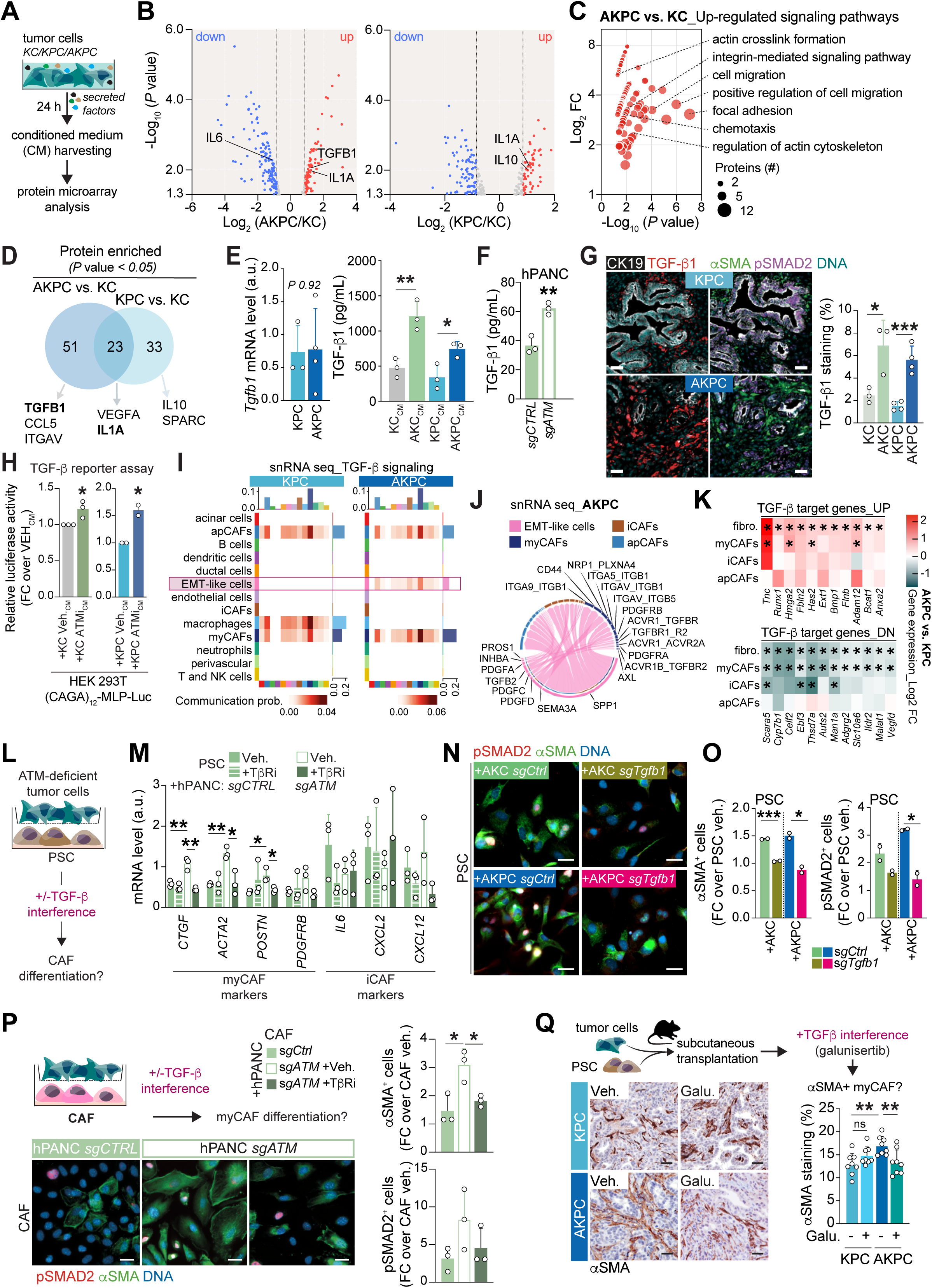
TGF-β1 released by ATM-deficient cancer cell drives myCAF differentiation. **A,** Schematic representation of the secretomics approach conducted on *Atm^+/+^*; *LSL-Kras^G12D/+^*; *Ptf1a^Cre/+^* (KC) (n=3), *Atm^+/+^*; *Trp53^fl/fl^*; *LSL-Kras^G12D/+^*; *Ptf1a^Cre/+^* (KPC) (n=3), and *Atm^fl/fl^*; *Trp53^fl/fl^*; *LSL-Kras^G12D/+^*; *Ptf1a^Cre/+^* (AKPC) (n=2) tumor cell lines (**B–D**). **B,** Volcano plot displaying differential protein abundance in AKPC vs. KC (left) and in KPC vs. KC (right) tumor cell secretomes. Red dots represent the upregulated proteins and blue dots represent the downregulated proteins with *P* ≤ 0.05 and fold change ≥ 1.8 in AKPC (n=2) and KPC (n=3) vs. KC (n=3) cell lines. **C,** KEGG, GO, and BioCarta-based signaling pathway and biological process term enrichment analyses for proteins enriched in AKPC vs. KC cell tumor cell secretomes. **D,** Venn diagram showing the overlap of significantly enriched proteins between AKPC and KPC tumor cell secretomes. **E,** qPCR analysis of *Tgfb1* expression in KPC and AKPC tumor cell lines (left) and ELISA for TGF-β1 on conditioned medium (CM) from KC, AKC, KPC, and AKPC tumor cells (right). Results show means ± SD of ≥ 3 cell lines, each used at least in 2 independent experiments for the qPCR analysis. For the ELISA, at least 2 CM batches per cell line were tested. Each dot represents a cell line. *, *P* < 0.05; **, *P* < 0.01, unpaired Student *t* test. **F,** ELISA for TGF-β1 on CM from *sgATM* and *sgCTRL* CRISPR-edited hPANC tumor cells. At least 2 CM batches per cell line were tested. Each dot represents a cell line. **, *P* < 0.01, unpaired Student *t* test. **G,** Representative histologic sections stained by immunofluorescence staining for CK19 (white), TGF-β1 (red) and αSMA (green) and SMAD2 p-S465/S467 (purple) on KPC and AKPC endpoint pancreatic tumors (left). Cells were counterstained with DAPI (blue). Scale bars, 100 µm. Quantification of TGF-β1^+^ surface (right) KC, (AKC) (shown in (**Supplementary Fig. S7H**)), KPC, and AKPC endpoint pancreatic tumors. Results show means ± SD of 4 KPC and AKPC mice and 3 KC and AKC mice. Each dot represents a mouse. **, *P* < 0.01; ***, *P* < 0.001, unpaired Student *t* test. **H,** Relative luciferase activity of HEK 293T cells transiently transfected with the SMAD-responsive (CAGA)_12_-Luc reporter construct and treated for 48 hours with CM from KC and KPC tumor cells treated or not for 24 h with 1 µM AZD0156 (ATM inhibitor). Results show means ± SD of ≥ 2 cell lines. At least 2 CM batches per cell line were tested. Each dot represents a cell line. *, *P* < 0.05; **, *P* < 0.01, unpaired Student *t* test. **I,** Heatmap representing the inferred communication probability via TGF-β signaling between the different cell clusters in KPC (left) and AKPC (right) tumors. **J**, Chord diagram showing the inferred ligand-receptor interactions between EMT-like cells and myofibroblastic (myCAF), inflammatory (iCAF) and antigen-presenting (apCAF) cancer-associated fibroblast subpopulations in AKPC tumors. **K,** Heatmap showing mRNA expression of known TGF-β target genes up (top) or downregulated (bottom) in fibroblasts (117) and SMAD3/4 target genes (right) in AKPC vs. KPC fibroblasts, myCAFs, iCAFs and apCAFs. Stars indicate significant difference between AKPC vs. KPC with an adjusted *P* < 0.05. **L,** Schematic representation of the Transwell-based co-culture assay (**M**). **M,** qPCR analysis of myCAF and iCAF marker gene expression in PSCs co-cultured for 72 hours with *sgATM* or *sgCTRL* CRISPR-edited hPANC tumor cells and treated or not with the TGFBR1 inhibitor SB431542 (TβRi; 10 µM), as in (**L**). Results show means ± SD of 3 cell lines, each used in at least 2 independent experiments. Each dot represents a cell line. *, *P* < 0.05; **, *P* < 0.01, unpaired Student *t* test. **N** and **O,** Immunofluorescence staining for αSMA (green) and SMAD2 p-S465/S467 (red) on PSCs co-cultured for 72 hours with *sgTgfb1* or *sgCTRL* CRISPR-edited AKC or AKPC tumor cells (left) (**N**) and quantification of αSMA^+^ (left) and SMAD2 p-S465/S467^+^ (right) PSCs (**O**). Cells were counterstained with DAPI (blue). Scale bars, 20 µm. Results show means ± SD of 2 cell lines, each used in 3 independent experiments. Each dot represents a cell line. *, *P* < 0.05; ***, *P* < 0.001, unpaired Student *t* test. **P,** Schematic representation of the Transwell-based co-culture assay (top), immunofluorescence staining for αSMA (green) and SMAD2 p-S465/S467 (red) on CAF lines co-cultured for 72 hours with *sgAtm* or *sgCTRL* CRISPR-edited hPANC tumor cells (bottom left), and quantification of αSMA^+^ (left) and SMAD2 p-S465/S467^+^ (right) CAFs (right). Cells were counterstained with DAPI (blue). Scale bars, 20 µm. Results show means ± SD of 3 cell lines, each used in 2 independent experiments. Each dot represents a cell line. *, *P* < 0.05; ***, *P* < 0.001, unpaired Student *t* test. **Q,** Schematic representation of the subcutaneous assay (top left), immunohistochemistry staining for αSMA (bottom left) and quantification of αSMA^+^ staining (right) on resected subcutaneous tumors arising either from KPC or AKPC cells co-transplanted with PSCs and treated or not with the TGFBR1 inhibitor galunisertib (galu.; 12.5 mg/kg). Scale bars, 50 µm. Results show means ± SD. Each dot represents a subcutaneous tumor. ns, not significant; **, *P* < 0.01, unpaired Student *t* test. FC, fold change; Galu., galunisertib; GO, gene ontology; KEGG, Kyoto encyclopedia of genes and genomes; Veh., vehicle.

Among the enriched proteins captured from ATM-null secretomes, TGF-β1 emerged as the most promising candidate, particularly considering its role in driving myCAF differentiation (26). However, the specific mutations in cancer cells that lead to increased TGF-β1 release remain largely unknown. In contrast, KPC cell secretomes revealed enrichments in proteins involved in B cell regulation and ECM binding signaling pathways, rather indicative of a KPC-dependent inflammatory phenotype (**Supplementary Fig. S7A**). Surprisingly, the interleukin IL1α, a major iCAF inducer (26), was detected in both AKPC and KPC secretomes suggesting the possibility of spatial coexistence or dynamic plasticity of CAF subtypes within the TME (**Fig. 4B** and **D**). Interestingly, ATM-dependent TGF-β1 regulation operates through post-translational mechanisms as indicated by unchanged mRNA but significantly increased protein release in both ATM-deficient mouse cancer lines and CRISPR-edited *ATM*-KO human patient-derived xenograft (PDX) hPANC lines (**Fig. 4E** and **F; Supplementary Fig. S7B** and **S7C**). Consistently, we detected no significant differences in TGF-β ligands mRNA and protein expression between KPC and AKPC ductal and EMT-like cells (**Supplementary Fig. S7D** and **S7E**). Concomitantly, the myCAF population displayed the highest levels of type I and II TGF-β receptor expression, underscoring their potential role as primary recipient cells in TGF-β signaling independent of the epithelial genotype (**Supplementary Fig. S7E**). Although the gene expression patterns of TGF-β receptors varied among KPC and AKPC malignant cells, these variations did not translate into significant differences in protein abundance (**Supplementary Fig. S7E** and **S7F**). In contrast, the secretomes of ATM-deficient cells, which are enriched with TGF-β1, demonstrated a greater ability to stimulate a TGF-β signaling reporter compared to those of ATM-proficient cells (**Supplementary Fig. S7G**). Consistently, we observed a pronounced TGF-β1 accumulation in the TME of ATM-deficient tumors, spatially overlapping with regions enriched in αSMA^+^ and pSMAD2^+^ signals, indicating localized pathway activation within the stromal compartment (**Fig. 4G; Supplementary Fig. S7H**). Finally, to attribute the increased TGF-β1 secretion to the loss of ATM signaling via its kinase domain, we treated ATM-wildtype KC and KPC cells with a clinical-grade ATM inhibitor (ATMi). ATM kinase inhibition increased the TGF-β signaling activity of conditioned medium from ATM-proficient cells, suggesting a direct role of ATM in regulating TGF-β1 bioavailability (**Fig. 4H**). To further clarify the specific TGF-β-mediated crosstalk between tumor cells and CAFs in ATM-deficient tumors, we employed CellChat analysis (40) to examine cell-cell communication and identified distinct signaling patterns between KPC and AKPC tumors (**Supplementary Fig. S7I**). Notably, TGF-β signaling played a significant role in interactions between EMT-like cells and CAFs in ATM-deficient tumors, whereas it was primarily involved in interactions between macrophage and CAF subpopulations in KPC tumors (**Fig. 4I**). Examination of ligand-receptor pairs between EMT-like cells and CAFs revealed the involvement of several TGF-β superfamily members e.g., *Tgfb2*, *Tgfbr1*, *Acvr1*, along with multiple genes encoding for integrin (e.g., *Itgav*, *Itgb1*), known to regulate TGF-β bioavailability (41), exclusively in AKPC tumors (**Fig. 4J; Supplementary Fig. S7J**). In line, CAF subpopulations present in AKPC tumors exhibited more active TGF-β-responsive gene regulation, indicating a positive feed-forward loop specifically driven by the epithelial absence of ATM (**Fig. 4K; Supplementary Fig. S7K**). Further investigations into genes and proteins encoding TGF-β activators (e.g., integrins, *Thbs1*, *Bmp1*) revealed a trend of upregulation in CAFs and malignant cells in an ATM-deficient context (**Supplementary Fig. S7L** and **S7M).**

To further probe causality between ATM deficiency, TGF-β signaling, and myCAF programming, we treated AKC, AKPC, and CRISPR-edited *ATM*-KO hPANC tumor cell**–**PSC co-cultures with the TGFBR1 kinase inhibitor SB431542 (TβRi, abolishing TGF-β pathway canonical intracellular signaling (42)), thereby clearly demonstrating the direct implication of the TGF-β/SMAD axis (**Fig. 4L** and **M; Supplementary Fig. S7N–S7P**). Conversely, an upregulation of iCAF markers was observed upon TβRi, further illustrating the CAF fate switch (**Supplementary Fig. S7O**). The *ex vivo* PUB assay also confirmed our findings (**Supplementary Fig. S7Q**). Building on our previous findings implicating BMP signaling in ATM-deficiency (5), we examined its potential contribution to CAF reprogramming and possible crosstalk with TGF-β signaling (43). However, analysis of our multiomics dataset revealed no enrichment of a BMP4-responsive transcriptional signature between AKPC and KPC tumors (**Supplementary Fig. S8A** and **S8B**). Although recombinant BMP4 induced a myCAF phenotype in PSCs, the chemical inhibition of BMP signaling in PSC– ATM-deficient tumor cell co-cultures did not significantly suppress myCAF programing, showing only a modest increase in iCAF marker expression (**Supplementary Fig. S8C** and **S8D**). In contrast, co-culture experiments using CRISPR-edited *TGFB1*-KO AKC and AKPC tumor cells further confirmed tumor-derived TGF-β1 as the central driver of myCAF programming in the ATM-null setting (**Fig. 4N** and **O; Supplementary Fig. S8E** and **S8F**).

Noting that CAFs can originate from precursors beyond PSCs (44), we next explored CAF ontogeny in our models. By applying PSC- and non-PSC-derived transcriptional signatures to our multiomics dataset (44), we found an enrichment of the non-PSC-derived signature in myCAF and iCAF clusters, while apCAFs more frequently showed a PSC-derived profile, irrespective of tumor genotype (**Supplementary Fig. S8G**). To assess whether our findings extend beyond PSC-derived models, we conducted co-culture experiments with CAF lines isolated from two independent mouse models (*Kras^G12D^*^/+^; *Trp53^lox/+^*; *Tgfbr2^lox/lox^*and *Kras^G12D^*^/+^; *Tgfbr2^lox/lox^*). While ATM-deficient tumor cells similarly induced myCAF reprograming, marker expression patterns differed between PSCs and primary CAFs (**Fig. 4P; Supplementary Fig. S8H–S8J**). These results suggest that myCAF differentiation is not restricted by precursor origin, thought the specific CAF source may shape transcriptional profiles.

Finally, we conducted an *in vivo* subcutaneous experiment employing co-culture of tumor cells and PSCs, treated with galunisertib, a clinical grade TGFBR1 kinase inhibitor (45,46) (**Fig. 4Q**). Corroborating our previous findings, ATM-deficient tumors exhibited a marked enrichment in αSMA^+^ myCAFs. Strikingly, TGF-β signaling interference showed no effect on αSMA levels in ATM-proficient KPC tumors whereas AKPC specimen showed significantly impoverished αSMA^+^ stroma (**Fig. 4Q**). Additionally, the KRAS^G12D^ inhibitor MRTX1133, known to enhance myCAF accumulation (47), indeed significantly increased αSMA^+^ fibroblasts in both AKPC and KPC tumors (**Fig. 4Q; Supplementary Fig. S8K**). In contrast, TGF-β axis interference reshaped myCAFs only in ATM-deficient tumors, highlighting the strong connection between epithelial genotype and TME reprogramming, even in the absence of KRAS signaling as the primary oncogenic driver in PDAC (**Supplementary Fig. S8K**).

Altogether, these data highlighted that ATM-deficiency drives a unique TGF-β-mediated signaling axis between malignant cells and CAFs, primarily through post-translation regulation of TGF-β activity, thereby promoting a myCAF-enriched microenvironment distinct from the inflammatory phenotype observed in ATM-proficient tumors.

### A deregulated actomyosin network in ATM-deficient PDAC cells supports TGF-**β**-induced myCAF differentiation

TGF-β bioavailability has been described to be dependent of integrin expression patterns intimately linked to actin cytoskeleton dynamics and contractile force (48). In line, with our *in vivo*, *in vitro*, transcriptome and proteome data indicating more adhesive and contractile properties upon ATM deletion (**Fig. 1H–1K** and **3**), ACTN4, a critical actin crosslinker of the actomyosin machinery, was upregulated in the ATM-null background (**Fig. 5A–5C; Supplementary Fig. S9A** and **S9B**). In addition, we observed a substantial upregulation of PXN and VINC, proteins involved in focal adhesion dynamics, cytoskeleton rearrangement, and cell contractility regulation (**Fig. 5C; Supplementary Fig. S9B**). CRISPR-edited *ATM* knockout hPANC cells further recapitulated this contractile phenotype in a human PDAC model (**Fig. 5D**). To further characterize the contractile phenotype of ATM-deficient cells, we monitored AKPC ability to progress through confined spaces designed for assessing single-cell contractility-dependent migration, also predictive of metastatic potential (49). Following our previous results, ATM-depleted AKPC cells showed a superior ability to accomplish single-cell migration through microchannels, attesting high cellular plasticity and contractile features (**Fig. 5E**). Interestingly, ATM-deficient AKPC cells were also characterized by an enrichment in mitochondria activity and oxidative phosphorylation gene sets, denoting a possible dysregulation in reactive oxygen species (ROS) production (**Fig. 3B** and **E; Supplementary Fig. S9C**). In line, AKPC, AKC, and ATM knockout hPANC cells exhibited higher ROS levels than ATM-proficient counterparts (**Fig. 5F; Supplementary Fig. S9D**), a phenotype mirrored by ATM inhibition which also induced ROS accumulation and a contractile profile (**Supplementary Fig. S9E–S9G**). ROS production is particularly accentuated by various oncogenic stresses as the accumulation of DNA alterations (50,51), and has also been proposed to affect cellular contractility (52). To test whether ROS drive the contractile and secretory phenotype of ATM-deficient PDAC cells and promote myCAF programming, we used a loss-of-function approach targeting ROS production and contractility regulators (**Fig. 5G**). CRISPR/Cas9-mediated knockout of *Keap1*, the main intracellular inhibitor of NRF2 (53), enhanced antioxidant activity and markedly reduced ROS levels in AKPC and AKC tumor cells (**Fig. 5G** and **H**). *Keap1*-deficient cells notably displayed improved epithelial organization, reduced ACTN4 and PXN expression, and decreased motility, closely resembling the phenotype observed in PXN- or ACTN4-deficient AKC and AKPC cells (**Fig. 5I** and **J**). Additionally, ATM-deficient cells were subjected to mitochondrial ROS inhibition (ROSi) using the small-molecule inhibitor YCG063, further validating the identified ROS-contractility axis (**Supplementary Fig. S9H–S9L**). Vice versa, ROS production induction, mediated by the oxidant menadione (2-methyl-1,4-naphthoquinone (54)), increased ACTN4 and PXN expression in AKPC cells, overall demonstrating a conceptual link between ROS overproduction and tumor cell contractility (**Supplementary Fig. S9M** and **S9N**). Finally, we investigated whether the ROS-mediated contractility of tumor cells specific to ATM loss might regulate protein secretion and influence subsequent CAF differentiation. To test this, we measured TGF-β1 release in CRISPR-edited *Keap1*, *Pxn*, and *Actn4* knockout AKC and AKPC tumor cells. While ACTN4 or PXN loss caused only a modest non-significant decrease in TGF-β1 levels, suggesting that cytoskeletal disruption alone only partially impacts the secretory phenotype, *Keap1* knockout considerably reduced TGF-β1 secretion (**Fig. 5K**). These findings place ROS upstream of both contractility and TGF-β1 bioavailability regulation in ATM-deficient PDAC cells. Consistently, both co-culture and conditioned medium experiments showed impaired activation of canonical TGF-β signaling and reduced PSC differentiation into αSMA^+^ myCAFs (**Fig. 5L** and **M; Supplementary Fig. S9O–S9S**).

**Figure 5.**
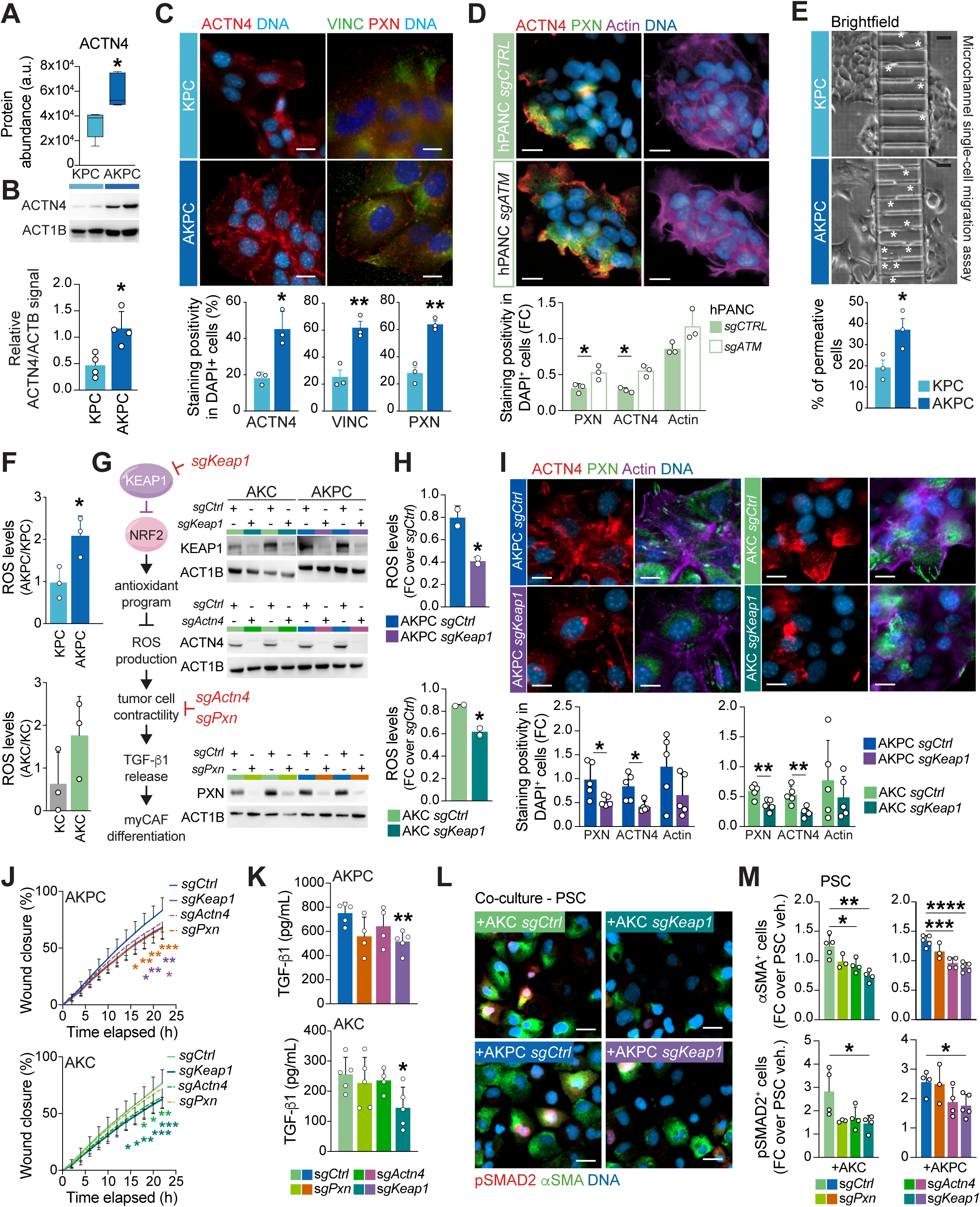
A deregulated actomyosin network in ATM-deficient PDAC cells supports TGF-β-induced myCAF differentiation. **A,** Protein abundancies for ACTN4 in *Atm^+/+^*; *Trp53^fl/fl^*; *LSL-Kras^G12D/+^*; *Ptf1a^Cre/+^* (KPC, n=5) and *Atm^fl/fl^*; *Trp53^fl/fl^*; *LSL-Kras^G12D/+^*; *Ptf1a^Cre/+^* (AKPC, n=5) cells. Results are represented as Tukey box plots of 5 biological replicates. *, *P* < 0.05, unpaired Student *t* test. **B,** Western blot analysis (top) and quantification (bottom) of ACTN4 in KPC and AKPC tumor cells. Results show means ± SD of 2 cell lines each used in four independent experiments. Each dot represents an independent experiment. *, *P* < 0.05, unpaired Student *t* test. **C,** Immunofluorescence staining for ACTN4 (red, left), VINC (green) and PXN (red, right) on KPC and AKPC tumor cells (top) and quantification of ACTN4^+^, VINC^+^, and PXN^+^ staining in DAPI^+^ cells (bottom). Cells were counterstained with DAPI (blue). Scale bars, 20 µm. Results show means ± SD of 3 independent experiments. At least 2 cell lines per genotype were tested. Each dot represents an independent experiment. *, *P* < 0.05; **, *P* < 0.01, unpaired Student *t* test. **D,** Immunofluorescence staining for ACTN4 (red) and PXN (green), and direct fluorescence staining of cortical F-actin by phalloidin-Atto565 (purple) on *sgATM* and *sgCTRL* CRISPR-edited hPANC tumor cells (top) and quantification of ACTN4^+^, PXN^+^, and F-actin^+^ staining in DAPI^+^ cells (bottom). Cells were counterstained with DAPI (blue). Scale bars, 20 µm. Results show means ± SD of 3 cell lines, each used in 2 independent experiments. Each dot represents a cell line. *, *P* < 0.05, unpaired Student *t* test. **E,** Representative images of the microchannel single cell migration assay conducted with KPC and AKPC tumor cells (top) and proportions of permeative cells (bottom). White asterisks show migrating tumor cells. Scale bars, 50 µm. Results show means ± SD of 4 independent experiments. 3 cell lines/genotype were tested. Each dot represents a cell line. *, *P* < 0.05, unpaired Student *t* test. **F,** Levels of reactive oxygen species (ROS) in KPC and AKPC (top), KC, and AKC (bottom) tumor cells. Results show means ± SD of 3 cell lines used in at least 2 independent experiments. Each dot represents a cell line. *, *P* < 0.05, unpaired Student *t* test. **G,** Schematic representation of the hypothetic ROS-mediated cascade leading to myCAF differentiation in ATM-deficient tumors with, highlighted in red, the genes knocked out by a CRISPR/Cas9-based approach for mechanistical investigation (left) and western blot analyses (right) of KEAP1, ACTN4, and PXN in *sgKeap1*, *sgActn4*, *sgPxn*, or *sgCtrl* CRISPR-edited AKC and AKPC tumor cells, respectively. **H,** Levels of ROS in *sgKeap1* and *sgCtrl* CRISPR-edited AKPC (top) and AKC (bottom) tumor cells. Results show means ± SD of 2 cell lines used in at least 3 independent experiments. Each dot represents a cell line. *, *P* < 0.05, unpaired Student *t* test. **I,** Immunofluorescence staining for ACTN4 (red) and PXN (green), and direct fluorescence staining of cortical F-actin by phalloidin-Atto565 (purple) on *sgKeap1* and *sgCtrl* CRISPR-edited AKPC and AKC tumor cells (top) and quantification of PXN^+^, ACTN4^+^, and F-actin^+^ staining in DAPI^+^ cells (bottom). Cells were counterstained with DAPI (blue). Scale bars, 20 µm. Results show means ± SD of 5 independent experiments. Two cell lines per genotype were tested. Each dot represents an independent experiment. *, *P* < 0.05; **, *P* < 0.01, unpaired Student *t* test. **J,** Incucyte-based wound healing assay on *sgKeap1*, *sgActn4*, *sgPxn*, or *sgCtrl* CRISPR-edited AKC and AKPC tumor cells. Results show means ± SD of two cell lines per genotype used in in at least 3 independent experiments. *, *P* < 0.05; **, *P* < 0.01; ***, *P* < 0.001, 2way ANOVA. **K,** ELISA for TGF-β1 on CM from *sgKeap1*, *sgActn4*, *sgPxn*, or *sgCtrl* CRISPR-edited AKC and AKPC tumor cells. At least 2 CM batches per cell line were tested. Each dot represents a cell line. *, *P* < 0.05; **, *P* < 0.01, unpaired Student *t* test. **L** and **M,** Immunofluorescence staining for αSMA (green) and SMAD2 p-S465/S467 (red) on PSCs co-cultured for 72 hours with *sgKeap1* and *sgCtrl* CRISPR-edited AKC and AKPC tumor cells (**L**) and quantification of αSMA^+^ and SMAD2 p-S465/S467^+^ PSCs vs. in inactivated PSCs (**M**). Cells were counterstained with DAPI (blue). Scale bars, 20 µm. Results show means ± SD of at least 3 independent experiments where two cell lines per genotype were tested. Each dot represents an independent experiment. *, *P* < 0.05; **, *P* < 0.01; ***, *P* < 0.001; ****, *P* < 0.0001, unpaired Student *t* test. FC, fold change; Veh., vehicle.

Thus, we suggest that ATM deficiency causes genomic instability and oxidative stress subsequently increasing actomyosin contractility in turn affecting cell secretory phenotype and promoting TGF-β release to fashion a myCAF-enriched TME.

### The myCAF-enriched TME in ATM-deficient PDAC drives chemoresistance and invasiveness

HR deficiency attributes a particular vulnerability to platinum-based regimens such as FOLFIRINOX (4,9,28) while a relevant proportion of HRD cancers do not respond albeit their genetic set-up predicts otherwise (27,55,56). To test whether niche reprogramming in ATM-deficient PDAC mediates chemoresistance, we treated respective cancer genotypes with oxaliplatin in our transwell-based co-culture system. While oxaliplatin was efficient on all tumor cell genotypes, the presence of PSCs mediated drug resistance only in ATM-depleted cells, underlining the tumor-promoting function of the TGF-β-driven myCAF fate reprogramming (**Fig. 6A** and **B; Supplementary Fig. S10A** and **S10B**). To challenge TGF-β interference together with FOLFIRINOX as a potential interventional route for dual targeting of tumor epithelium and stroma, we used the PUB system as an *ex vivo* drug testing platform (**Fig. 6C**). As expected, due to their HRDness (4), only ATM-null tumor cells displayed significant responsiveness to the platinum-based FOLFIRINOX regimen. Furthermore, inhibiting TGF-β1 signaling boosted FOLFIRINOX cytotoxic effects only in the condition employing ATM-depleted cells. Of note, the sole TGF-β signaling inhibition already caused relevant cytotoxicity on ATM-deficient cancer cells, implicating a possible cell-autonomous dependence to this pleiotropic cytokine **(Fig. 6C** and **D and Supplementary Fig. S10C** and **S10D**). Stromal targeting worsened outcome in mice due to the formation of dedifferentiated, aggressive PDACs and neither improved survival in genetically unselected human PDAC (19,57). Similarly, treating AKC tumors with FOLFIRINOX or other DDR-interfering agents selected highly invasive clones (9,10). To exclude that similar clonal selection occurs upon combined FOLFIRINOX plus TGF-β inhibitor treatment, we again employed the PUB platform (36) (**Supplementary Fig. S10E**). In accordance with the metastatic burden in AKC and AKPC mice (**Fig. 1I**), ATM-deficient tumor cells exhibited the greatest ability to infiltrate the ECM of the PUB. Nevertheless, their invasive properties were significantly diminished by the abrogation of TGF-β1 signaling (**Supplementary Fig. S10F–S10I**) licensing such strategy for a subsequent preclinical trial in syngeneic mice (**Fig. 6**). Interestingly, TGF-β1 ablation reduced the motility of AKC and AKPC cells (**Supplementary Fig. S10J**) but did not increase their sensitivity to FOLFIRINOX components (**Supplementary Fig. S10K–S10M**). This suggests that while tumor-derived TGF-β1 directly regulates cancer cell migration, its contribution to chemoresistance is indirect and mediated through the ATM deficiency-induced TGF-β-dependent myCAF programming, rather than by cell-autonomous mechanisms.

**Figure 6.**
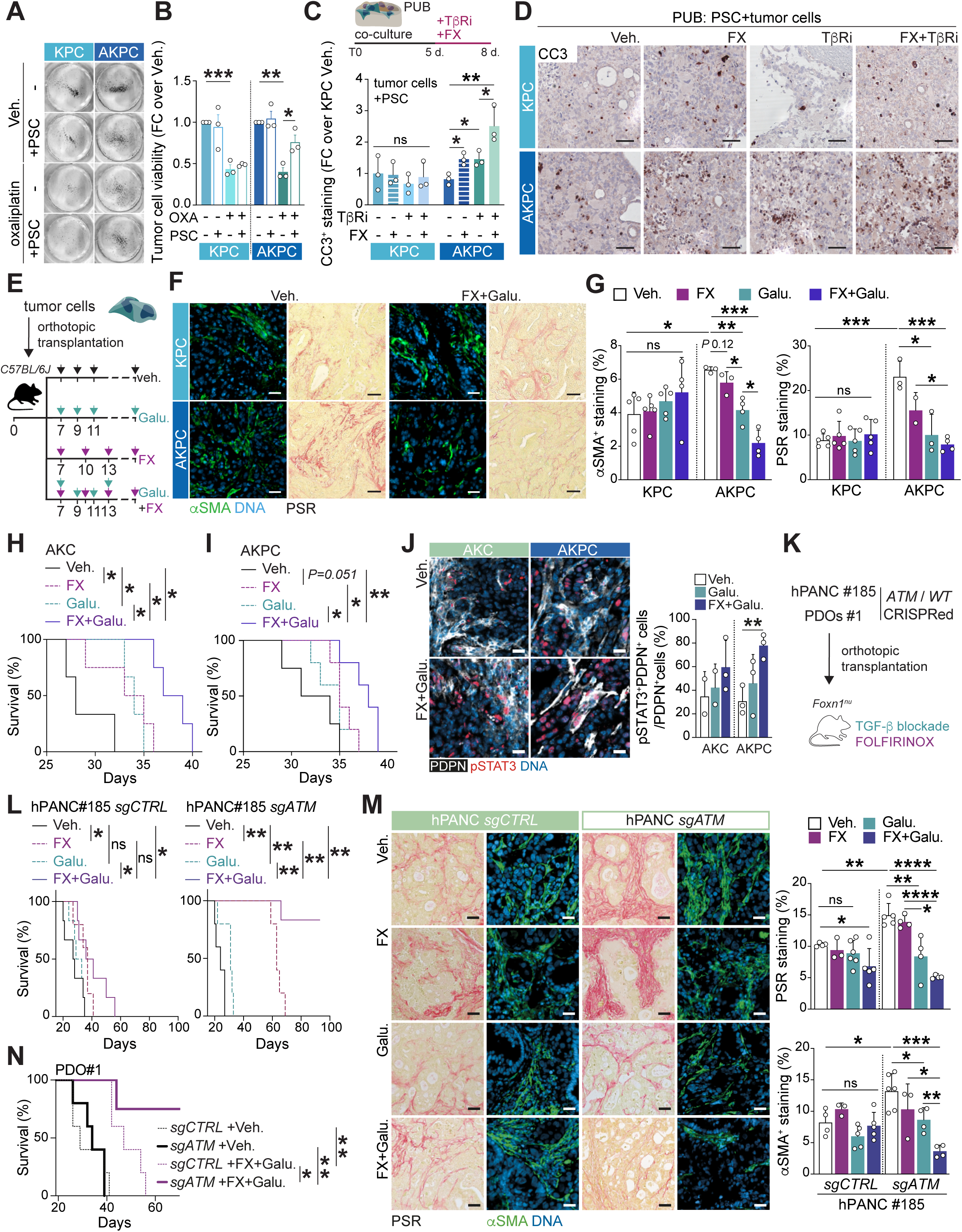
Perturbing tumor–CAF dialog through TGF-β signaling interference improves FOLFIRINOX therapeutic efficacy in ATM-deficient PDAC. **A** and **B,** Colony formation assay on *Atm^+/+^*; *Trp53^fl/fl^*; *LSL-Kras^G12D/+^*; *Ptf1a^Cre/+^* (KPC) and *Atm^fl/fl^*; *Trp53^fl/fl^*; *LSL-Kras^G12D/+^*; *Ptf1a^Cre/+^* (AKPC) tumor cells co-cultured or not with pancreatic stellate cells (PSCs) in a Transwell system and treated or not with 6 µM oxaliplatin (**A**) and quantification of tumor cell viability (**B**). Results show means ± SD of 3 independent experiments. At least 2 cell lines/genotype were tested. Each dot represents an independent experiment. *, *P* < 0.05; **, *P* < 0.01; ***, *P* < 0.001, unpaired Student *t* test. **C** and **D,** Schematic representation of the porcine urinary bladder-based *ex vivo* co-culture assay (**C** top), quantification of cleaved caspase-3^+^ (CC3) surface (**C** bottom), and immunohistochemistry staining for CC3 on resected co-culture grafts involving PSCs and KPC or AKPC tumor cells for 8 days (**D**) and treated or not with the TGFBR1 inhibitor SB431542 (TβRi; 10 µM) and FOLFIRINOX (FX; 75 nM 5-fluorouracil, 25 nM irinotecan, and 5 nM oxaliplatin) as in (**C** top). Scale bars, 100 µm. Results show means ± SD of 3 independent experiments. At least 2 cell lines/genotype were tested. Each dot represents an independent experiment. *, *P* < 0.05; **, *P* < 0.01; ns, not significant, unpaired Student *t* test. **E,** Schematic representation of the orthotopic assay shown in (**F–J**) with treatment administration schedule. **F,** Immunofluorescence staining for αSMA (green) and histologic sections stained by picrosirius red on resected orthotopic tumors arising either from KPC or AKPC cells treated or not with the TGFBR1 inhibitor galunisertib (galu.; 12.5 mg/kg) and FX as in (**E**). Cells were counterstained with DAPI (blue). Scale bars, 100 µm. **G,** Quantification of αSMA^+^ and picrosirius red-positive surface in resected orthotopic tumors of the assay shown in (**E**). Results show means ± SD. Each dot represents a mouse. *, *P* < 0.05; **, *P* < 0.01; ***, *P* < 0.001, unpaired Student *t* test. **H** and **I,** Kaplan-Meier analysis of survival of C57BL/6J mice orthotopically transplanted with *Atm^fl/fl^*; *LSL-Kras^G12D/+^*; *Ptf1a^Cre/+^* (AKC) (**H**) and AKPC (**I**) tumor cells and treated or not with galu. and FX (50.0 mg/kg folinic acid, 25.0 mg/kg 5-fluorouracil, 25.0 mg/kg irinotecan, and 2.5 mg/kg oxaliplatin) as in (**E**). *\*, P* < 0.05; *\*\*, P* < 0.01, log-rank (Mantel-Cox) test. **J,** Immunofluorescence staining for PDPN (white) and STAT3 p-Tyr705 (red) (left) and quantification of PDPN^+^pSTAT3^+^ cells (right) in resected orthotopic tumors of the assay shown in (**E**). Cells were counterstained with DAPI (blue). Scale bars, 50 µm. Results show means ± SD of ≥ 2 AKC and AKPC mice. Each dot represents a mouse. **, *P* < 0.01, unpaired Student *t* test. **K,** Schematic representation of the orthotopic assay shown in (**L** and **M**). **L,** Kaplan-Meier analysis of survival of Nude-*Foxn1^nu^* mice orthotopically transplanted with hPANC #185 *sgCTRL* (left) and hPANC *sgATM* (right) treated or not with galu. and FX (50.0 mg/kg folinic acid, 25.0 mg/kg 5-fluorouracil, 25.0 mg/kg irinotecan, and 2.5 mg/kg oxaliplatin) as in (**K**). *\**, *P* < 0.05; *\*\**, *P* < 0.01, log-rank (Mantel-Cox) test. **M,** Histologic sections stained by picrosirius red and immunofluorescence staining for αSMA (green) on resected tumors of the orthotopic assay shown in (**L**). Cells were counterstained with DAPI (blue). Scale bars, 100 µm. Results show means ± SD. Each dot represents a mouse. *, *P* < 0.05; **, *P* < 0.01; ***, *P* < 0.001; ****, *P* < 0.0001, unpaired Student *t* test. ns, not significant; Veh. Vehicle. **N,** Kaplan-Meier analysis of survival of Nude-*Foxn1^nu^* mice orthotopically transplanted with PDO #1 *sgCTRL* and PDO #1 *sgATM* treated or not with galu. plus FX (50.0 mg/kg folinic acid, 25.0 mg/kg 5-fluorouracil, 25.0 mg/kg irinotecan, and 2.5 mg/kg oxaliplatin). *\**, *P* < 0.05; *\*\**, *P* < 0.01, log-rank (Mantel-Cox) test.

Upon orthotopic transplantation, AKC and AKPC tumor cells indeed formed PDACs with dense fibrotic microenvironment enriched in αSMA^+^ myCAFs, a phenotype reversed upon blockade of TGF-β signaling with galunisertib (**Fig. 6E–6G; Supplementary Fig. S11A–S11C**). FOLFIRINOX and galunisertib alone operated genotype-specific and exclusively prolonged survival of ATM-null tumor-bearing mice to a similar extent (**Fig. 6H** and **I; Supplementary Fig. S11D** and **S11E**). Remarkably, the combination of FOLFIRINOX and galunisertib, also associated with a significant reduced fibrotic phenotype (**Fig. 6F** and **G; Supplementary Fig. S11A–S11C**), caused an additional preclinical benefit as mirrored by the improved median overall survival of AKC and AKPC groups. Magnetic resonance imaging dynamics over time reflected these findings (**Supplementary Fig. S11F**). Similarly, proliferation was decreased while apoptosis increased upon treatment with the combination therapy exclusively in the ATM-null genotypes (**Supplementary Fig. S11G–S11J**). Of note, therapeutic benefit upon treatment was comparatively lower in in the absence of P53 (**Fig. 6H** and **I**). Given previous evidence that myCAF targeting can be counteracted by compensatory iCAF expansion (26), we examined CAF plasticity following TGF-β signaling inhibition. While galunisertib had minimal effect on PDPN^+^Ly6C^+^ iCAFs in KC and KPC tumors, ATM-deficient AKC and AKPC tumors exhibited significant iCAF accumulation, characterized by increased PDPN^+^pSTAT3^+^ cells and a decreased αSMA^+^ myCAFs/Ly6C^+^ iCAFs ratio (**Fig. 6J; Supplementary Fig. S11K** and **S11L**). These findings suggest a CAF phenotypic shift that may undermine therapeutic efficacy through iCAF-driven pro-tumorigenic signaling.

To evaluate the translational impact of our findings, we employed two human-derived preclinical models (orthotopic transplantation of hPANC or CRISPR-edited *ATM*-KO PDOs) and demonstrated genotype-specific efficacy of combining TGF-β pathway interference with FOLFIRINOX in ATM-deficient PDAC (**Fig. 6K–6M**). Notably, therapeutic effects were more pronounced than in mouse cancer cell-derived models, with five of six ATM-KO hPANC-transplanted mice and three of four ATM-KO PDO-transplanted mice treated with FOLFIRINOX plus galunisertib surviving to the experimental endpoint. This benefit correlated with marked TME remodeling, including reduced myCAF accumulation and fibrosis (**Fig. 6M**).

### ATM-deficient human pancreatic cancers exhibit distinctive modifications within their specific tumor microenvironments

To assess whether ATM-specific stromal alterations are also present in human PDAC, we performed transcriptomic deconvolution of the TCGA-PAAD bulk RNA-seq data (58) using BayesPrism, a Bayesian framework for inferring cell-type composition from bulk transcriptomes (59) (**Fig. 7A; Supplementary Fig. S12A** and **S12B**). BayesPrism deconvolution was performed using scRNA-seq reference annotated for 11 major cell types from two independent studies (17,60). UMAP embedding demonstrated cell subsets segregation, including malignant, healthy ductal cells, and fibroblast clusters (myCAFs, iCAFs, non-CAFs; **Supplementary Fig. S12A**). Deconvolution accuracy was supported by strong correlation between inferred pseudobulk transcriptomes and references profiles (**Supplementary Fig. S12B**). Given the low incidence of ATM mutant cases, we stratified tumors based on *ATM* expression levels as a surrogate for functional ATM status (**Supplementary Fig. S12C**), a metric previously linked to overall survival in human PDAC (5). Notably, *ATM* expression negatively correlated with the estimated abundance of malignant ductal cells but positively with healthy ductal cells, supporting an association between low *ATM* levels and malignant expansion (**Fig. 7A**). This aligns with our mouse data and underscores the utility of this approach for further dissecting CAF profiles. We next examined the relationship between *ATM* expression in malignant ductal cells and deconvoluted fibroblast subsets (**Fig. 7B**). *ATM* expression in malignant ductal cells was inversely correlated with myCAF abundance and positively associated with iCAF proportions, mirroring the CAF programming seen in ATM-deficient AKC and AKPC models (**Fig. 7B**). These results further support a tumor-intrinsic role for ATM loss in shaping CAF contexture. To extend this framework to additional DDR-related genes, which similarly exhibit low mutations frequency, we again applied expression-based stratification to enable comparative analysis (**Fig. 7C; Supplementary Fig. S12C**). We found that *ATM* expression in malignant ductal cells, along with *RAD51C* and *FANCG*, uniquely exhibited a significant negative correlation with the myCAF/iCAF ratio, in contrast to other DDR components such as *ATR*, *PALB2*, and *BRCA1*, which showed strong positive associations (**Fig. 7D**), as previously reported (61). Consistently, while *BRCA1* and *ATR* expression showed no correlation with ROS-associated transcriptional program, reduced expression of *ATM* and *FANCG* in malignant ductal cells was significantly associated with ROS signature enrichment (**Fig. 7E**), in line with the functional ROS-myCAF axis identified earlier (**Fig. 5**). These findings not only reinforce the distinct stromal remodeling linked to ATM deficiency, but also point to non-redundant roles of specific DDR genes in shaping the CAF landscape of PDAC, underscoring the need for tailored stromal-targeted interventions.

**Figure 7.**
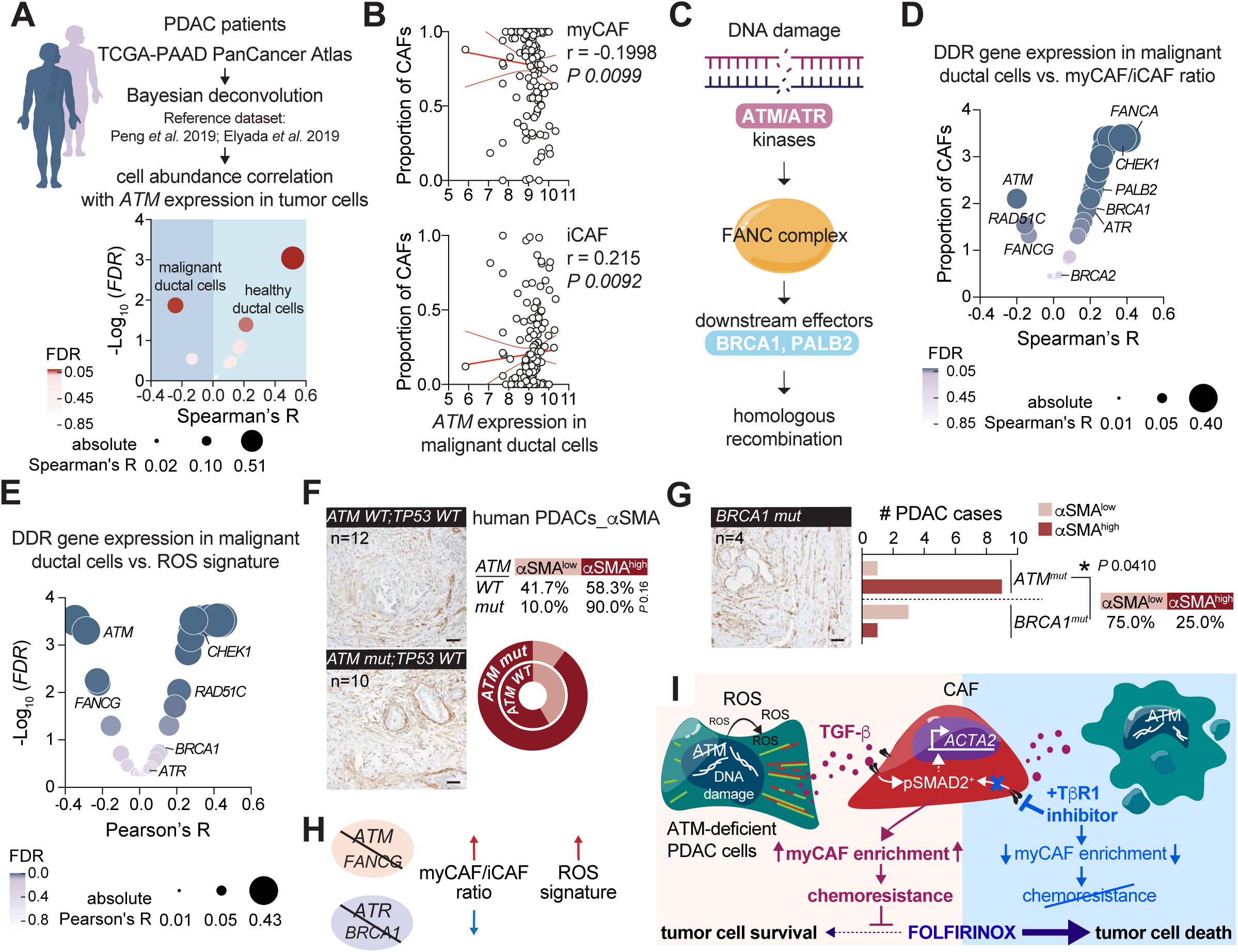
*ATM*-deficient human pancreatic cancers similarly display a myCAF and ROS-enriched environment. **A,** Schematic representation (top) of the workflow used to infer cell type composition and correlate ATM expression in tumor cells with stroma profiles. The BayesPrism method was applied to the TCGA-PAAD bulk RNA-seq dataset, using two reference datasets (17,60), to infer cell type composition. Correlation plot showing the results of Spearman’s correlation test between *ATM* expression and cell abundance. **B,** Dot plot showing the correlation between *ATM* mRNA expression in malignant ductal cells and proportions of inferred myCAFs (top) and iCAFs (bottom). Spearman’s correlation test. **C,** Schematic representation of DNA damage repair (DDR) signaling pathway and the DDR-related genes investigated in (**D–G**). **D** and **E,** Correlation plot showing the correlation between DDR-related genes expression in inferred malignant ductal cells and myCAFs/iCAFs ratio (**D**) and ROS signature (**E**). **F,** Immunohistochemistry staining for αSMA on human *TP53* wildtype and *ATM* mutated or not PDACs (n=22), and proportions of cases with αSMA low and high staining scores. Scale bars, 100 µm. Fisher’s exact test. **G,** Immunohistochemistry staining for αSMA on human *BRCA1* mutated PDACs (n=4), and proportions of cases with αSMA low and high staining scores. Scale bars, 100 µm. *, *P* < 0.05, Fisher’s exact test. **H,** Schematic representation illustrating the correlation between low *ATM*, *ATR*, *BRCA1*, and *FANCG* gene expression, ROS signature, and CAF differentiation in PDAC. **I,** Model recapitulating the protumorigenic tumor–CAF interplay operating in ATM-deficient pancreatic cancers, an actionable therapeutic target. HR, homologous recombination; ROS, reactive oxygen species; TCGA, The Cancer Genome Atlas.

To validate our findings, we used a unique set of clinically annotated *ATM*-mutated human PDACs to evaluate the correlation between ATM loss-of-function and myCAF enrichment (**Supplementary Table S2**). Most *ATM* pathogenic mutations are truncating, resulting in unstable products (62), whereas *TP53* mutations stabilize the protein, conferring oncogenic gain-of-function (63). Additionally, analysis of DDR gene mutation frequency and co-occurrence in the TCGA-PAAD cohort revealed a mutual exclusivity between *ATM* and *TP53* mutations (**Supplementary Fig. S12C**). This highlights the importance of focusing on *ATM*-mutated but *TP53* wildtype patients to specifically assess the impact of epithelial ATM loss-of-function on CAF fate. Our previous observations were substantiated by immunohistochemistry for the myCAF marker αSMA, showing a clear trend toward higher αSMA expression in peritumoral stroma of *ATM*-mutated PDAC compared to *ATM* wildtype tumors (**Fig. 7F**). To further explore genotype-specific effects on TME remodeling and the potential for targeted stromal interventions, we next analyzed *BRCA1-* and *ATR*-mutated PDAC models. Interestingly, αSMA staining in *BRCA1*-mutant tumors revealed a distinct CAF profile, with a predominance of αSMA^low^ cases, significantly contrasting with the αSMA-enriched phenotype observed in *ATM*-mutant PDAC (**Fig. 7G**). In parallel, to mechanistically assess the role of ATR in epithelial–stromal crosstalk, we generated CRISPR-mediated *ATR* knockout in KC cells (**Supplementary Fig. S12D**). Although ATR functionally complements ATM in HR repair, ATR deficiency did not recapitulate the ROS–TGF-β–myCAF axis seen in ATM-depleted cells, highlighting a distinct, non-redundant role for ATM in stromal remodeling (**Supplementary Fig. S12E–S12G**). Consistently, treatment of *ATR*-mutant PDX model with FOLFIRINOX plus galunisertib effectively controlled tumor growth, but offered no additional benefit compared to FOLFIRINOX alone (**Supplementary Fig. S12H**). Together, these comprehensive analyses underscore the pivotal role of ATM loss-of-function, and the broader epithelial HRD status, in orchestrating TME programing, as evidenced by the enriched myCAF pattern and heightened ROS stress in human ATM-deficient tumors (**Fig. 7H** and **I**).

## DISCUSSION

Achieving a one-size-fits-all approach in PDAC treatment is an unattainable goal, as the current stratification criteria primarily focus on cancer epithelial genetics and provide limited guidance for treatment. Most additional classifiers are mainly prognostic in nature and fall short of providing instructive treatment information. However, a comprehensive attack on the multifaceted intratumoral communication hubs involving both epithelium and niche could potentially add an extra layer to our personalized treatment arsenal. Nevertheless, caution is warranted due to the failure of initial clinical trials that targeted cancer–TME communication hubs such as Hedgehog signaling (19,23). These failures are likely due to the non-selective and non-genotypically stratified nature of these stroma-depleting interventions. In fact, the heterogeneity among CAF subpopulations, either supporting or restraining PDAC progression and therapeutic responses, might pose a significant constraint (64). Unfortunately, the underlying mechanistic determinants of this heterogeneity remain largely unexplored thus hampering so far, any genotype-tailored stromal interventions.

Along this line, the current study demonstrates that ATM in pancreatic cancer not only functions as a classical tumor suppressor of epithelial-intrinsic aggressive features, but also tightly triggers TGF-β release from the cancerous epithelium involving a ROS–actomyosin–contractility axis. Subsequently, TGF-β1 promotes a myCAF enrichment in both mouse and human cancers. Consequently, this forward programming axis enhances mesenchymal phenotypes in the neoplastic epithelium, thereby promoting cancer aggressiveness and extensive ECM deposition (5), as mirrored in a myCAF-enriched TME (16,24). Vice versa, interference with this crosstalk using clinical grade TGF-β inhibitors in combination with standard-of-care chemotherapy uncloses a druggable hallmark of ATM- and potentially other HRD-deficient pancreatic cancers (**Figure 7I**). Previously, we showed that a high degree of genomic instability in ATM-null PDAC is a direct consequence of HRD and the subsequent inability to properly repair DSBs (4,8). AKC cells subsequently upregulate P53, suggesting a compensatory activation that could restrain ATM loss-of-function effects on pancreatic carcinogenesis. Remarkably, codeletion of ATM and P53 led to highly invasive PDACs surpassing the severe oncogenic effects of single P53 loss (5,65,66), stressing the existence of autonomous roles for ATM and P53. Intriguingly, our study reveals that the discovered druggable stromal programming axis operates independently of P53 loss-of-function thereby widening the conceptual and translational application of our work. Previous studies have demonstrated that distinct P53 gain or loss-of-function alterations differentially educate CAFs, respectively promoting or not a permissive TME for tumor cell dissemination (67,68). However, the ATM loss specifically reshapes the identity of myCAFs, overshadowing the impact of P53 loss-of-function. This highlights the existence of a hierarchical relationship within the mutational landscape that determines stromal traits. Another recent study also emphasized the coexistence of distinct CAF subtypes in HRD PDAC with *BRCA* mutations, with a particular enrichment in an immune-regulatory CAF subset expressing clusterin (61). Accordingly, examining the correlation between HR-related gene expression (4) and CAF profiles in human PDACs revealed iCAF programming in *BRCA1*- and *ATR*-low tumors, contrasting with myCAF-enriched *ATM*-low cases. Supporting these observations, *BRCA1*- and *ATR*-mutated cases did not recapitulate the ROS–TGF-β–myCAF axis characteristic of ATM loss and exhibited limited sensitivity to TGF-β blockade. Our work therefore indicates the existence of distinct genotype-specific TME programs in human cancers which could be employed as personalized add-on therapies in combination with standards of care.

Tumor–stroma communication mainly occurs through secreted cues, the malignant cells providing the foremost factor to trigger feedback and forward signaling with their neighboring cells (69). Our secretome profiling revealed higher release of TGF-β1 in ATM-deficient PDAC cells, which was accompanied by the accumulation of the cytokine in tumor tissue and subsequent activation of SMAD-mediated canonical TGF-β pathway in αSMA^+^ myCAFs. Myofibroblastic CAF differentiation indeed derives from an active TGF-β signaling, suggested to depend on juxtacrine or paracrine signals produced by adjacent tumor cells (69). Interestingly, both ATM-deficient and proficient malignant cells produce IL1α, which is a driver of the iCAF fate. Consistently, previous studies have highlighted the antagonism between IL1α and TGF-β signaling in fibroblasts, the latter abolishing iCAF phenotype differentiation via suppressing IL1R1 expression (26). Accordingly, shared myofibroblastic programs and evolution trajectories of both prominent CAF subtypes in ATM-deficient tumors underlined the dynamic cell state transition toward myCAF fate. Furthermore, ATM deficiency is associated with impaired immunoregulatory signaling, which may further limit the capacity to induce inflammatory reprogramming of CAFs (70). Immune profiling of AKPC tumors revealed a distinct immune landscape, characterized by reduced B-lineage cells and diminished CD8^+^ T cell infiltration, consistent with recent findings (71). Given the reported sensitivity of ATM-null pancreatic cancers toward immune checkpoint inhibition, further investigation of immune composition and function, particularly in the context of TGF-β signaling inhibition, may uncover an additional therapeutic axis in this model (70).

The regulation of TGF-β bioavailability involves several mechanisms, with integrin-mediated activation being one of the best-understood pathways. This mechanism allows the conversion of cytoskeleton tension into physical forces on the secreted latent TGF-β complex, leading to the release of the active cytokine (41). Our data support a model in which ATM loss promotes the acquisition of an oncogenic contractile and migratory phenotype, in line with the previously reported EMT program in these cancers (5). Furthermore, it is accompanied by increased expression of integrins and proteins involved in actomyosin machinery (ACTN4), focal adhesion, and cytoskeleton dynamics (PXN, VINC). Under mechanical tension, PXN facilitates the recruitment of VINC at focal adhesion sites through a myosin-dependent mechanism, connecting integrins to the actin cytoskeleton and thus enabling the transmission of traction and contractility forces during cell motility (72). Cell spreading and integrin arrangement are promoted through cytoskeleton remodeling, which is triggered by ROS-mediated actin oxidation, leading to actomyosin contractility and the formation of stress fibers. ATM deletion, by generating an HRD phenotype, leads to the accumulation of DNA damage (4,8), which in turn causes ROS production (50). Thus, we propose that mitochondrial reprogramming and elevated ROS levels in ATM-deficient PDACs enhance actomyosin contractility, thereby altering the tumor cell secretome to drive CAF reprogramming. This is supported by our ROS interference experiments, which not only promoted a more epithelial phenotype but also reduced TGF-β secretion. It is worth noting that pharmacological ATM inhibition alone was sufficient to elevate ROS production and TGF-β1 secretion, indicating that stromal reprogramming arises directly from loss of ATM kinase activity rather than irreversible HRD-associated genomic alterations. Interestingly, our study also highlighted transcription factors in malignant ATM-deficient cells, i.e., *Atf6* and *Srebp*, that play key roles in endoplasmic reticulum stress response, intimately linked to redox homeostasis (73). The predicted increased activity of these factors may indicate compensative events in response to the disrupted redox equilibrium caused by ATM loss-of-function in PDAC cells.

CAFs can promote chemoresistance in PDAC through various means (74,75). Indeed, while TGF-β1 knockout in ATM-deficient PDAC cells reduced their migratory capacity, it did not increase their sensitivity to FOLFIRINOX, suggesting that chemoresistance is not cell-autonomous in this context, but instead driven by the ATM loss-induced TGF-β-dependent myCAF programming. Most studies have primarily focused on the negative impact of the stromal physical barrier on drug delivery (67). Consistent with this notion, a significant stroma remodeling upon TGF-β pathway interference with less ECM-producing myCAFs and fibrosis was accompanied by an improved response to FOLFIRINOX *in vivo* in ATM-null tumors, independently of the P53 loss-of-function on the neoplastic epithelium. However, investigating cancer cell–PSC interactions *in vitro*, using a transwell-based indirect co-culture system, revealed an ECM-independent role of αSMA^+^ myCAF-secreted factors in promoting resistance to the cytotoxic effects of oxaliplatin in ATM-deficient cells. CAFs have the ability to sustain PDAC cells by antagonizing drug processing, to foster resistance to gemcitabine (76). Additionally, CAFs may significantly contribute to limit efficacy of antitumor therapies by scavenging drugs (77). Future studies need to clarify the exact molecular mechanisms through which different CAF subtypes exert their oncogenic functions. To date, clinical trials evaluating the efficacy of gemcitabine in combination with TGF-β signaling interference have shown modest but promising benefits even in non-selected pancreatic cancers (45,46). Consistent with a previous study that reported no tumor control in KPC mice following TGF-β interference with galunisertib (26), the therapeutic benefit observed in our study was specific to ATM-depleted PDACs. This suggests that combined TGF-β pathway interference and chemotherapy may be most effective in stratified subsets of PDAC patients based on ATM loss-of-function or other HRD-causing genotypes. However, considering the complex role of TGF-β in cancer, further investigation is warranted to identify potential biomarkers or genomic signatures that could be used to stratify patients for personalized therapies. Importantly, perturbation of TGF-β signaling axis implicates also alterations of the immune compartment. Tumor-associated macrophages and regulatory T cells have been reported to operate protumorigenic in pancreatic cancer by establishing an immunosuppressive TME (78) through TGF-β-mediated mechanisms (79). Accordingly, anti-TGF-β treatment could redraft stromal immune content toward a proinflammatory state, potentially enhancing antitumor immunity. Moreover, our data demonstrate that TGF-β blockade in ATM-deficient tumors shifts the balance of CAF subsets, driving an iCAF expansion. This adaptative reprogramming, consistent with previous reports (26), may limit long-term therapeutic benefit of TGF-β inhibition by fostering pro-tumorigenic signaling. Nevertheless, our study also demonstrates a greater therapeutic efficacy in ATM-deficient compared to ATM-proficient PDAC, underscoring the importance to decipher and incorporate context-dependent TME functions when designing stromal-targeted therapies. In this regard, our data further show that myCAF differentiation induced by ATM-deficient tumor cells is not strictly constrained by fibroblast ontogeny. While tumor genotype serves as a dominant driver of stromal reprogramming, the cellular origin of CAF also shapes their transcriptional profiles, warranting further investigation into whether these differences translate into distinct roles within the TME.

In conclusion, our study reveals a novel mechanistic link between ATM deficiency and the activation of TGF-β signaling in CAFs, resulting in their reprogramming into myCAFs, which in turn promotes PDAC aggressive features. The efficacy of our interference approach relies on the dual targeting of the intrinsic ATM-deficient cancer cell vulnerabilities (platin-sensitive) and the subsequent perturbation of extrinsic genotype-specific tumor–TME crosstalk (TGF-β-driven). This may represent a promising therapeutic strategy for a subset of PDAC patients harboring ATM loss-of-function or HRD. Although 17-25% of PDAC harbor DDR mutations, our data reveal variability in CAF profiles both within individual genotype and across different DDR-mutated tumors. This underscores the need for robust validation of genotype-specific stromal patterns before clinical translation. Our work thus suggests that the adequate targeting of a genotype-specific tumor-promoting dialog can be exploited to effectively abrogate oncogenic stromal features and enhance treatment response. Such a novel concept will encourage to profile and stratify mutational landscapes and gene expression levels in designing targeted interventions according to the subsequently arising stromal features to improve efficacies of future stroma-targeting strategies.

## ACKNOWLEDGMENTS

The authors especially thank Ariel Asuzano, Randa Kalaajieh, Yvonne Klöpfer, Katrin Köhn, Ralf Köhntop, Claudia Längle, Natalie Paul, Julia Ragg, and Sandra Widmann for their outstanding technical support. We thank Jana-Romana Fischer and Birgit Ott for excellent technical assistance. The authors thank Dr. Ninel Azoitei for helpful scientific exchange. The authors also thank Dr. Tim Eiseler and Prof. Dr. Volker Rasche for helpful scientific discussions. The authors thank the animal facility platform Tierforschungszentrum, Ulm University, and its members for animal care. We also thank the IPC-CRCM Experimental Pathology Platform (ICEP) for methodology, technical assistance and for anatomical pathology on patient-derived xenografts. The authors also thank the organoid core facility for sharing resources and providing support. Main funding was provided by the Deutsche Krebshilfe (German Cancer Aid) grants 70114761 to A. Kleger. Additional funding to A. Kleger came from the Deutsche Forschungsgemeinschaft (DFG) „Heisenberg-Programm“ KL 2544/6-1. Additional funding to L. Perkhofer came from the Deutsche Krebshilfe (German Cancer Aid) grants 70115292 and Deutsche Forschungsgemeinschaft (DFG) PE 3337/1-1. E. Roger, F. Arnold, and M. Hirschenberger received funding by the Bausteinprogramm of Ulm University (respectively, L.SBN.0193, L.SBN.0206, and L.SBN.0245). E. Roger was supported by the Hertha-Nathorff program (Ulm University Hospital). M.K. Melzer received funding from the University of Ulm in the Clinician Scientist Program and is additionally funded by the Else Kröner Research School for Physicians. K.M.J. Sparrer was supported by a Grant from the German Federal Ministry of Research, Technology and Space (IMMUNOMOD-01KI2014) and the German Research Foundation (CRC1279). A. Kleger and T. Seufferlein are speakers of an Else Kröner Research School for Physicians. A. Kleger and T. Seufferlein are PIs and F. Arnold and D. Srinivasan are students of HEIST RTG funded by the DFG GRK 2254/1. We are deeply grateful to Kuhn Elektro-Technik GmbH for supporting our research to fight pancreatic cancer.

## AUTHORS’ CONTRIBUTIONS

Conceptualization, E.R., J.G., A.K.; Methodology, E.R., J.G.; Formal Analysis, E.R., H.M.M., E.Z., D.S., A.H., Y.L., T.E., C.L., C.M., L.H., K.K., M.W., M.S.S.A., J.B., M.Ha., K.S., J.G., ; Investigation, E.R., H.M.M., E.Z., D.S., A.H., L.M., A.K.B., R.S., M.K.M., Y.L., A.S., L.H., F.A., M.M., B.N., J.S., L.G., J.P.M., C.L., C.M., M.K., A-L.V., M.W., M.S.S.A., K.S., L.P., J.G.; Resources, M.M., M.Hi., V.H., J.D.H., J.-M.L., T.S., A.A., R.K., R.R., M.A.M., P.C.H., K.M.J.S, C.J.H., K.J.G., A.S., N.D., L.P., J.G., A.K.; Writing – Original Draft, E.R., J.G., A.K.; Visualization, E.R.; Supervision, E.R., J.G., A.K.; Project Administration, E.R., L.P., J.G., A.K.; Funding Acquisition, E.R., L.P., J.G., A.K.

## METHODS

### Ethics statement

All animal care and procedures followed German, US, or French legal regulations and were previously and respectively approved by the governmental review board of the state of Baden-Württemberg (Permission no. 1273, and 1568), by the Institutional Animal Care and Use Committee of the University of California Irvine (protocol #AUP-23-084), by the Ethics Committee for animal experimentation of Marseille no.14 (approval number APAFIS 202308111552762). All mouse work aspects were carried out following strict guidelines to insure careful, consistent and ethical handling of mice. The human biomaterial used was provided by the biobank of the University Hospital of Ulm following the regulations of the Biobank and the vote of the Ethics Committee of the University of Ulm (PDAC Liver Metastasis): 72/2019 (human tissue and blood), and by the Universitätsklinikum Heidelberg following the regulations and the vote of the Ethics Committee of the Universitätsklinikum Heidelberg (Permission no. S-315/2020).

### Mice

*Atm^fl/fl^*, *Trp53^fl/fl^*, *LSL-Kras^G12D/+^*, and *Ptf1a^Cre/+^* were previously described (80–83). Male C57BL/6J mice were purchased from Janvier Labs. Female Nude-*Foxn1^nu^* mice were purchased from Inotiv. Mice were housed and bred in a conventional health status-controlled animal facility. All the aspects of the mouse work were carried out following strict guidelines to insure careful, consistent, and ethical handling of mice.

### Cell culture

*LSL-Kras^G12D/+^*; *Ptf1a^Cre/+^* (KC), *Atm^fl/fl^*; *LSL-Kras^G12D/+^*; *Ptf1a^Cre/+^*(AKC), *Trp53^fl/fl^*; *LSL-Kras^G12D/+^*; *Ptf1a^Cre/+^* (KPC) and *Atm^fl/fl^*; *Trp53^fl/fl^*; *LSL-Kras^G12D/+^*; *Ptf1a^Cre/+^*(AKPC) cells were respectively isolated from *LSL-Kras^G12D/+^*; *Ptf1a^Cre/+^*, *Atm^fl/fl^*; *LSL-Kras^G12D/+^*; *Ptf1a^Cre/+^*, *Trp53^fl/fl^*; *LSL-Kras^G12D/+^*; *Ptf1a^Cre/+^,* and *Atm^fl/fl^*; *Trp53^fl/fl^*; *LSL-Kras^G12D/+^*; *Ptf1a^Cre/+^*mice and immortalized as described previously (8). Cells were cultured in DMEM, containing 10% FBS and P/S (100 IU/mL penicillin and 100 μg/mL streptomycin sulfate). MIA PaCa-2 (ATCC CRL-1420) cells were purchased from the ATCC and were cultured in DMEM, containing 10% FBS and P/S. The CRISPR-edited ATM-deficient MIA PaCa-2 cell lines were generated as previously described (4). Human pancreatic stellate cells (PSCs) (35) were kindly provided by Prof. J.-Matthias Löhr (Karolinska Institute) and were cultured in DMEM, containing 10% FBS and P/S. CAF lines were kindly provided by Prof. Dr. Dieter Saur (Technical University of Munich) and were cultured in DMEM, containing 10% FBS and P/S. Briefly, CAF lines were derived from primary PDAC cultures established from autochthonous pancreatic tumors (*Kras^G12D/+^*; *Trp53^+/lox^*; *Tgfbr2^lox/lox^* and *Kras^G12D^*^/+^; *Tgfbr2^lox/lox^*) according to protocols adapted from (16). PCR was performed to assess the absence of recombined *Kras* band and select CAF lines. HEK-293T cells were kindly provided by Prof. Dr. Franz Oswald (Ulm University Hospital) and were cultured in DMEM, containing 10% FBS and P/S. Patient-derived organoids lines were propagated as previously described (84).

All cells were propagated at 37°C under 5% (v/v) CO_2_ atmosphere. All experiments were performed between passage 5 and 25. All experiments were conducted with at least two different cell lines per genotype. Mycoplasma tests were regularly performed using the MycoSPX PCR kit (Biontex).

5-fluorouracil (NSC 19893), AZD0156, galunisertib (LY2157299), irinotecan (CPT-11), leucovorin (folinic acid), and oxaliplatin (L-OHP) were purchased from Selleckchem. SB431542 was purchased from Axon. MRTX1133 was purchased from Biorbyt. Human IL1α was purchased from Bio-Techne. Human TGF-β1 was purchased from Peprotech. YCG063 and menadione were purchased from Merck-Millipore.

Tumor cell conditioned media were prepared from 1.7 × 10^6^ cells propagated in serum-free DMEM containing P/S for 24 h in 10 cm dishes. Supernatants were harvested, centrifuged for cell debris elimination, and stored at -80°C. For assessing cancer-associated fibroblast differentiation, PSCs were grown on Matrigel GFR-coated wells (1:4 dilution) in serum-free DMEM containing P/S. For co-culture approaches assessing PSC differentiation, tumor cells and PSCs (1:1 ratio; respectively, 50,000 tumor cells in 6-well translucent polystyrene cell culture inserts (0.4 µm pore size, ThinCert, Greiner Bio-One) and 50,000 PSCs) were propagated in serum-free DMEM containing P/S. All analyzes to investigate PSC differentiation into cancer-associated fibroblasts were conducted using the vehicle inactivated PSC condition as a reference for normalization.

### CRISPR/Cas9 genetic editing

Parental AKC and AKPC tumor cells were nucleofected with a complex consisting of HiFi Cas9 Nuclease V3 (80 pmol, Integrated DNA Technologies) and sgRNA (300 pmol, Integrated DNA Technologies), using either specific sgRNAs or a non-targeting control sgRNA (Integrated DNA Technologies; **Supplementary Table S4**). Nucleofection was carried out on a Lonza 4D-Nucleofector system using the P2 Primary Cell 4D-Nucleofector X kit (Lonza, V4XP-2032) and pulse code DN100. Effective knockout of target genes was tested at protein level using western blotting method or by Sanger sequencing (Eurofins Genomics).

### Colony formation assay

Tumor cells were seeded in 24-well plates and co-cultured with pancreatic stellate cells seeded (1:2 ratio; respectively, 12,000 tumor cells and 24,000 PSCs) in Matrigel-coated polycarbonate cell culture inserts (3 µm pore size, Nunc), in serum-free DMEM containing P/S for 24 h prior treatment. Co-cultures were then treated for 48 h with 6 µM oxaliplatin. Tumor cells were fixed with cold 4% formaldehyde and stained with 4% Giemsa. Images were captured using an iPhone 11 (Apple). Acquired pictures were subsequently analyzed using ImageJ software (National Institutes of Health).

### Cell viability assay

Cells were seeded in 96-well plates (1,000 to 5,000 cells/well). After cell attachment, cells were treated for 3 days. Cell viability was analyzed with a MTT assay (Sigma-Aldrich) accordingly to manufacturer’s instructions. Absorbance was measured at 590 nm wavelength using a spectrophotometer (Tecan Infinite M200 Pro). Viability percentages were normalized to vehicle-treated cell viability.

### TGF-**β** reporter assay

HEK-293T cells, used as recipient cells for TGF-β reporter assay, were kindly provided by Prof. Dr. Franz Oswald (Ulm University Hospital). HEK-293T cells were transfected with the pGL3-(CAGA)12-MLP-Luc (85) and Renilla (phRL-CMV) plasmids, prior treatment with tumor cell conditioned media. Briefly, HEK-293T cells were transfected using 2 µL of Lipofectamine 2000 reagent (Invitrogen) in Opti-MEM medium. After 6 h, the medium was replaced with conditioned media from tumor cells or serum-free DMEM medium containing P/S. After 48 h, the Dual-Luciferase Reporter Assay (Promega) was used following the manufacturer recommendations and the luminescence values were determined for each condition with a spectrophotometer (Tecan Infinite M200 Pro). Renilla luminescence values were used for correction to determine Luciferase luminescence corrected values. Luciferase luminescence corrected values were normalized to untreated transfected HEK-293T cells, used as a reference for basal reporter activity.

### TGF-**β**1 ELISA

Tumor cells were propagated for 24 h in serum-free DMEM containing P/S. Supernates were collected and processed following the manufacturer instructions (Quantikine Human/Mouse/Rat/Porcine/Canine TGF-beta 1 Immunoassay, R&D Systems).

### ROS levels quantification

Ten thousand tumor cells were seeded in black 96-well plate in DMEM containing 10% FBS and P/S. After cell attachment, the culture medium was replaced with serum-free DMEM containing P/S for 24 h and processed following the manufacturer recommendations for ROS levels quantification (ROS-Glo H2O2 Assay, Promega). Luminescence was measured with a spectrophotometer (Tecan Infinite M200 Pro).

### Microchannel single cell migration assay

The microchannel chips were processed as previously described (49). Collagen I-coated chips and 11 µm wide microchannels were used to assess single-cell migration. Ten thousand tumor cells were added to the invasion chip and cultured upon 0 to 10% FBS gradient. Phase contrast live-cell imaging was performed in a temperature-controlled culture chamber equipped with an Axiovert 200M inverted microscope (Carl Zeiss). Images were captured every 10 min for 24 h. Acquired pictures were subsequently analyzed using ImageJ software. Permeative cells were defined as tumor cells that left the 0% FBS chamber, entered the channels, and exited the opposite 10% FBS side.

### Wound healing assay

Assays using the AKC tumor cell lines treated or not with ROSi were conducted as follow: three hundred thousand tumor cells were seeded in 48-well plate in DMEM containing 10% FBS and P/S. After cell attachment, the tumor cell layer was scratched using sterile tips and the culture medium was replaced with serum-free DMEM containing P/S for 48 h. Brightfield images were captured after 0, 24 and 48 h. Acquired pictures were subsequently analyzed using ImageJ software.

Assays using CRISPR-edited AKC and AKPC tumor cell lines were conducted using the Incucyte SX5 system (Sartorius) as followed: 50,000 cells/well were seeded in Sartorius Incucyte ImageLock 96-well microplates in DMEM containing 10% FBS and P/S. After cell attachment, the Incucyte 96-well WoundMaker tool was used to generate uniform scratch wounds across all wells and the medium was replaced with DMEM 0% FBS and P/S. Wound closure was monitored for 24 h. The percentages of wound closure were determined with the following equation: (Tx wound area/T0 wound area)*100.

### *Ex vivo* co-culture assay on porcine urinary bladder

Porcine urinary bladders (PUBs) were processed as described previously (36). Briefly, PUBs were deepithelialized and sterilized with 0.1% peroxyacetic acid, before conducting air-liquid interface organ culture in 6-well plates containing DMEM-F12 supplemented with 2% P/S. Tumor cells and pancreatic stellate cells (1:3 ratio; respectively, 166,000 tumor cells and 500,000 PSCs) were seeded inside 5 mm-diameter plastic rings on the PUB surface in a 1:1 serum-free DMEM-F12:Matrigel GFR solution. After 5 days, co-cultures were treated for 3 days with FOLFIRINOX (75 nM 5-fluorouracil, 25 nM irinotecan, and 5 nM oxaliplatin) and/or 10 µM of the TGFBR1 inhibitor SB431542. PUBs were then fixed and processed for histology as previously described (36).

### Immunostaining

For immunocytofluorescence, cells were grown on glass coverslips, fixed in cold 4% formaldehyde, and permeabilized with 0.05% Tween 20 for 20 min before immunofluorescence experiments. Matrigel GFR coating of glass coverslips was performed prior pancreatic stellate cells seeding. F-actin was stained with phalloidin-Atto565 (1:500-1:1,000; Sigma-Aldrich). All histological experiments were performed as previously described (5,8) or according to standard protocols on formalin-fixed paraffin-embedded tissues.

For immunofluorescence, primary antibodies against ACTN4 (1:250; clone EPR2533(2); Abcam ab108198), αSMA (1:500-1:1,000; clone 1A4; Abcam ab7817), B220 (1:25; clone RA-6B2; Santa Cruz Biotechnology sc-19597), CD11b (1:100; clone M1/70; Cell Signaling 46512), CDH1 (1:250; clone 24E10; Cell Signaling 3195), CDH2 (1:200; clone D4R1H; Cell Signaling 13116), CK19 (1:35; clone TROMA-III; DSHB 2133570), CTGF (1:50 clone E-5; Santa Cruz Biotechnology sc-365970), CXCL2 (1:50; clone EPR28746-89; Abcam ab317569), H2AX p-S139 (1:1,000; clone 20E3; Cell Signaling 9718), Ly6C (1:200; clone RM1151; Abcam ab317272), P53 (1:2,000; clone 1C12; Cell Signaling 2524), PDPN (1:200; clone 8.1.1; BioLegend 127404), POSTN; CD11b (1:100; clone E4K8C; Cell Signaling 93169), PXN (1:100; clone 5H11; Invitrogen AHO0492), SMAD2 p-S465/467 (1:100-1:200; clone 138D4; Cell Signaling 3108), STAT3 p-Y705 (1:200; clone D3A7; Cell Signaling 9145), TJP1 (1:250; clone 1A12; Invitrogen 33-9100), and VINC (1:250; clone 42H89L44; Invitrogen 700062) were used.

The secondary antibodies were anti-rabbit Alexa Fluor 647 dye-conjugated (Invitrogen A21244), anti-mouse Alexa Fluor 488 dye-conjugated (Invitrogen A11029), anti-mouse Alexa Fluor 568 dye-conjugated (Invitrogen A-11004), and anti-rat Alexa Fluor 568 dye-conjugated (Invitrogen A11077), and were used at a 1:500 dilution. For CK19 immunofluorescence, biotin-conjugated anti-rat antibodies and FITC-conjugated streptavidin were used. DNA was stained with DAPI contained within the ProLong Diamond Antifade mounting medium (Thermo Fischer Scientific).

For immunohistochemistry, primary antibodies against αSMA (0.03 µg/mL; clone 1A4; Abcam ab7817), CDH1 (1:1,000; clone 24E10; Cell Signaling 3195), CK19 (1:35; clone TROMA-III; DSHB 2133570), cleaved caspase-3 (1:1,000; clone 5A1E; Cell Signaling 9664), and Ki67 (1:100; clone SP6; Invitrogen MA5-14520) were used.

Brightfield and immunofluorescence images were acquired at ambient temperature using a Plan-Apochromat 20×/0.8 M27 objective mounted on a Zeiss AxioObserver.Z1/7 microscope (Carl Zeiss) equipped with AxioCam 506 color camera and AxioCam 702 mono cameras (Carl Zeiss). Polarized light images were acquired using a Zeiss Axioscan7 slide scanner (Carl Zeiss). Acquired pictures were subsequently analyzed using ImageJ/Fiji software or QuPath (86). The “Directionality” ImageJ plugin was used to infer the preferred orientation of collagen fibers in the picrosirius red polarized input images. All quantifications were performed on at least three random pictures. All analyzed pictures were carefully checked by eye to exclude artifacts and false positive areas. Immunostainings for αSMA on human PDAC tissues were carefully checked and analyzed by a board-certified pathologist.

### Western blotting

Total protein extracts were prepared using RIPA lysis buffer (50 mM Tris, pH 7.5, 150 mM NaCl, 1% Nonidet P-40, 0.5% sodium deoxycholate, 0.1% SDS and commercial protease and phosphatase inhibitor cocktail tablets (Roche)) and were subjected to electrophoresis on SDS/PAGE. The separated proteins were transferred onto PVDF membranes (Millipore) by electroblotting. Primary antibodies against ACT1B (1:1,000; clone C4; Santa Cruz Biotechnology sc-47778), ACTN4 (1:1,000; clone EPR2533(2); Abcam ab108198), ATM (1:1,000; clone Y170; Abcam ab32420), GAPDH (1:1,000; clone 14C10; Cell signaling 2118), KEAP1 (1:1,000; clone D6B12; Cell Signaling 8047S), PXN (1:1,000; clone 5H11; Invitrogen AHO0492), TGF-β1 (1:1,000; clone EPR21143; Abcam ab215715) were used. The second antibodies were anti-mouse IgG, HRP-linked whole Ab (from sheep) (Amersham ECL NA931) and anti-rabbit IgG, HRP-linked whole Ab (from donkey) (Amersham ECL NA934), both used at a 1:5,000 dilution.

### Flow cytometry on cancer-associated fibroblasts

For isolating cancer cells and CAFs, AKPC and KPC tumors were processed as previously described (5). CAFs were sorted using the mouse Tumor-Associated Fibroblast Isolation Kit (Miltenyi Biotec) according to the manufacturer’s protocol. For flow cytometric analyses, cells were fixed in cold 4% formaldehyde, permeabilized with 90% methanol at 4°C for 15 min, and incubated with antibodies for 30 min. Antibodies used were anti-αSMA (1:1,000; clone 1A4; Abcam ab7817) and anti-mouse Ly6C-APC-eFluor780 (0.5 µg/reaction; clone HK1.4; Invitrogen 17.5932.82). Secondary antibody was anti-mouse Alexa Fluor 488 dye-conjugated (1:500; Invitrogen A110291). Cells were sorted on an Attune NxT Acoustic Focusing Cytometer (Thermo Fisher Scientific).

### qPCR

Total RNAs were extracted using the RNeasy Mini Kit (Qiagen). First-strand cDNAs were prepared using 250 ng of RNA and SuperScript II Reverse Transcriptase in the presence of random primers (Thermo Fisher Scientific) according to the manufacturer protocol. Quantitative PCR were performed using an Applied Biosystems QuantStudio 3 System (annealing temperature 60°C) and PowerUp SYBR Green Master Mix (Thermo Fisher Scientific). As described previously (4), all the real-time values were averaged and compared using the threshold cycle (CT) method, where the amount of target RNA (2^-ΔΔCT^) was normalized to the endogenous expression of *18S* (ΔCT). The amount of target mRNA in control cells was set as the calibrator at 1.0. Primers used for quantitative RT-PCR were purchased from Sigma-Aldrich (KiCqStart SYBR Green Primers: *ACTA2*, *CXCL12*, *IL6*, *PDGFRA*, *PDGFRB*, and *POSTN; Acta2*, *Cxcl1*, *Il11*, *Il6*, *Postn*, *Pdgfrb*, and *Tagln*) and Biomers (*18S*: Fw: 5’-GTAACCCGTTCCCATT-3’, Rev: 5’-CCATCCAATCGTAGCG-3’; *CTGF*: Fw: 5’-CCTGGTCCAGACCACAGAGT-3’, Rev: 5’-TGGAGATTTTGGGAGTACGG-3’; *CXCL2*: Fw: 5’-TGCAGGGAATCTCAAG-3’, Rev: 5’-TGAGACAAGCTGCCCA-3’; mouse *18S*: Fw: 5’-GTAACCCGTTCCCATT-3’, Rev: 5’-CCATCCAATCGTAGCG-3’; mouse *Tgfb1*: Fw: 5’-TGGAGCAACATGTGGAACTC-3’, Rev: 5’-GTCAGCAGCCGGTTACCA-3’).

### Sample collection and processing for single cell analyses

PDAC tumors were harvested from two female KPC and two female AKPC mice and snap-frozen. 20-30 mg tissue samples were resuspended in 1ml of wash buffer (0.25 M sucrose, 25 mM KCI, 5 mM MgCl_2_, 10 mM Tris-HCI (pH 7.5), 1 mM DTT (Sigma, D9779), 1X protease inhibitor (Fischer, PIA32965), 1□U□µl^−1^ recombinant RNAsin (Promega, PAN2515), 0.1% IGEPAL (Sigma, I8896) in molecular biology water) and then homogenized using a Dounce homogenizer with 2 pestles. First, a loose pestle was used for 5-10 strokes to break the tissue into small pieces and then a tight pestle was used for 15-25 strokes to fully homogenize the sample into a uniform solution. Homogenized tissue was filtered through a 30 μm filter (CellTrics, Sysmex) and then centrifuged with a swinging-bucket centrifuge (100 rcf, 10 min, 4°C, Eppendorf, 5920 R). Nuclei were resuspended in 1 ml of wash buffer (0.25 M sucrose, 25 mM KCI, 5 mM MgCl_2_, 10 mM Tris-HCI (pH 7.5), 1 mM DTT (Sigma, D9779), 1X protease inhibitor (Fischer, PIA32965), 1□U□µl^−1^ recombinant RNAsin (Promega, PAN2515) in molecular biology water) and centrifuged again (100 rcf, 10 min, 4°C, Eppendorf, 5920 R). Next, nuclei were resuspended in 400 µl of sorting buffer (1X protease inhibitor (Fischer, PIA32965), 1□U□µl^−1^ recombinant RNAsin (Promega, PAN2515), 1% fatty acid free BSA (Proliant, 7500804), 2uM 7-AAD (Invitrogen, A1310) in PBS (Corning, 21-040-CV)) and incubated on ice for 10 minutes. 120,000 nuclei were sorted into a Lobind tube with 90 µl collection buffer (5□U□µl^−1^ recombinant RNAsin (Promega, PAN2515), 5% fatty acid free BSA (Prolian, 7500804) in PBS (Corning, 21-040-CV) using an SH800 cell sorter (Sony). The resulting nuclei suspension was centrifuged again (500 rcf, 5 min, 4°C, Eppendorf, 5920 R) and resuspended in 100 µl of collection buffer (1% fatty acid free BSA (Proliant, 7500804), 1X protease inhibitor (Fischer, PIA32965), 10 mM Tris-HCI (pH 7.4), 0.1% Tween-20 (Sigma, P7949-100ML), 0.1% IGEPAL (Sigma, I8896), 0.01% digitonin (Promega, G9441), 1mM DTT (Sigma, D9779), 1□U□µl^−1^ recombinant RNAsin (Promega, PAN2515), 3 mM MgCl_2_ and 10 mM NaCl in molecular biology water). This suspension was incubated on ice for 1 minute and centrifuged again (500 rcf, 5 min, 4°C, Eppendorf, 5920 R). The supernatant was carefully removed without disturbing the pellet and 10 µl of 1X Nuclei Buffer (10x Genomics) was used to gently resuspend the pellet. Nuclei were counted using a hemocytometer and approximately 20,000 nuclei were used as inputs for the tagmentation reaction.

### Paired single nucleus RNA-seq and ATAC-seq

Nuclei from the two KPC mice were pooled together and nuclei from the two AKPC mice were also pooled together, for a total of 2 sets of libraries. Then single cell multiome ATAC and gene expression libraries were generated according to manufacturer instructions (Chromium Next GEM Single Cell Multiome ATAC□+□Gene Expression Reagent Bundle, 10x Genomics, 1000283; Chromium Next GEM Chip J Single Cell Kit, 10x Genomics, 1000234; Dual Index Kit TT Set A, 10x Genomics, 1000215; Single Index Kit N Set A, 10x Genomics, 1000212). The following PCR cycles were used: 7 cycles for ATAC index PCR, 7 cycles for complementary DNA (cDNA) amplification, and 13–16 cycles for RNA index PCR. A Qubit fluorometer (Life Technologies) was used to quantify final libraries and a TapeStation (High Sensitivity D1000, Agilent) was used to check size distribution. Finally, libraries were sequenced on NextSeq 500 and NovaSeq 6000 sequencers (Illumina) with the following read lengths (Read1□+□Index1□+□Index2□+□Read2): ATAC (NovaSeq 6000), 50□+□8□+□24□+□50; ATAC (NextSeq 500 with custom recipe), 50□+□8□+□16□+□50; RNA (NextSeq 500, NovaSeq 6000), 28□+□10□+□10□+□90.

### Paired single nucleus RNA-seq and ATAC-seq data quantification and alignment

Raw fastq files from both libraries were processed with 10x Genomics Cell Ranger ARC (v2.0.0) count command. This command aligned all reads to the mm10 mouse genome reference and created bam files and raw feature barcode matrices for both snRNA and snATAC.

### Paired single nucleus RNA-seq and ATAC-seq quality control and doublet identification

First, barcodes associated with empty droplets or multiplets, droplets with multiple barcodes and only one nucleus found by CellRanger ARC count, were identified, and removed. Barcodes were considered empty droplets if they had less than or equal to 500 unique RNA genes with counts or less than or equal to 1,000 unique ATAC fragments. To retain only data from high quality barcodes, additional filters were applied. Barcodes were removed if they had RNA percent mitochondrial reads greater than 5% or equal to or more than 5,000 ATAC mitochondrial reads. These counts from background contamination were removed from the snRNA-seq counts matrix using SoupX (87) (v1.6.2). SoupX identifies genes composing the background from empty droplets, uses these to estimate the proportion of contamination per barcode, and then lowers counts associated with all background genes for each cell.

Doublets were identified using Scrublet (88) (v0.2.3) on each library separately. Scrublet creates simulated doublet signatures using the input data clusters and then calculates a per-barcode doublet likelihood score based on the similarity to synthetic doublets. Scrublet was run on each library separately with an expected doublet rate of 0.06. All barcodes with a Scrublet doublet likelihood score above 0.25 were considered doublets and removed. After removing low quality barcodes and doublets, 10,404 high quality cells remained from both libraries.

### Paired single nucleus RNA-seq and ATAC-seq dimensionality reduction and clustering

Before dimensionality reduction and clustering, all snATAC-seq fragments were quantified across 5kb regions tiling the genome, to create a windows-based counts matrix. Seurat v4 (4.3.0) and Signac (89) (1.10.0) were used to perform all subsequent analyses. First data normalization and dimensionality reduction were performed on the snRNA and snATAC data separately. snRNA-seq counts were normalized using SCTransform (90) (v0.4.0), reduced to principal components with PCA and then projected into 2-dimensional space using UMAP (arXiv 2020 180203426) on the top 50 PCs. snATAC-seq counts were normalized with TFIDF, SVD was used to perform dimensionality reduction and then the top 2 through 50 dimensions were used to run UMAP. The single top dimension was excluded as it usually correlates with sequencing depth. Seurat’s Weighted Nearest Neighbor (91) clustering was used to combine information from both assays together into one dimensionality reduction map. The FindMultiModalNeighbors command was run using the top 50 RNA dimensions and top 2 through 50 ATAC dimensions. Then UMAP was run, and clustering was performed with the Leiden algorithm (92) and a resolution of 0.5. Both libraries were dimensionality reduced and clustered separately first to ensure they were good quality and then integrated together using Harmony (93) batch correction (v1.0.1) and then reanalyzed following the same methods.

### Assignment of cell type identities

Cell types were identified by the cluster specific expression of known marker genes for PDAC tumors. We used previously characterized (17,32) mouse marker genes to classify the following cell types: acinar cells (Ctrb1), B cells (Bank1), dendritic cells (Cacnb3), ductal cells (Prom1), endothelial cells (Egfl7), myeloid cells (Adgre1) perivascular cells (Rhoj), T and NK cells (Il17r) and cancer associated fibroblasts (Dcn). To identify known subtypes of ductal cells and CAFs, the cells from each cell type were reclustered separately using a resolution of 0.5. EMT-like cells, a subtype of ductal tumor cells expressing markers of the epithelial-to-mesenchymal transition, were classified by the expression of (Hmga2). Previously identified CAF subpopulations were also detected after sub clustering all fibroblasts: myofibroblastic CAFs (Col12a1, Tnc, Col8a1), inflammatory CAFs (Scep1, Plpp3, Scara5), and antigen-presenting CAFs (Ptgis, Fth1, Cxadr).

### Analysis of lymphoid and myeloid cells

To identify subpopulations of lymphoid and myeloid cells, cell barcodes annotated as B cells, T and NK cells or myeloid cells were subset and reanalyzed using optimized parameters. snRNA-seq counts were normalized using SCTransform (90), regressing out mitochondrial reads. The top 2000 variable features were used for linear dimensionality reduction with PCA, followed by batch correction with Harmony (93), and projection into UMAP space using the top 17 (lymphoid)/21 (myeloid) harmony-corrected principal components. snATAC-seq peaks with 0 counts across remaining cells were removed, counts were normalized using TFIDF, before linear dimensionality reduction using SVD and batch integration with Harmony (93). The top 2 through 6 harmony-corrected LSI dimensions were used for UMAP embedding of the snATAC-seq data. RNA and ATAC modalities were integrated through the Weighted Nearest Neighbor (91) approach with 20 k-nearest neighbors. Clustering was performed using the Leiden algorithm at a resolution of 0.7 (lymphoid)/0.5 (myeloid). Lymphoid subtypes were annotated based on the expression of canonical marker genes: B cells (*Bank1*, *Pax5*, *Ebf1*), CD4^+^ T cells (*Il7r*, *Cd3g*, *Cd4*), CD8^+^ T cells (*Il7r*, *Cd3g*, *Cd8a*, *Cd8b1*), Tregs (*Il7r*, *Cd3g*, *Cd4*, *Foxp3*, *Ctla4*) and NK cells (*Klrb1b*, *Klrk1*, *Gzma*). Myeloid clusters were annotated based on previously defined marker gene signatures (34), and signature marker genes: M1-like TAMs (*Cd80*, *H2-Aa*, *H2-Ab1*), M2-like TAMs (*Arg1*, *Fn1*), phagocytotic TAMs (*Mrc1*, *Cd163*) and proliferating TAMs (*Mki67*, *Top2a*).

### Identification of cell type peaks from snATAC-seq data

Regions of accessible chromatin consistently found in each cell type were identified by peak calling with macs2 (94). Cell type pseudobulk profiles were assembled by combining the fragments for all barcodes assigned to each cell type. Cell types with more than 100 million total fragments were randomly down sampled to 100 million fragments and then the macs2 (v2.2.7.1) callpeak tool was used to identify peaks with FDR less than 0.05. Duplicate peaks were retained. All cell type peaks were merged into an overall peak list using bedtools merge. ATAC fragments for each merged peak were quantified by barcode and used to make an snATAC-seq peaks-based counts matrix. Finally, all peaks were reduced to a fixed width of 500bps around the peak summit.

### Identification of genes and pathways differing between genotypes and cell types

Seurat’s findMarkers function was used to identify genes and accessible chromatin peaks that differed between cells within a cell type from each genotype (KPC and AKPC). A Wilcoxon rank sum test was used to compare the gene and peak counts from both groups and Bonferroni multiple test correction was performed to identify significant changes. For the differentially expressed genes from snRNA-seq data, gene set enrichment was performed using fGSEA (bioRxiv 2021 101101/060012) (v1.28.0). First the differential expression rank of each gene was calculated by multiplying the negative log10 p-value of the difference by the log2 fold change. All genes with a rank of 0 were assigned to a pseudocount of the minimum positive non-zero rank value. Then fGSEA was performed on the rank stats for all pathways with less than 500 genes from the following pathway sets from mSigDB (95) mm10 database: MH (Orthology-mapped hallmark gene sets), M5 GO (ontology gene sets, gene ontology), and M2 CP:Reactome (Curated gene sets, Realtime gene sets). All pathways with an Benjamini Hochberg adjusted p-value less than 0.1 were considered significantly different between genotypes. To assess the enrichment of predefined gene signatures in different cell types, Seurat’s AddModuleScore function was used.

### Creation of genotype-specific CAF subtype trajectories using gene expression

The monocle3 package (96) (v1.3.4) was used to perform gene expression-based trajectory analysis of CAF subtypes for both the KPC and AKPC genotypes, separately. For each genotype, a cell data set object was constructed using the raw RNA counts from all CAF cells, then the top 100 dimensions from PCA were used to reduce the dimensionality using the UMAP method with 10 nearest neighbors and a minimum distance of 0.01. Cells were clustered using the Louvain algorithm, 40 nearest neighbors and a partition q-value of 0.01. Finally, the trajectory was learned by using monocle’s learn_graph function, with an additional loop-closing step, only learning a single graph across all partitions and with up to 3 layers of nearest neighbor trees.

### Prediction of cell-cell signaling interactions

CellChat (40) (v1.6.1) was used to predict cell-cell signaling interactions between all cell types from scRNA-seq counts data of known genes for ligands and receptors. CellChat was run on each library separately using the CellChatDB mouse reference of known ligand and receptor signaling pairs involved in secreted signaling. The probability of interactions between cell types was calculated using average cell type gene expression, with bootstrapping to calculate p-values for each predicted interaction. After filtering out interactions between small numbers of cells, communication is aggregated into signaling pathways and summarized. Next, to compare signaling interactions between the two genotype libraries, the two CellChat objects were merged. The number of interactions and overall interaction strength per pathway were compared, as well as the amount of incoming and outgoing signals per cell type and genotype. Finally, latent communication patterns of coordinated pathways across cell types were calculated using CellChat’s non-negative matrix factorization method.

### Integrated analysis of transcription factor motif accessibility and target gene expression

Transcription factor (TF)-target regulatory relationships were obtained from TRRUST 2.0 (97) and ORegAnno 3.0 (98) databases. Mouse TFs with at least seven target gene counts (activated, repressed or both) were selected for the analysis. Position weight matrices of TF-binding motifs were obtained from the vertebrate CORE collection of JASPAR2020 database (99). Enrichment scores of motifs in accessible regions were calculated for each cell using chromVAR (100) through the Signac function RunChromVAR() (89).

Differential accessible TF-binding motifs within each cell type from both AKPC and KPC genotypes were identified using the Wilcoxon Rank Sum test implemented in Seurat’s FindAllMarkers() function with Bonferroni multiple test correction. All motifs with an adjusted p-value smaller than 0.05 were considered significantly enriched. Similarly, Seurat’s FindAllMarkers() function was used with a Wilcoxon Rank Sum test to identify differentially expressed genes between cell types from both genotypes. The differential expression rank for each gene was calculated by multiplying the negative log10 p-value by the sign of the log2 fold change.

To identify TFs displaying both significant motif and target gene enrichment in a specific cell type, readout for predicted TF activity, GSEA was performed on the target genes of the transcription factors that were enriched in a cell type from either AKPC or KPC or both genotypes using GSEA in pre-ranked mode (95). Statistical significance for target gene enrichment was set at a p-value cutoff of 0.05. Running enrichment scores for leading edge genes were normalized by gene set size.

### Bulk RNA sequencing

Library preparation for bulk 3’-sequencing of poly(A)-RNA was done as described previously (101). Briefly, barcoded cDNA of each sample was generated with a Maxima RT polymerase (Thermo Fisher Scientific) using oligo-dT primer containing barcodes, unique molecular identifiers (UMIs) and an adapter. 5’ ends of the cDNAs were extended by a template switch oligo (TSO); after pooling of all samples, full-length cDNA was amplified with primers binding to the TSO-site and the adapter. cDNA was tagmented with the Nextera XT kit (Illumina) and 3’-end-fragments finally amplified using primers with Illumina P5 and P7 overhangs. In comparison to Parekh et al. (101), the P5 and P7 sites were exchanged to allow sequencing of the cDNA in read1 and barcodes and UMIs in read2 to achieve a better cluster recognition. The library was sequenced on a NextSeq 500 (Illumina) with 75 cycles for the cDNA in read1 and 16 cycles for the barcodes and UMIs in read2. Gencode gene annotations version M18 and the mouse reference genome major release GRCm38 were derived from the Gencode homepage (https://www.gencodegenes.org/). Dropseq tools v1.12 (102) was used for mapping the raw sequencing data to the reference genome. The resulting UMI filtered countmatrix was imported into R v3.4.4. Prior differential expression analysis with DESeq2 1.18. (103), dispersion of the data was estimated with a parametric fit including the treatment the cell lines underwent as explanatory variable in the model. The Wald test was used for determining differentially regulated genes between the treated and non-treated group and shrunken log_2_ fold changes were calculated afterward, with setting the type argument of the lfcShrink function to normal. A gene was determined to be differentially regulated if the absolute log_2_ fold change was greater than 0.848 or lower than -0.848 and the adjusted *P* value was below 0.05. Gene set enrichment analyses were performed using R (The R Project for Statistical Computing; version v3.4.4).

### Proteomics sample preparation

Tumor cells were seeded in DMEM, containing 10% FBS and P/S, and propagated until reaching 70% confluency. The medium was replaced with serum-free DMEM containing P/S, and the cells were propagated for an additional 24 h before cell harvesting. Cell pellets were washed using cold 1× PBS and spun down at 4°C. Cell pellets were snap-frozen using liquid nitrogen prior -80°C storage and processing. Cell pellets were lysed in 8 M urea, 100 mM NH_4_HCO_3_, 1 mM DTT, pH 8.0. The total protein content of the cell lysate was determined using the BCA Protein Assay Kit (Thermo Scientific). Two hundred µg of total protein per sample were reduced (10 mM DTT, 30 min at 30°C), alkylated (55 mM chloroacetamide, room temperature, 30 min, in the dark) and diluted to 1.3 M urea with 50 mM NH_4_HCO_3_. Protein digestion was performed by adding trypsin (Trypsin Gold mass spectrometry grade, Promega, 1:50 enzyme-to-substrate ratio) and incubation overnight at 37°C. Digests were acidified with 1% formic acid (FA) and desalted using 50 mg tC18 reversed-phase (RP) solid-phase extraction cartridges (Sep-Pak, Waters, WAT054960). Peptide solutions were dried and frozen at -80°C until TMT-labeling.

### TMT labeling and offline peptide fractionation

In total, 10 samples were analyzed, including 5 biological replicates of KPC and AKPC cell lines. The 10 samples were measured in a TMT11-plex experiment, together with one pooled sample. TMT11-plex-labeling was carried out according to Zecha et al. (104). Briefly, peptide solutions (200 μg protein amount per sample) were reconstituted in 20 μL of 100 mM HEPES buffer (pH 8.5). Next, TMT11-plex reagents (Thermo Fisher Scientific) in 100% anhydrous acetonitrile were added in a final concentration of 11.6 mM to all peptide sample. After incubation for 1 h at 25°C and 400 rpm, the labeling reaction was stopped by adding 2 μL of 5% hydroxylamine (15 min, at 25°C and 400 rpm). The 11 peptide samples were pooled and dried down. Subsequently, they were desalted using 500 mg tC18, RP solid-phase extraction cartridges (Sep-Pak, Waters, WAT020805), dried and frozen at -80 °C. Finally, TMT11-plex experiment was off-line fractionated using basic reverse phase chromatography. Briefly, 150 µg of peptides were reconstituted in 25 mM ammonium bicarbonate (pH 8.0) and loaded onto a C18 column (XBridge BEH130, 3.5 mm, 2.1 3 150 mm, Waters Corp.). Peptides were eluted with increasing acetonitrile concentration to 96 fractions and subsequently pooled to 32 fractions. All fractions were dried down before LC-MS/MS analysis.

### LC-MS/MS measurements

Nano-flow LC-MS/MS measurements of the TMT-labeled peptides were performed on a Fusion Lumos Tribrid mass spectrometer (Thermo Fisher Scientific) equipped with an Ultimate 3000 RSLCnano system. For each analysis, peptides were delivered to a trap column (ReproSil-pur C18-AQ, 5 μm, Dr. Maisch, 20 mm × 75 μm, self-packed) at a flow rate of 5 μL/min in 100% solvent A (0.1% formic acid in HPLC grade water). After 10 min of loading, peptides were transferred to an analytical column (ReproSil Gold C18-AQ, 3 μm, Dr. Maisch, 450 mm × 75 μm, self-packed with self-pulled tip using a Laser-Pipette-Puller P-2000) and separated using a 50 min gradient from 8% to 32% of solvent B (0.1% formic acid in acetonitrile and 5% (v/v) DMSO) at 300 nL/min flow rate. Both nanoLC solvents contained 5% DMSO (v/v). The Fusion Lumos was operated in data dependent acquisition (DDA), positive ionization and multi-notch MS3 mode (105). Full scan MS1 spectra were recorded over a range of 360-1500 m/z at a resolution of 60k using an automatic gain control (AGC) target value of 4e5 and maximum injection time (maxIT) of 50 ms. Precursor ions were filtered according to charge state (2–6), dynamic exclusion (60 s with a ± 10 ppm window), and monoisotopic precursor selection. MS2 spectra were recorded in the orbitrap at 30K resolution and with a quadrupole isolation window of 0.7 m/z (scan range mode: Auto:m/z Normal). Fragmentation was performed with collision induced dissociation (CID) and a normalized collision energy (NCE) of 35% (AGC target value 5e4, maximal injection time 60 ms). For TMT reporter ion quantification an additional MS3 spectrum was acquired in the orbitrap over an m/z range of 100-1000 m/z at 50k resolution. For this, fragment ions were selected by multi-notch isolation, allowing a maximum of 10 notches. MS1 precursor isolation was set to 0.7 m/z, while the MS2 isolation window width was set to 2 m/z. Fragmentation in the orbitrap was performed by HCD at 55% NCE (AGC target value 1e5, maximal injection time 120 ms). The overall cycle time of the method was 3 s.

### MaxQuant database searching

Peptide identification and quantification were performed using MaxQuant (106) (version 1.6.17.0) with its built-in search engine Andromeda (107). Raw files were searched against the UniProt mouse reference database (downloaded from UniProt July 2020, 55,462 protein entries) supplemented with common contaminants. MS3-based TMT quantification was enabled. For all searches, carbamidomethylated cysteine was set as fixed modification and oxidation of methionine and N-terminal protein acetylation as variable modifications. Trypsin/P was specified as the proteolytic enzyme with up to 2 missed cleavage sites. Precursor tolerance was set to ± 4.5 ppm, and fragment ion tolerance to ± 20 ppm. Results were adjusted to a 1% false discovery rate (FDR) on peptide spectrum match (PSM) and protein level employing a target-decoy approach using reversed protein sequences.

### Proteomic data normalization and statistical analysis

Before statistical analysis, we first log_10_-transformed and median-centered the protein intensity in each TMT channel, to remove the potential influence resulting from different sample loading amounts. Next, we used Student’s t-test to identify proteins that were significantly differentially expressed between experimental conditions. All analyses were performed using R (version 4.1.2).

### Secretome analysis on antibody microarrays

Antibody microarrays were produced and used as described (108). Briefly, antibodies were spotted onto epoxy-coated slides (Nexterion-E; Schott) using a MicroGrid-2 contact printer (BioRobotics) with SMP3B pins (Telechem). A common protein reference sample made from 24 pancreatic cancer cell lines (109) was incubated on a microarray in parallel. The reference provided information about experimental performance and allowed direct data normalization. Protein concentration of secretome and reference samples was adjusted to 4.0 mg/mL. Labelling was conducted using the fluorescent dyes DY649 or DY549 (Dyomics), respectively, at a molar protein/dye ratio of 7.5 in PBS (pH 7.2) at 4°C for 2 h. Subsequently, 10% glycine in PBS was added to quench unreacted dye. For analyses, the microarray surface was blocked with 10% non-fat dry milk in TBS, 0.05% Tween 80 for 3 h at room temperature. The blocked slides were incubated with 25 µg/mL each of labeled secretome sample and common reference at 4°C overnight. After washing, a PowerScanner system (Tecan) was used for image capture at constant laser power and photo-multiplier tube gain. Image analysis was performed with GenePix Pro 6.0 software (Molecular Devices) generating numerical values of signal intensities. The data was analyzed using the R software package (version 4.0.4). The Loess method of the LIMMA package was applied for data normalization with background correction offset (v. 3.50.1) (110); variations were corrected by applying the functions ‘normalizeWithinArrays’, and ‘normalizeBetweenArrays’. Empirical Bayes test with Bonferroni-Hochberg adjustment was used for multiple testing (111).

### Protein term enrichment analysis

Protein term enrichment analyses were conducted using the annotation tool Database for Annotation, Visualization, and Integrated Discovery (DAVID) (https://david.ncifcrf.gov/home.jsp) with the BioCarta, Gene Ontology, KEGG, Reactome, and UniProt datasets. Significant enrichments are defined with *P* < 0.05.

### Orthotopic assay

For syngeneic orthotopic transplantations, KC, AKC, KPC, and AKPC mouse tumor cells were implanted by an injection of 25 × 10^3^ cells in 100 µL of 1:1 serum-free DMEM:Matrigel GFR into the pancreas of eight-week-old male C57BL/6J mice (n=5 per condition). Tumor take was ≥ 72% for each cell line. Overall survival was calculated as the time elapsed between orthotopic transplantation and euthanasia at a predefined ethical endpoint. For xenogeneic orthotopic transplantations, CRISPR-edited hPANC cells and PDOs were implanted by an injection of 25 × 10^3^ cells in 100 µL of 1:1 serum-free DMEM:Matrigel GFR into the pancreas of eight-week-old female Hsd:Athymic Nude-*Foxn1^nu^* mice (n=6 per condition). Tumor take was ≥ 90% for all models. Overall survival was calculated as the time elapsed between treatment/enrolment start and euthanasia at a predefined ethical endpoint. The solutions of galunisertib (TGFBR1 inhibitor, 12.5 mg/kg) and FOLFIRINOX (leucovorin, 50.0 mg/kg; 5-fluorouracil, 25.0 mg/kg; irinotecan, 25.0 mg/kg; oxaliplatin, 2.5 mg/kg) were administrated by i.p. injection. Briefly, the treatment administration schedule was the following: i.p. FOLFIRINOX injections every third day and i.p. galunisertib injections every second day. Tumors were then resected and fixed in cold 4% formaldehyde for 24 h and embedded in paraffin for histological analysis.

### Subcutaneous assay

For subcutaneous transplantation, tumor cell were implanted by subcutaneous injection of 4 × 10^5^ KPC or AKPC tumor cells and 4 × 10^5^ PSCs in 100 μl of 1:1 serum-free DMEM:Matrigel GFR into the flank of seven-week-old female Hsd:Athymic Nude-*Foxn1^nu^* mice (n=8 tumors per condition). Tumor take was 100% for all cell lines. The solutions of galunisertib (TGFBR1 inhibitor, 12.5 mg/kg) and MRTX1133 (Kras^G12D^ inhibitor, 3 mg/kg) were administrated by i.p. injection. Briefly, the treatment administration schedule was the following: MRTX1133 i.p. injections twice a day and i.p. galunisertib injections every second day. Tumors were then resected and fixed in cold 4% formaldehyde for 24 h and embedded in paraffin for histological analysis.

### Patient-derived xenografts

For subcutaneous implantation, 4 mm^3^ pieces of patient-derived xenografts (patient-derived *in vivo* expanded pancreatic tumors), were embedded in ECM Gel from Engelbreth-Holm-Swarm murine sarcoma (Sigma-Aldrich) and transplanted into the flank of six to eight-week-old female Hsd:Athymic Nude-*Foxn1^nu^* mice. Tumor take was 100% for all patient-derived xenografts. Tumor size was measured thrice a week with a caliper, and the tumor volumes were determined with the following equation: *v* = (*l* × *w*^2^) × π / 6 (where *v* is volume, *l* is length, and *w* is width). For each group, 8 patient-derived xenografts were implanted. Data are represented as mean ± SEM. Tumors were randomized when the volume reached 50 to 100 mm^3^ before starting treatment. The solutions of galunisertib (TGFBR1 inhibitor, 12.5 mg/kg) and FOLFIRINOX (leucovorin, 50.0 mg/kg; 5-fluorouracil, 25.0 mg/kg; irinotecan, 25.0 mg/kg; oxaliplatin, 2.5 mg/kg) were administrated by i.p. injection. Briefly, the treatment administration schedule was the following: i.p. FOLFIRINOX injections every third day and i.p. galunisertib injections every second day. Mice were euthanized when tumors reached a maximal volume of 1,000 mm^3^. Tumors were then resected and fixed in cold 4% formaldehyde for 24 h and embedded in paraffin for histological analysis.

### Magnetic resonance imaging

Magnetic resonance imaging (MRI) was performed to assess tumor growth using a dedicated ultrahigh field 11.7T small animal system (BioSpec 117/16, Bruker Biospin) equipped with a 9 cm gradient insert (BGA-S9). MRI data were acquired using ParaVision 6.01 software with a four-channel receive-only surface coil placed anterior to the tumor area. Anesthesia was maintained using 1.5% isoflurane (Abbvie) and adjusted to maintain a safe respiration rate of about 60 cycles per minute. The imaging protocol comprised a multi-slice 2D RARE technique in coronal and axial slice orientation with acquisition parameters as: TE/TR = 23 ms/1500 ms, spatial resolution Δr = 90×90×500 µm^3^, and RARE factor = 8. All MR images were analyzed using the ITK-SNAP software tool, which allowed for accurate manual segmentation of the tumors in 3D.

### Analysis of publicly available databases

RNA-seq and SNV data from the TCGA-PAAD cohort were obtained using TCGAbiolinks (112,113). Mutual co-occurrence and exclusivity of selected DDR gene mutations were assessed using Fisheŕs exact test. Bayesian cell type and gene expression deconvolution of the TCGA bulk RNA-seq data was performed using BayesPrism (59) with the Peng *et al.* dataset (60) as single cell reference. Raw single cell count matrices and cell type annotations were obtained from GSA accession CRA001160. Doublet barcodes were marked using scDblFinder (114) and removed from the analysis. Additional quality control filters were percent mitochondrial reads < 30% and unique RNA genes between the 2.5th and 97.5th percentiles. scRNA-seq counts were log-normalized and scaled, regressing out percent mitochondrial reads and cell cycle scores. Linear dimensionality reduction was performed using the top 3000 variable features, followed by integration with Harmony (93), and projection into UMAP space using the top 21 harmony-corrected principal components. Clusters were identified using the Louvain algorithm with 25 k-nearest neighbors at a resolution of 0.18. Cell type identities were assigned to each cluster by cross-referencing with the annotations from the original study (60). For annotation of CAFs, fibroblasts and stellate cells were subset and re-analyzed using the same workflow with 2000 variable features, 25 harmony-corrected PCs, 10 k-nearest neighbors and a clustering resolution of 0.2. Fibroblasts were annotated as myCAF, iCAF and non-CAF according to previously described marker gene signatures (17,115). BayesPrism deconvolution (59) was run on protein-coding genes using default parameters, treating major cell type annotations as cell types and CAF annotations as cell states. Validation of the deconvolution approach was performed by using pseudobulk gene expression profiles as bulk input and correlating estimated and true cell type and cell state abundances. Deconvoluted cell type-specific gene expression from the TCGA RNA-seq data was normalized using variance stabilizing transformation (103). Gene signature enrichment was calculated through gene set variation analysis (GSVA) (116). Correlations between malignant gene expression levels and cell type/state abundances and GSVA enrichment scores were computed using Spearman and Pearson correlation analysis, respectively.

### Statistical analysis

GraphPad Prism software was used for statistical analysis and graphical representation of the data. Statistical significances were tested using unpaired Student’s t-test. For survival comparisons, statistical significances were tested using log-rank (Mantel-Cox) test. Correlation analyses were conducted using Pearson’s correlation test. For comparing human PDAC case numbers, contingency tables were generated, and statistical significances were tested using Fisher’s exact test. All tests were considered to be statistically significant when *P* < 0.05.

## SUPPLEMENTARY FIGURE LEGENDS

**Supplementary Figure S1.** ATM loss-of-function promotes mesenchymal phenotype and DNA damage accumulation in pancreatic cancer cells, in a P53-deficient context. **A,** Kaplan-Meier analysis of survival of *Atm^+/+^*; *Trp53^fl/fl^*; *LSL-Kras^G12D/+^*; *Ptf1a^Cre/+^* (KPC) (n=21), *Atm^fl/+^*; *Trp53^fl/fl^*; *LSL-Kras^G12D/+^*; *Ptf1a^Cre/+^* (A^het^KPC), and *Atm^fl/fl^*; *Trp53^fl/fl^*; *LSL-Kras^G12D/+^*; *Ptf1a^Cre/+^* (AKPC) (n=30) mice. **, *P* < 0.01, log-rank (Mantel-Cox) test. Median survival of KPC, A^het^KPC, and AKPC mice are shown. **B,** Immunohistochemistry staining for CDH1 on KPC and AKPC endpoint pancreatic tumors. Scale bars, 50 µm. **C,** Representative brightfield images of KPC and AKPC cells. Scale bars, 100 µm. **D**, Bar graph showing the cell viability of KC, AKC, KPC and AKPC tumors cells cultured for 72h. Results show means ± SD of at least 3 independent experiments where two cell lines per genotype were tested. Each dot represents an independent experiment. **E–G,** Immunofluorescence staining for H2AX p-S139 (green) (**E**) and quantification of nucleus size (**F**) and H2AX p-S139^+^ nucleus (**G**) in KPC and AKPC tumor cells. Cells were counterstained with DAPI (blue). Scale bars, 10 µm. Results show means ± SD of 3 KPC and 2 AKPC cell lines used in at least 2 independent experiments and a minimum of 140 nucleus/cell line were analyzed (**F**) and means ± SD of 3 cell lines used in at least 2 independent experiments (**G**). **, *P* < 0.01; ****, *P* < 0.0001, unpaired Student *t* test. ns, not significant.

**Supplementary Figure S2.** HRD ATM-deleted PDAC exhibits a unique immune and myCAF stroma. **A** and **B,** Immunohistochemistry staining for CD3ε, CD8, and CD206 and immunofluorescence staining for CD11b (white) and B220 (red), and Ly6c (red) on *Atm^+/+^*; *LSL-Kras^G12D/+^*; *Ptf1a^Cre/+^* (KC) and *Atm^fl/fl^*; *LSL-Kras^G12D/+^*; *Ptf1a^Cre/+^* (AKC), *Atm^+/+^*; *Trp53^fl/fl^*; *LSL-Kras^G12D/+^*; *Ptf1a^Cre/+^* (KPC) and *Atm^fl/fl^*; *Trp53^fl/fl^*; *LSL-Kras^G12D/+^*; *Ptf1a^Cre/+^*(AKPC) endpoint pancreatic tumors (**A**) and quantification of CD3ε^+^, CD8^+^, CD206^+^, CD11b^+^, B220^+^, CD11b^+^B220^+^, Ly6c^+^, and CD11b^+^Ly6c^+^ cells. Scale bars, 50 µm. Results show means ± SD of ≥ 3 biological replicates. Each dot represents one mouse. *, *P* < 0.05, unpaired Student *t* test. **C,** Histologic sections stained by picrosirius red (left) and immunofluorescence staining for αSMA (green) and SMAD2 p-S465/S467 (red) in KC and AKC endpoint pancreatic tumors. Cells were counterstained with DAPI (blue). Scale bars, 100 µm. **D** and **E,** Histologic sections stained by picrosirius red and imaged using polarized light (PL) microscopy (**D**) and quantification of type I (red) vs. type III (green) collagen fibers (top) and mean maximum relative (max. rel.) peak frequency of collagen fibers directionality (bottom) in KPC and AKPC endpoint pancreatic tumors (**E**). Scale bars, 100 µm. Results show means ± SD of ≥ 3 biological replicates (**E** top) and are represented as Tukey box plots of 4 biological replicates (**E** bottom). Each dot represents one mouse. *, *P* < 0.05, unpaired Student *t* test. **F,** Immunofluorescence staining for αSMA (green) on *Atm^fl/+^*; *Trp53^fl/fl^*; *LSL-Kras^G12D/+^*; *Ptf1a^Cre/+^* (A^het^KPC) endpoint pancreatic tumors (top) and quantification of αSMA^+^ surface in KPC, A^het^KPC, and, AKPC tumors. Scale bars, 50 µm. Results show means ± SD of ≥ 4 biological replicates. Each dot represents one mouse. *, *P* < 0.05, unpaired Student *t* test.

**Supplementary Figure S3.** Single nucleus multiomics analysis and immune cell composition in KPC and AKPC tumors. **A,** UMAP embedding visualizing snRNA-seq (top left), snATAC-seq (bottom left) and integrated snRNA/ATAC-seq (top right) clusters identified in *Atm^+/+^*; *Trp53^fl/fl^*; *LSL-Kras^G12D/+^*; *Ptf1a^Cre/+^* (KPC) and *Atm^fl/fl^*; *Trp53^fl/fl^*; *LSL-Kras^G12D/+^*; *Ptf1a^Cre/+^* (AKPC) endpoint pancreatic tumors. **B** and **C,** Distribution of RNA (**B** top) and ATAC (**C** top) counts, RNA percentage of mitochondrial counts (**B** bottom) and ATAC mitochondrial fragments (**C** bottom) detected in each cluster identified in KPC and AKPC tumors. **D,** Dot plot depicting highest expressed genes segregating the cell clusters identified in KPC and AKPC tumors. Color intensity of dot plot indicates the average expression for the corresponding gene. Dot size corresponds to the proportion of cells with non-zero expression of the gene in each cell cluster. **E** and **F,** UMAP embedding visualizing integrated snRNA-seq and ATAC-seq data of cell clusters identified in KPC (top) and AKPC (bottom) (**E**) and comparisons of RNA counts, RNA percentage of mitochondrial (MT) counts (top), ATAC counts and ATAC MT fragments (bottom) between AKPC and KPC tumors (**F**). **G,** Violin plots depicting dissociation (top) and hypoxia (bottom) signature expression levels per cell type in KPC and AKPC tumors. **H** and **I**, Immune cell subsets characterization of lymphocytes (**H**) and myeloid cells (**I**) identified with the snRNA/ATAC-sequencing analysis of KPC and AKPC tumors. UMAP embedding visualizing integrated snRNA/ATAC-seq (top left) clusters of lymphocytes (**H**) and myeloid cells (**I**) identified in KPC and AKPC tumors. Dot plot showing the genes segregating the different identified immune cell subtypes (right). Proportion of cell subtypes per total cells and lymphocytes (**H**) or myeloid cells (**I**) in each genotype are depicted (bottom left). UMAP, Uniform manifold approximation and projection. TAMs, tumor-associated macrophages.

**Supplementary Figure S4.** Molecular profiling of the cancer-associated fibroblast clusters identified with the single-nucleus multiomics analysis of KPC and AKPC tumors. **A,** Dot plot depicting expression of genes from the myofibroblastic, inflammatory and antigen-presenting cancer-associated fibroblasts (respectively, myCAF, iCAF and apCAF) signatures in myCAF, iCAF and apCAF clusters from *Atm^+/+^*; *Trp53^fl/fl^*; *LSL-Kras^G12D/+^*; *Ptf1a^Cre/+^* (KPC; n=2) and *Atm^fl/fl^*; *Trp53^fl/fl^*; *LSL-Kras^G12D/+^*; *Ptf1a^Cre/+^* (AKPC; n=2) endpoint tumors. Color intensity of dot plot indicates the average expression for the corresponding gene. Dot size corresponds to the proportion of cells with non-zero expression of the gene in each cell cluster. **B,** Expression levels of *ACTA2* in KPC and AKPC tumor cell clusters, represented as UMAP plot. **C,** GO, REACTOME and Hallmark-based gene-set enrichment analysis of the transcriptomics data showing upregulated and downregulated pathways with *P* ≤ 0.05 in AKPC vs. KPC fibroblasts. **D,** UMAP embedding visualizing the inferred positioning of myCAFs, iCAFs, and apCAFs along transcriptomic-based single-cell trajectory. **E,** Gene set enrichment analysis performed on the target genes of the transcription factors (TF) enriched in myofibroblastic myCAFs (top) or iCAFs (down) from KPC and AKPC tumors. UMAP, Uniform manifold approximation and projection.

**Supplementary Figure S5.** HRD ATM-deleted PDAC cell secretome drives the differentiation of pancreatic stellate cells into myCAF. **A,** Schematic representation of the pancreatic stellate cells (PSCs) culture on GFR Matrigel-coated plates used for assessing CAF differentiation in co-culture and conditioned medium (CM)-based approaches (top), immunofluorescence staining for αSMA (bottom left), quantification of αSMA^+^ cells, and qPCR analysis of *ACTA2* expression (bottom right) on PSCs cultured as 2D monolayer or on GFR Matrigel coating for 72 hours. Cells were counterstained with DAPI. Scale bars, 20 µm. Results show means ± SD of ≥ 2 (IF) or 3 (qPCR) independent experiments. Each dot represents an independent experiment. *, *P* < 0.05; **, *P* < 0.01, unpaired Student *t* test. **B,** qPCR analysis of inflammatory cancer-associated fibroblast (iCAF) and myofibroblastic (myCAF) marker gene expression in PSCs cultured on GFR Matrigel coating and treated with 10 ng/mL TGF-β1 (myCAF inducer) or 1 ng/mL IL1α (iCAF inducer) for 72 hours. Results show means ± SD of ≥ 3 independent experiments. Each dot represents an independent experiment. *, *P* < 0.05; **, *P* < 0.01; ***, *P* < 0.001, unpaired Student *t* test. **C** and **D,** Immunofluorescence staining for αSMA (**C** left) and SMAD2 p-S465/S467 (**D** left), and quantification of αSMA^+^ (**C** right) and SMAD2 p-S465/S467^+^ PSCs (**D** right) cultured on GFR Matrigel coating and treated or not with 10 ng/mL TGF-β1 for 72 hours. Cells were counterstained with DAPI. Scale bars, 20 µm. Results show means ± SD of ≥ 4 independent experiments. Each dot represents an independent experiment. *, *P* < 0.05, unpaired Student *t* test. **E,** qPCR analysis of myCAF and iCAF marker gene expression in PSCs co-cultured for 72 hours with KC (top) or AKC (bottom) tumor cells as in (Figure 2J). Results show means ± SD of ≥ 2 cell lines used in at least 2 independent experiments. Each dot represents a cell line. *, *P* < 0.05; **, *P* < 0.01, unpaired Student *t* test. **F,** Immunofluorescence staining for αSMA (green) and SMAD2 p-S465/S467 (red) on PSCs co-cultured for 72 hours with KC, AKC, KPC, or AKPC tumor cells (left) as in (Figures 2J and **2K**), and quantification of αSMA^+^ and SMAD2 p-S465/S467^+^ PSCs vs. in inactivated PSCs (right). Cells were counterstained with DAPI (blue). Scale bars, 20 µm. **G,** Schematic representation of the CM-based approach used to assess the ability of tumor cell secretome to differentiate PSCs into cancer-associated fibroblasts (CAFs) (**H**). **H,** Immunofluorescence staining for αSMA (green) and SMAD2 p-S465/S467 (red) on PSCS cultured with KC_CM_, AKC_CM_, KPC_CM_, and AKPC_CM_ for 72 hours (left), and quantification of αSMA^+^ and SMAD2 p-S465/S467^+^ PSCs (right). Scale bars, 20 µm. Results show means ± SD of ≥ 5 CM batches. Each dot represents a CM batch. CM from at least 2 different cell lines per genotype were tested. *, *P* < 0.05; **, *P* < 0.01, unpaired Student *t* test. **I** and **J,** Immunofluorescence staining for PDPN (white), CXCL2 (green), and POSTN (red) (top) and PDPN (white), Ly6c (green), and CTGF (red) (bottom) on KC, AKC, KPC, and AKPC endpoint pancreatic tumors (**I**), and quantification of PDPN^+^CTGF^+^, PDPN^+^POSTN^+^, PDPN^+^CXCL2^+^, and PDPN^+^Ly6c^+^ cells (**J**). Cells were counterstained with DAPI (blue). Scale bars, 100 µm. Results show means ± SD of ≥ 3 KPC and AKPC mice and ≥ 2 KC and AKC mice. Each dot represents a mouse. *, *P* < 0.05; **, *P* < 0.01, unpaired Student *t* test. **K** and **L,** Schematic representation of the porcine urinary bladder-based *ex vivo* co-culture assay (top), immunofluorescence staining for CK19 (green), αSMA (green), and SMAD2 p-S465/S467 (red) on resected co-culture grafts involving PSCs and KC, AKC, KPC, or AKPC tumor cells for 5 days (**K**), and quantification of αSMA^+^ surface (top) and of αSMA^+^SMAD2 p-S465/S467^+^ PSCs (bottom) (**L**) from the *ex vivo* co-culture assay shown in (**K**). Cells were counterstained with DAPI (blue). Scale bars, 50 µm. Results show means ± SD of ≥ 3 independent experiments. At least 2 cell lines per genotype were tested. Each dot represents an independent experiment. *, *P* < 0.05; **, *P* < 0.01, unpaired Student *t* test. **M** and **N,** Immunofluorescence staining for αSMA (green) and SMAD2 p-S465/S467 (red) on PSCs co-cultured for 72 hours with *ATM^+/+^*or *ATM^+/Δ^* CRISPRed human MIA PaCa-2 tumor cells (**M**), and quantifications of αSMA^+^ and SMAD2 p-S465/S467^+^ PSCs (**N**). Cells were counterstained with DAPI (blue). Scale bars, 20 µm. Results show means ± SD of 2 cell lines, each used in 3 independent experiments. Each dot represents a cell line. *, *P* < 0.05, unpaired Student *t* test. FC, fold change; PUB, porcine urinary bladder; Veh., vehicle.

**Supplementary Figure S6.** Transcriptomic and proteomic profiling of ATM-deficient tumor cells reveals an actin cytoskeleton remodeling and mesenchymal phenotype. **A,** Gene set enrichment analysis performed on the target genes of the transcription factors (TF) enriched in ductal (top) and EMT-like (down) cells from *Atm^+/+^*; *Trp53^fl/fl^*; *LSL-Kras^G12D/+^*; *Ptf1a^Cre/+^*(KPC; n=2) and *Atm^fl/fl^*; *Trp53^fl/fl^*; *LSL-Kras^G12D/+^*; *Ptf1a^Cre/+^* (AKPC; n=2) endpoint tumors. Highlighted are transcription factors with both significantly enriched motif accessibility (adjusted *P* < 0.05) and target gene enrichment (*P* < 0.05). **B,** Heatmap showing the normalized enriched scores for core enriched target genes of transcription factors with differentially predicted activity in ductal cells from KPC and AKPC tumors (left) and representative REACTOME and GO:BP functional annotations for core enriched target genes upregulated in KPC (top) or AKPC (bottom) ductal cells. **C,** Representative KEGG, KW, and GO:CC functional annotations for proteins upregulated (left) and downregulated (right) with *P* ≤ 0.05 in AKPC (n=5) vs. KPC (n=5) cells. **D,** Protein abundancies for epithelial (CDH1, TJP1, TJP2, and OCLN) and mesenchymal (VIM, FN1 and ZEB1) protein markers in AKPC and KPC cells. Results are represented as Tukey box plots of 5 biological replicates. *, *P* < 0.05; **, *P* < 0.01, unpaired Student *t* test. KEGG, Kyoto encyclopedia of genes and genomes; KW, keywords.

**Supplementary Figure S7.** TGF-β signaling induced by ATM-deficient cancer cell drives myCAF differentiation. **A,** KEGG, GO, and BioCarta-based signaling pathway and biological process term enrichment analyses for proteins enriched in *Atm^+/+^*; *Trp53^fl/fl^*; *LSL-Kras^G12D/+^*; *Ptf1a^Cre/+^* (KPC; n=3) vs. *Atm^+/+^*; *LSL-Kras^G12D/+^*; *Ptf1a^Cre/+^*(KC; n=3) cell tumor cell secretomes. **B,** qPCR analysis of *Tgfb1* gene expression in KC and *Atm^fl/fl^*; *LSL-Kras^G12D/+^*; *Ptf1a^Cre/+^* (AKC) cells. Results show means ± SD of 2 cell lines, each used at least in 2 independent experiments. Unpaired Student *t* test. **C,** Western blot analysis of ATM in *sgATM* or *sgCTRL* CRISPRed hPANC tumor cells. **D,** Dot plot showing Tgfb1 gene expression in the cell clusters identified in KPC and AKPC tumors. Color intensity indicates the average expression for the corresponding gene. Dot size corresponds to the proportion of cells with non-zero expression of the gene in each cell cluster. **E,** Dot plot showing *Tgfb2, Tgfb3, Tgfbr1*, and *Tgfbr2* gene expression in the myofibroblastic CAFs (myCAF), inflammatory CAFs (iCAF), antigen-presenting CAFs (apCAFs), ductal, and EMT-like cells in KPC and AKPC tumors. Color intensity indicates the average expression for the corresponding gene. Dot size corresponds to the proportion of cells with non-zero expression of the gene in each cell cluster. **F,** Protein abundances for TGF-β1 ligand and TGFBR1 and TGFBR2 receptors in AKPC and KPC cells. Results are represented as Tukey box plots of 5 biological replicates. **G,** Schematic representation of the TGF-β reporter assay (left) and relative luciferase activity of HEK 293T cells transiently transfected with the SMAD-responsive (CAGA)_12_-Luc reporter construct and treated for 48 hours with KC, AKC, KPC, and AKPC tumor cell conditioned medium (CM) (right). Results show means ± SD of ≥ 2 cell lines. At least 3 CM batches per cell line were tested. Each dot represents a cell line. *, *P* < 0.05; **, *P* < 0.01, unpaired Student *t* test. **H,** Representative histologic sections stained by immunofluorescence staining for CK19 (white) and TGF-β1 (red) on KC and AKC endpoint pancreatic tumors. Cells were counterstained with DAPI (blue). Scale bars, 100 µm. **I,** Inferred outgoing communication patterns of secreting cells in KPC and AKPC tumors. **J,** Chord diagram showing the inferred ligand-receptor interactions between EMT-like cells and myCAF, iCAF subpopulations in KPC tumors. **K,** Heatmap showing mRNA expression of known TGF-β target genes regulated in fibroblasts (left) and SMAD3/4 target genes (right) in AKPC vs. KPC fibroblasts, myCAFs, iCAFs and apCAFs and ductal and EMT-like cells. Stars indicate significant difference between AKPC vs. KPC with an adjusted *P* < 0.05. **L,** Dot plot showing transcriptional expression of selected genes, involved in TGF-β bioavailability regulation, in the myCAF, iCAF, apCAF, ductal and EMT-like cells from KPC and AKPC tumors. Color intensity indicates the average expression for the corresponding gene. Dot size corresponds to the proportion of cells with non-zero expression of the gene in each cell cluster. **M,** Protein abundancies of selected proteins involved in TGF-β bioavailability regulation in KPC and AKPC tumor cells. Results are represented as Tukey box plots of 5 biological replicates. **N,** Immunofluorescence staining for αSMA (green) and SMAD2 p-S465/S467 (red) (left) and quantification of αSMA^+^ (top right) and SMAD2 p-S465/S467^+^ (bottom right) PSCs co-cultured for 72 hours with AKC or AKPC tumor cells and treated or not with 10 µM SB431542 (TβRi). Cells were counterstained with DAPI (blue). Scale bars, 20 µm. Results show means ± SD of 3 independent experiments. Representative images of vehicle conditions are shown in (**Supplementary Figure S5F**). At least 2 cell lines per genotype were tested. Each dot represents an independent experiment. *, *P* < 0.05; **, *P* < 0.01, unpaired Student *t* test. **O,** qPCR analysis of myCAF and iCAF marker gene expression in PSCs co-cultured for 72 hours with AKC (top) or AKPC (bottom) tumor cells as in (**N**). Results show means ± SD of ≥ 2 cell lines used in at least 2 independent experiments. Each dot represents a cell line. *, *P* < 0.05; ***, *P* < 0.001, unpaired Student *t* test. **P,** Immunofluorescence staining for αSMA (green) and SMAD2 p-S465/S467 (red) (left) and quantification of αSMA^+^ (top right) and SMAD2 p-S465/S467^+^ (bottom right) PSCs co-cultured for 72 hours with *sgATM* or *sgCTRL* CRISPRed hPANC tumor cells and treated or not with 10 µM SB431542. Cells were counterstained with DAPI (blue). Scale bars, 20 µm. Results show means ± SD of 3 cell lines tested in at least 3 independent experiments. Each dot represents a cell line. *, *P* < 0.05; **, *P* < 0.01; ***, *P* < 0.001, unpaired Student *t* test. **Q,** Schematic representation of the porcine urinary bladder-based *ex vivo* co-culture assay (top), immunofluorescence staining for αSMA (green) and SMAD2 p-S465/S467 (red) (bottom), and quantification of αSMA^+^ and of αSMA^+^SMAD2 p-S465/S467^+^ PSCs (right) on resected co-culture grafts involving PSCs and AKPC or AKC (vehicle conditions are shown in **Supplementary Figure S5K** and **S5L**) tumor cells for 8 days. Cells were counterstained with DAPI (blue). Scale bars, 100 µm. Results show means ± SD of 3 (AKPC) and 2 (AKC) independent experiments. Each dot represents an independent experiment. *, *P* < 0.05; **, *P* < 0.01; ***, *P* < 0.001, unpaired Student *t* test.

**Supplementary Figure S8.** TGF-β signaling triggered by ATM-deficient cancer cell drives myCAF differentiation in PSC- and non-PSC-based models. **A,** UMAP embedding visualizing distribution and density of BMP4 signature across cell populations identified from snRNA/ATAC-seq from *Trp53^fl/fl^*; *LSL-Kras^G12D/+^*; *Ptf1a^Cre/+^* (KPC; n=2) and *Atm^fl/fl^*; *Trp53^fl/fl^*; *LSL-Kras^G12D/+^*; *Ptf1a^Cre/+^*(AKPC; n=2) tumors. B, Violin plots depicting BMP4 signature expression levels across cell types identified from snRNA/ATAC-seq in KPC and AKPC tumors. **C,** qPCR analysis of myofibroblastic (myCAF) and inflammatory cancer-associated fibroblast (iCAF) marker gene expression on PSCs treated with 200 ng/ml BMP4 in combination or not with 200 µM of the ALK2/3/6 inhibitor (ALKi) LDN-193189 for 72 hours. Results show means ± SD of 3 independent experiments. Each dot represents an independent experiment. *, *P* < 0.05; **, *P* < 0.01, unpaired Student *t* test. **D,** qPCR analysis of myCAF and iCAF marker gene expression on PSCs co-cultured with AKC or AKPC tumor cells and treated or not with 200 µM LDN-193189 for 72 hours. Results show means ± SD of 2 cell lines, each used in 2 independent experiments. Each dot represents a cell line. *, *P* < 0.05, unpaired Student *t* test. **E,** Western blot analysis of TGF-β1 in *sgTgfb1* or *sgCtrl* CRISPRed AKC and AKPC tumor cells. **F,** qPCR analysis of myCAF and iCAF marker gene expression in PSCs co-cultured with *sgTgfb1* or *sgCtrl* CRISPRed AKC and AKPC tumor cells. Results show means ± SD of 2 cell lines, each used in at least 2 independent experiments. Each dot represents a cell line. *, *P* < 0.05; **, *P* < 0.01; ***, *P* < 0.001, unpaired Student *t* test. **G,** Violin plots depicting PSC and non-PSC signature expression levels in KPC and AKPC tumors (left) and in myCAFs, iCAFs and apCAFs (right) identified from snRNA/ATAC-seq in KPC and AKPC tumors. **H** and **I,** Immunofluorescence staining for αSMA (green) and SMAD2 p-S465/S467 (red) on CAFs co-cultured for 72 hours with KC, AKC, KPC, and AKPC tumor cells and treated or not with the TGFBR1 inhibitor SB431542 (TβRi; 10 µM) (**H**), and quantification of αSMA^+^ and SMAD2 p-S465/S467^+^ CAFs vs. in vehicle-treated CAFs (**I**). Cells were counterstained with DAPI (blue). Scale bars, 20 µm. Results show means ± SD of 2 CAF lines. Two tumor cell lines per genotype were tested in at least two independent experiments. Each dot represents a CAF line. *, *P* < 0.05; **, *P* < 0.01, unpaired Student *t* test. **J,** qPCR analysis of myCAF and iCAF marker gene expression on CAFs co-cultured for 72 hours with KC, AKC, KPC, and AKPC tumor cells and treated or not with 10 µM SB431542. Results show means ± SD of 2 tumor cell lines, each used in co-culture experiments with 3 different CAF lines. Each dot represents a cell line. *, *P* < 0.05; **, *P* < 0.01, unpaired Student *t* test. **K,** Schematic illustration of the subcutaneous assay (left), immunohistochemistry staining for αSMA (middle), and quantification of αSMA^+^ staining (right) on resected tumors arising either from KPC or AKPC cells co-transplanted with PSCs and treated with the KRAS^G12D^ inhibitor MRTX1133 (3 mg/kg) in combination or not with the TGFBR1 inhibitor galunisertib (galu.; 12.5 mg/kg). Scale bars, 50 µm. Vehicle groups are shown in (Figure 4Q) and statistical comparisons of the MRTX1133-treated groups vs. the vehicle groups in AKPC and KPC tumors are depicted. Results show means ± SD. Each dot represents a subcutaneous tumor. **, *P* < 0.01; ****, *P* < 0.0001; unpaired Student *t* test. FC, fold change; ns, not significant; Veh., vehicle.

**Supplementary Figure S9.** ATM loss in pancreatic tumor cells correlates with ROS-mediated myCAF differentiation. **A,** Western blot analysis (top) and quantification (bottom) of ACTN4 in *Atm^+/+^*; *Trp53^+/+^*; *LSL-Kras^G12D/+^*; *Ptf1a^Cre/+^* (KC) and *Atm^fl/fl^*; *Trp53^+l+^*; *LSL-Kras^G12D/+^*; *Ptf1a^Cre/+^* (AKC) tumor cells. Results show means ± SD of 2 cell lines. Each dot represents a cell line. *, *P* < 0.05, unpaired Student *t* test. **B,** Immunofluorescence staining for ACTN4 (red) and PXN (green) on KC vs. AKC tumor cells (left) and quantifications of ACTN4^+^ and PXN^+^ cells (right). Cells were counterstained with DAPI (blue). Scale bars, 20 µm. Results show means ± SD of ≥ 2 independent experiments. At least 2 cell lines/genotype were tested. Each dot represents a cell line. *, *P* < 0.05; **, *P* < 0.01, unpaired Student *t* test. **C**, REACTOME and KEGG-based gene set enrichment analysis plots of oxidative phosphorylation and respiratory electron transport in *Atm^fl/fl^*; *Trp53^fl/fl^*; *LSL-Kras^G12D/+^*; *Ptf1a^Cre/+^*(AKPC; n=4) vs. *Atm^+/+^*; *Trp53^fl/fl^*; *LSL-Kras^G12D/+^*; *Ptf1a^Cre/+^* (KPC; n=5) cell lines. **D,** Levels of reactive oxygen species (ROS) in *sgATM* and *sgCTRL* CRISPRed hPANC. Results show means ± SD of 3 cell lines used in at least 3 independent experiments. Each dot represents a cell line. Unpaired Student *t* test. **E,** Levels of ROS in KC and KPC tumor cells treated or not for 24 hours with the ATM inhibitor (ATMi) AZD0156 (1 µM). Results show means ± SD of 3 cell lines used in at least 2 independent experiments. Each dot represents a cell line. *, *P* < 0.05, unpaired Student *t* test. **F** and **G,** Immunofluorescence staining for ACTN4 (red) and PXN (green) and direct fluorescence of cortical F-actin by phalloidin-Atto565 (purple) on KPC and KC tumor cells treated or not for 24 hours with 1 µM AZD0156 (**F**), and quantification of ACTN4^+^, PXN^+^, and F-actin^+^ staining in DAPI^+^ cells (**G**). Cells were counterstained with DAPI (blue). Scale bars, 10 µm. Results show means ± SD of 3 independent experiments. Two cell lines per genotype were tested. Each dot represents an independent experiment. *, *P* < 0.05; **, *P* < 0.01, unpaired Student *t* test. **H,** Levels of ROS in AKPC tumor cells treated or not with the ROS inhibitor (ROSi) YCG063 (2 µM) for 24 hours. Results show means ± SD of 2 cell lines used in at least 3 independent experiments. Each dot represents a cell line. *, *P* < 0.05, unpaired Student *t* test. **I,** Immunofluorescence staining for ACTN4 (red), and VINC (green) and PXN (red) and representative image of the microchannel single cell migration assay conducted with AKPC tumor cells treated with 2 µM YCG063 for 24 hours (top) and quantification of ACTN4^+^, PXN^+^, and F-actin^+^ staining in DAPI^+^ cells (bottom) compared to vehicle shown in (Figure 5C). Cells were counterstained with DAPI (blue). White asterisks show migrating tumor cells. For the immunofluorescence experiments: scale bars, 20 µm. Results show means ± SD of 3 independent experiments. At least 2 cell lines per genotype were tested. Each dot represents an independent experiment. *, *P* < 0.05; **, *P* < 0.01, unpaired Student *t* test. For the microchannel single cell migration assay: scale bar, 50 µm. Results show means ± SD of 2 cell lines per genotype were tested in 2 independent experiments. Each dot represents a cell line. Unpaired Student *t* test. **J,** Western blot analysis (top) and quantification (bottom) of ACTN4 in AKPC tumor cells. Results show means ± SD of 3 independent experiments. Two tumor lines were used. Each dot represents an independent experiment. Unpaired Student *t* test. **K,** Direct fluorescence of cortical F-actin by phalloidin-Atto565 (white) and immunofluorescence staining for ACTN4 (red) and PXN (green) on AKC tumor cells treated with 2 µM YCG063 for 24 hours (left) and quantification of ACTN4^+^ and PXN^+^ cells (right) compared to vehicle shown in (**B**). Cells were counterstained with DAPI (blue). Scale bars, 10 µm. Results show means ± SD of ≥ 2 independent experiments. Two cell lines per genotype were tested. Each dot represents a cell line. **, *P* < 0.01, unpaired Student *t* test. **L,** Representative brightfield images at 0 hour and after 48 hours (left) of AKC cells subjected to a wound healing assay and treated or not with the ROSi (2 µM) (left) and quantification of wound closure over time (right). Results show means ± SD of 3 independent experiments. Two cell lines per genotype were tested. Each dot represents a cell line. *, *P* < 0.05, unpaired Student *t* test. **M,** Levels of ROS in AKPC tumor cells treated or not with the oxidant menadione (2-methyl-1,4-naphthoquinone, 25 µM; menad.) for 6 hours. Results show means ± SD of 2 cell lines used in 2 independent experiments. Each dot represents a cell line. **, *P* < 0.01, unpaired Student *t* test. **N,** Immunofluorescence staining for ACTN4 (red) and PXN (green) (top) and quantification of ACTN4^+^, PXN^+^, and F-actin^+^ staining in DAPI^+^ cells compared to vehicle (bottom) in AKPC tumor cells treated or not with 25 µM menadione for 6 hours. Cells were counterstained with DAPI (blue). Scale bars, 20 µm. Results show means ± SD of 3 cell lines used in 2 independent experiments. Each dot represents a cell line. **, *P* < 0.01; ***, *P* < 0.001, unpaired Student *t* test. **O,** Immunofluorescence staining for αSMA (green) and SMAD2 p-S465/S467 (red) on PSCs co-cultured for 72 hours with *sgKeap1*, *sgActn4*, *sgPxn*, or *sgCtrl* CRISPRed AKC and AKPC tumor cells as in (Figures 5L and **5M**). **P,** qPCR analysis of myofibroblastic (myCAF) and inflammatory cancer-associated fibroblast (iCAF) marker gene expression in PSCs co-cultured for 72 hours with *sgKeap1*, *sgActn4*, *sgPxn*, or *sgCtrl* CRISPRed AKC and AKPC tumor cells. Results show means ± SD of 2 cell lines, each used in at least 2 independent experiments. Each dot represents a cell line. *, *P* < 0.05; ****, *P* < 0.0001, unpaired Student *t* test. **Q,** Schematic representation of the conditioned medium-based assay (**R** and **S**). **R** and **S,** Immunofluorescence staining for αSMA (green) and SMAD2 p-S465/S467 (red) on PSCs treated for 72 hours with conditioned medium (CM) from AKC and AKPC tumor cells treated or not with 2 µM YCG063 for 24 hours (**R**) as in (**Q**), and quantification of αSMA^+^ (center) and SMAD2 p-S465/S467^+^ PSCs compared to vehicle (**S**). Cells were counterstained with DAPI (blue). Scale bars, 10 µm. Results show means ± SD of 2 cell lines used for generating at 3 CM batches per condition (vehicle and ROSi). Each dot represents a cell line. **, *P* < 0.01, unpaired Student *t* test, not significant. FDR, false discovery rate; KEGG, Kyoto encyclopedia of genes and genomes; NES, normalized enrichment score.

**Supplementary Figure S10.** Perturbing tumor–CAF dialog through TGF-β signaling interference suppresses aggressiveness and chemoresistance in ATM-deficient PDAC. **A** and **B,** Colony formation assay on *Atm^+/+^*; *LSL-Kras^G12D/+^*; *Ptf1a^Cre/+^* (KC) and *Atm^fl/fl^*; *LSL-Kras^G12D/+^*; *Ptf1a^Cre/+^*(AKC) tumor cells co-cultured or not with pancreatic stellate cells (PSCs) in a Transwell-based system and treated or not with 6 µM oxaliplatin (OXA) (**A**) and quantification of tumor cell viability (**B**). Results show means ± SD of 3 independent experiments. At least 2 cell lines per genotype were tested. Each dot represents an independent experiment. *, *P* < 0.05; **, *P* < 0.01, unpaired Student *t* test. **C,** Immunohistochemistry staining for cleaved-caspase 3 (CC3) on resected co-culture grafts involving PSCs and KC or AKC tumor cells cultured on the porcine urinary bladder (PUB) *ex vivo* platform for 8 days and treated or not with the TGFBR1 inhibitor SB431542 (TβRi; 10 µM) and FOLFIRINOX (FX; 75 nM 5-fluorouracil, 25 nM irinotecan, and 5 nM oxaliplatin) from the PUB *ex vivo* assay shown in (Figure 6C). Scale bars, 100 µm. **D,** Quantification of CC3^+^ surface. Results show means ± SD of ≥ 2 independent experiments. At least 2 cell lines per genotype were tested. Each dot represents an independent experiment. *, *P* < 0.05, unpaired Student *t* test. **E,** Schematic illustration of the PUB used as an *ex vivo* system for assessing tumor cell invasion. **F–I,** Immunohistochemistry staining for CK19 on resected co-culture grafts involving PSCs and KPC or AKPC (**F**), and KC or AKC (**H**) tumor cells shown in (Figure 6C and **6D**) and quantification of CK19^+^ cells (**G** and **I**). Scale bars, 50 µm. Results show means ± SD of 3 independent experiments. At least 2 cell lines per genotype were tested. Each dot represents an independent experiment. *, *P* < 0.05, unpaired Student *t* test. **J,** Incucyte-based wound healing assay on *sgTgfb1* or *sgCtrl* CRISPRed AKC and AKPC tumor cells. Results show means ± SD of two cell lines per genotype used in in at least 3 independent experiments. *, *P* < 0.05; **, *P* < 0.01; ***, *P* < 0.001, 2way ANOVA. **K–M,** Viability assay analysis of oxaliplatin (**K**), 5-fluorouracil (**L**), and irinotecan (**M**) treatment in *sgTgfb1* or *sgCtrl* CRISPRed AKC and AKPC tumor cells. ns, not significant.

**Supplementary Figure S11.** Perturbing tumor–CAF dialog through TGF-β signaling interference improves FX therapeutic efficacy in orthotopic model of ATM-deficient PDAC. **A** and **B,** Immunofluorescence staining for αSMA (green) and histologic sections stained by picrosirius red on resected orthotopic tumors arising from *Atm^+/+^*; *Trp53^fl/fl^*; *LSL-Kras^G12D/+^*; *Ptf1a^Cre/+^* (KPC), *Atm^fl/fl^*; *Trp53^fl/fl^*; *LSL-Kras^G12D/+^*; *Ptf1a^Cre/+^*(AKPC) (**A**), *Atm^+/+^*; *Trp53^+/+^*; *LSL-Kras^G12D/+^*; *Ptf1a^Cre/+^* (KC), and *Atm^fl/fl^*; *Trp53^+/+^*; *LSL-Kras^G12D/+^*; *Ptf1a^Cre/+^* (AKC) (**B**) tumor cells treated or not with the TGFBR1 inhibitor galunisertib (galu.; 12.5 mg/kg) as in (Figure 6E). Cells were counterstained with DAPI (blue). Scale bars, 100 µm. **C,** Quantification of αSMA^+^ and picrosirius red-positive surface in resected orthotopic tumors shown in (**A** and **B**). Results show means ± SD. Each dot represents a mouse. *, *P* < 0.05; **, *P* < 0.01, unpaired Student *t* test. **D** and **E,** Kaplan-Meier analysis of survival of C57BL/6J mice orthotopically transplanted with KC (**D**) and KPC (**E**) tumor cells and treated or not with galu. and FX (50.0 mg/kg folinic acid, 25.0 mg/kg 5-fluorouracil, 25.0 mg/kg irinotecan, and 2.5 mg/kg oxaliplatin) from the orthotopic assay shown in (Figure 6E). ns, not significant, log-rank (Mantel-Cox) test. **F,** Magnetic resonance imaging of orthotopic AKPC tumor-bearing mice before treatment start (T0), during treatment (T1), and at endpoint (T2) treated or not with galu. and FX as in (Figure 6E). Red areas highlight pancreatic tumors. Tumor progression is indicated as percentage of tumor volume upon treatment normalized to initial tumor volume (T0). **G** and **H,** Immunohistochemistry staining for CC3 (**G** left) and Ki67 (**H** left) on resected orthotopic tumors arising either from KPC or AKPC cells treated or not with galu. and FX, from the orthotopic assay shown in (Figure 6E), and quantification of CC3^+^ (**G** right) and Ki67^+^ surface (**H** right) (**J**). Scale bars, 50 µm. *, *P* < 0.05; **, *P* < 0.01; ***, *P* < 0.001, unpaired Student *t* test. **I** and **J,** Immunohistochemistry staining for CC3 (top) and Ki67 (bottom) (**I**) on resected orthotopic tumors arising either from KC or AKC cells treated or not with galu. and FX, from the orthotopic assay shown in (Fig. 6E), and quantification of CC3^+^ (left) and Ki67^+^ surface (right) (**J**). Scale bars, 50 µm. *, *P* < 0.05; **, *P* < 0.01; ***, *P* < 0.001, unpaired Student *t* test. **K,** Immunofluorescence staining for PDPN (white) and Ly6C (green) (left) and quantification of PDPN^+^Ly6C^+^ cells (right) in resected orthotopic tumors of the assay shown in (Figure 6E). Cells were counterstained with DAPI (blue). Scale bars, 50 µm. Results show means ± SD of ≥ 2 AKC and AKPC mice. Each dot represents a mouse. **, *P* < 0.01, unpaired Student *t* test. **L,** Flow-cytometric analysis of αSMA^+^ myCAF/Ly6c^+^ iCAF ratio in resected orthotopic tumors arising from AKPC tumor cells treated or not with galu. and FX. Results show means ± SD. Each dot represents a mouse. *, *P* < 0.05; **, *P* < 0.01, unpaired Student *t* test.

**Supplementary Figure S12.** HR gene expression in human PDAC correlates with distinct CAF signature enrichments. A, UMAP embedding visualizing scRNA-seq data from PDAC patient (59), showing annotated cell clusters (left). Fibroblast subsets (myCAFs, iCAFs, non-CAFs) were identified based on CAF gene signatures from Elyada *et al.* (17). **B**, Pearson correlation coefficients between true and estimated cell proportions from Bayesian deconvolution using pseudo-bulk reference data (as in **A**). **C**, Analysis of the TCGA-PAAD dataset showing mutations frequencies (dot size) and statistical associations between recurrent driver genes (KRAS, TP53, CDKN2A, SMAD4) and DNA damage repair-related genes. **D**, Sanger sequencing and indel analysis of two independent *Atm^+/+^*; *Trp53^+/+^*; *LSL-Kras^G12D/+^*; *Ptf1a^Cre/+^* (KC) tumor lines following CRISPR/Cas9-mediated *Atr* gene editing in exon 4. Sequencing traces from CRISPR/Cas9-control and *sgAtr*-edited samples are depicted. **E,** Levels of reactive oxygen species (ROS) in *sgAtr* and *sgCtrl* CRISPRed KC. Results show means ± SD of 4 independent experiments. Two cell lines per genotype were used. Each dot represents an independent experiment. Unpaired Student *t* test. **F,** ELISA for TGF-β1 on conditioned medium (CM) from *sgAtr* and *sgCtrl* CRISPRed KC tumor cells. At least 2 CM batches per cell line were tested. Each dot represents a CM batch. *, *P* < 0.05; **, *P* < 0.01, unpaired Student *t* test. **G,** qPCR analysis of myCAF and iCAF marker gene expression in PSCs co-cultured for 72 hours with *sgAtr* and *sgCtrl* CRISPRed KC tumor cells. Results show means ± SD of ≥ 2 cell lines used in 2 independent experiments. Each dot represents a cell line. *, *P* < 0.05; **, *P* < 0.01, unpaired Student *t* test. **H,** Schematic illustration of the subcutaneous assay with treatment administration schedule (left) and time-dependent development over the course of 18 days (PDX-*ATR^mut^*) or 21 days (PDX-*ATR^wt^*) of subcutaneously engrafted tumors arising from 2 PDXs with representative hematoxylin and eosin. *, *P* < 0.05; **, 2-way ANOVA.

## SUPPLEMENTARY TABLE LEGENDS

**Table S1.** Cell count and proportion of cell types identified with the single-nucleus multiomics analysis.

**Table S2.** Clinical characteristics and mutations of patient-derived organoids

**Table S3.** Clinical characteristics and mutations of PDAC patients.

**Table S4.** CRISPR/Cas9 guide sequences. F, female; M, male; N/A, not available; PDAC, pancreatic ductal adenocarcinoma; TNM, tumor, nodes, metastasis classification.

