## Supplementary Figures for "Exploiting TGF-β-mediated Stromal Programming in Homologous Recombination-Deficient Pancreatic Cancer"

### Supplementary Fig. 1

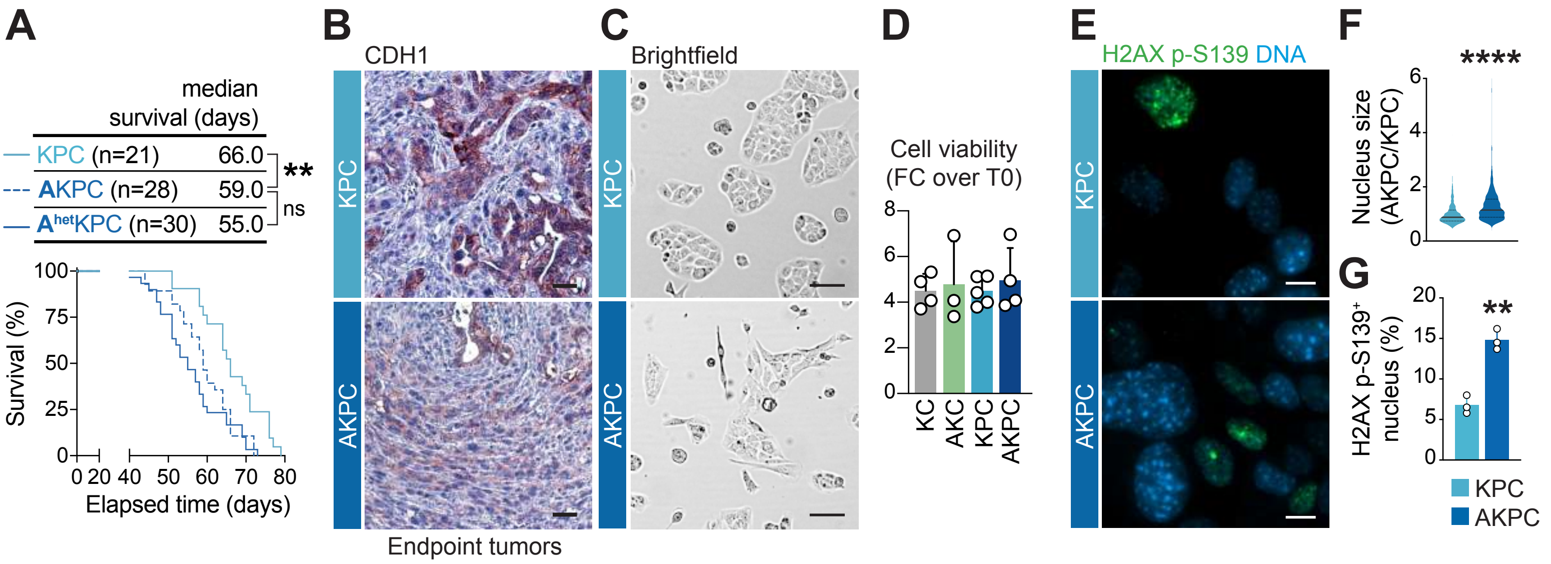

Supplementary Fig. 2

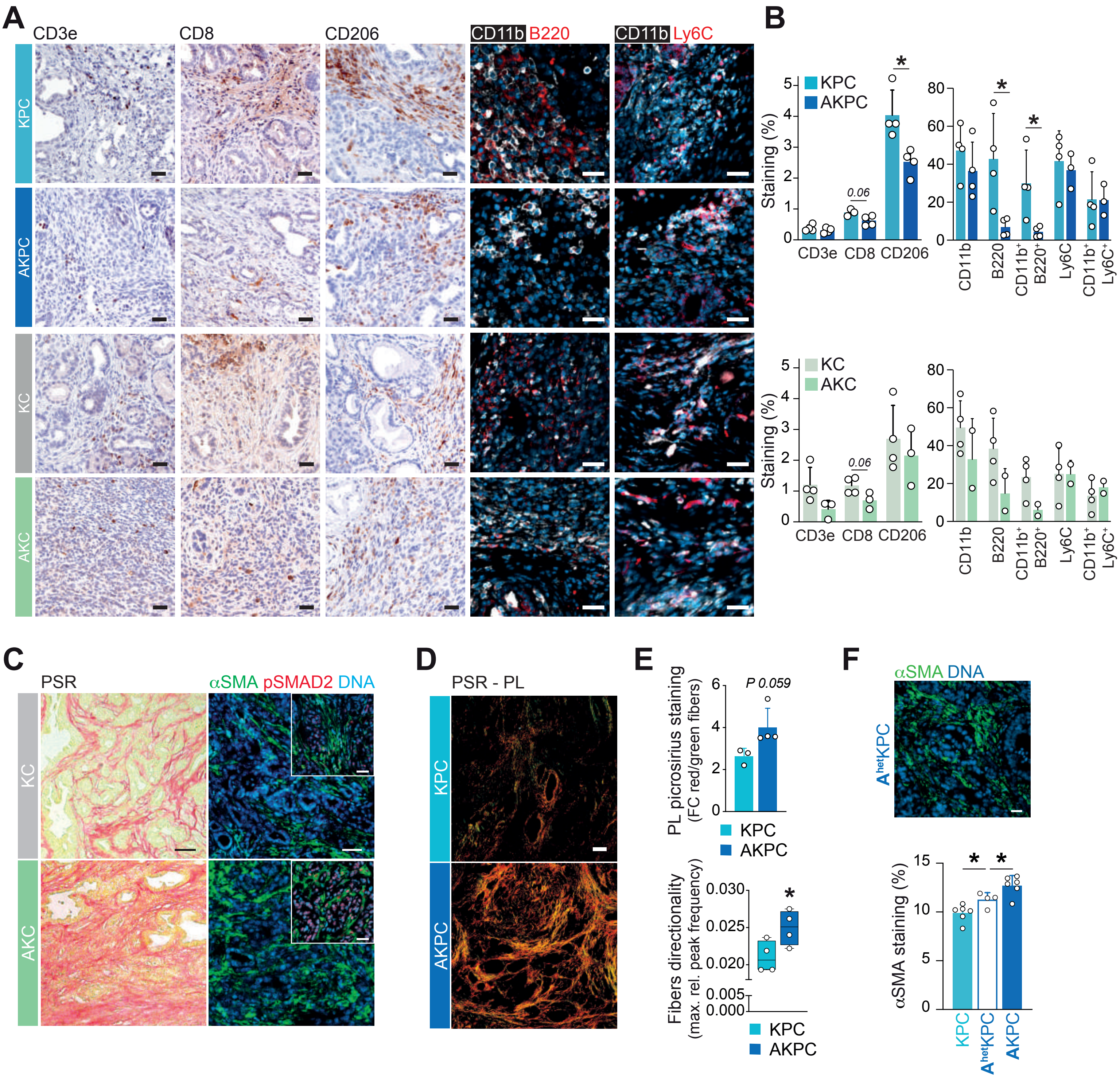

Supplementary Fig. 3

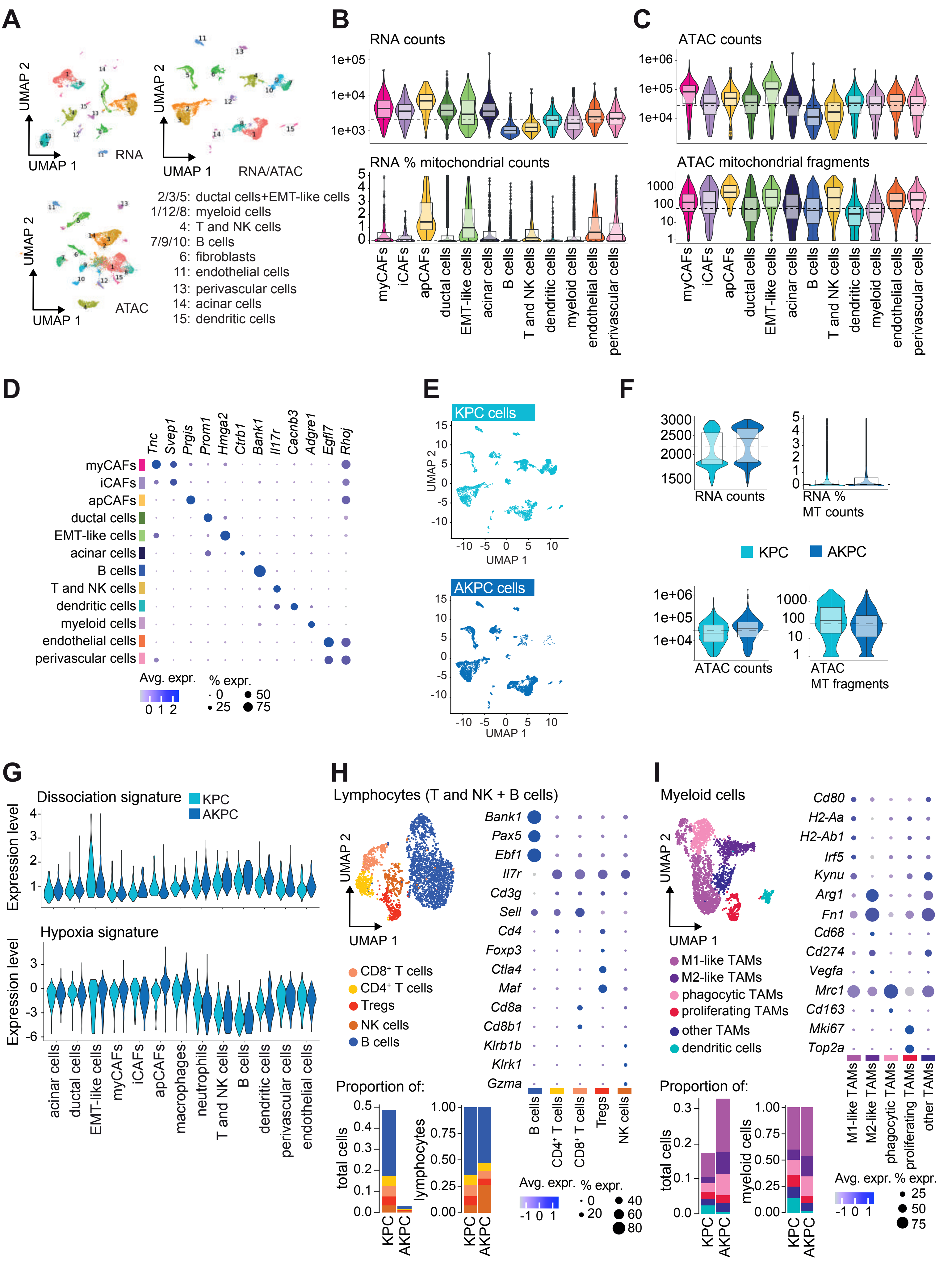

### Supplementary Fig. 4

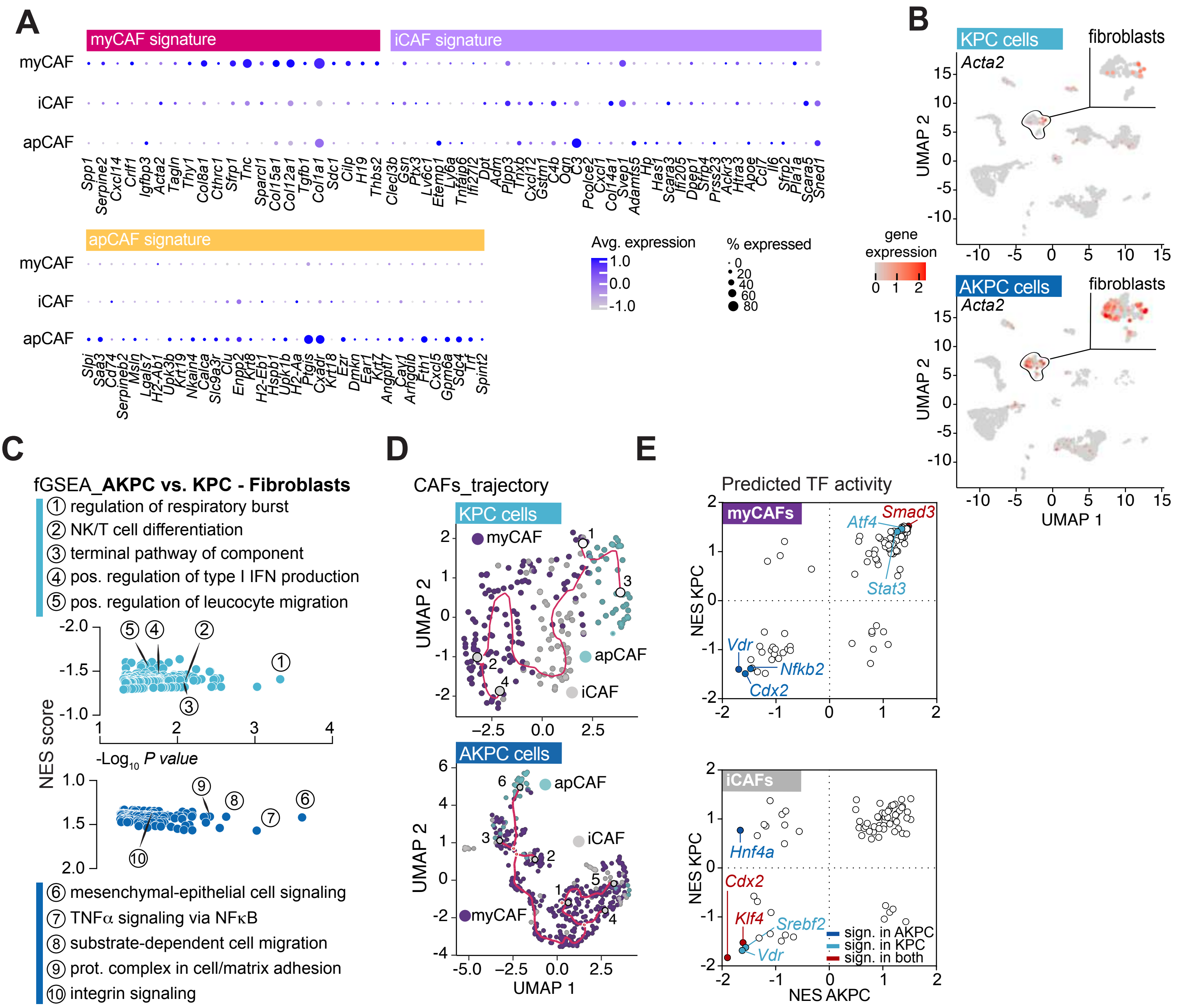

Supplementary Fig. 5

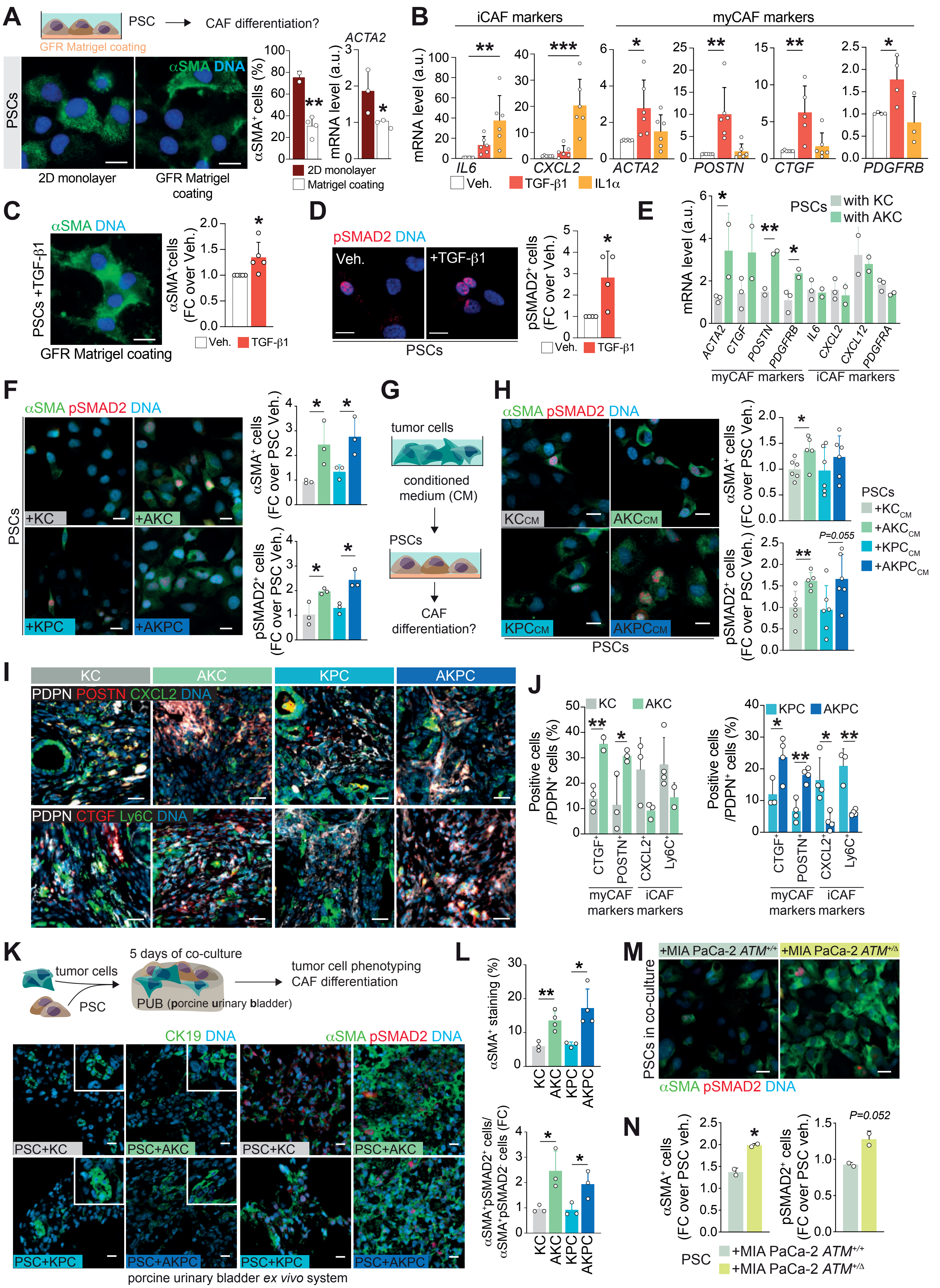

### Supplementary Fig. 6

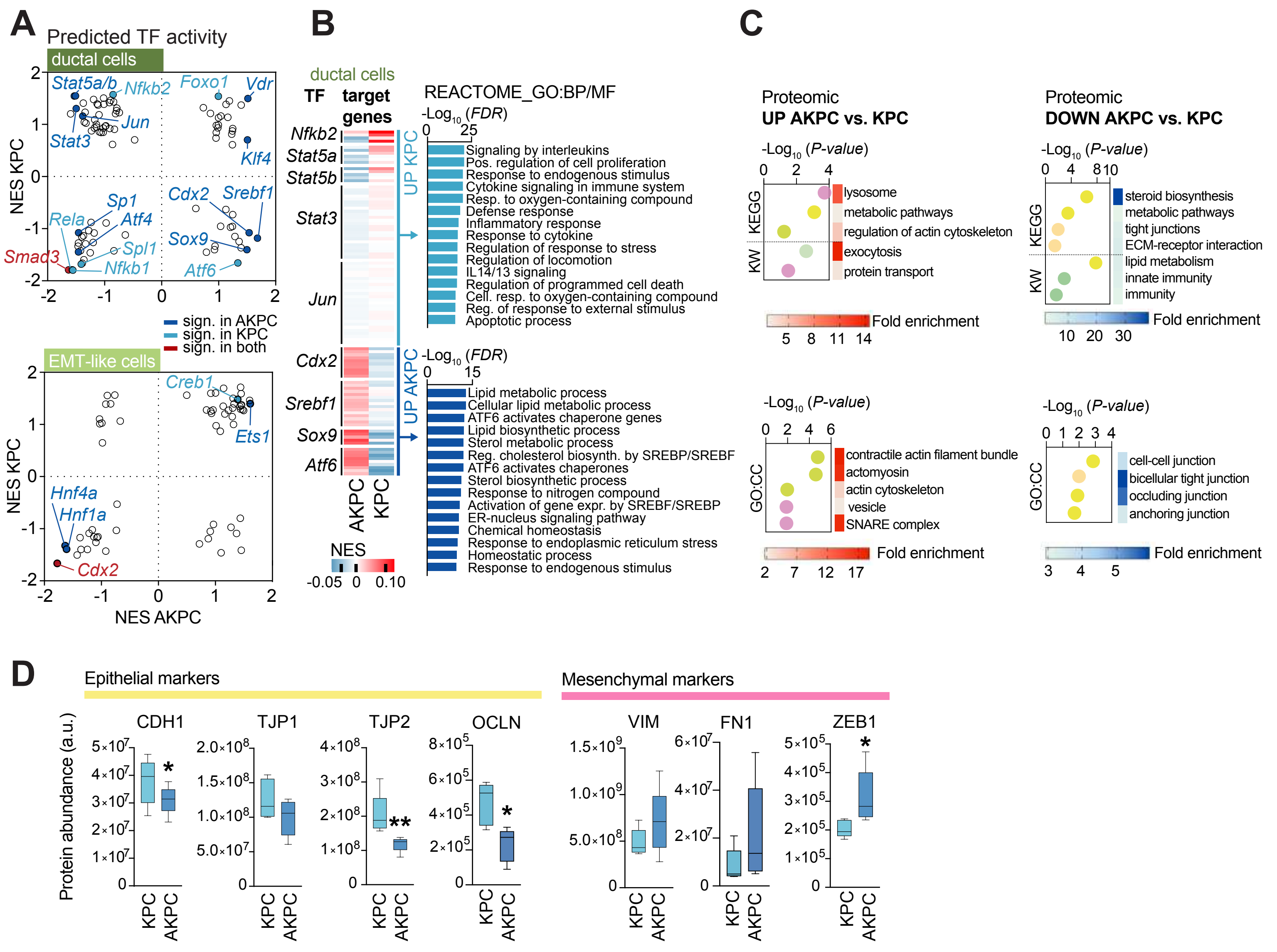

### Supplementary Fig. 7

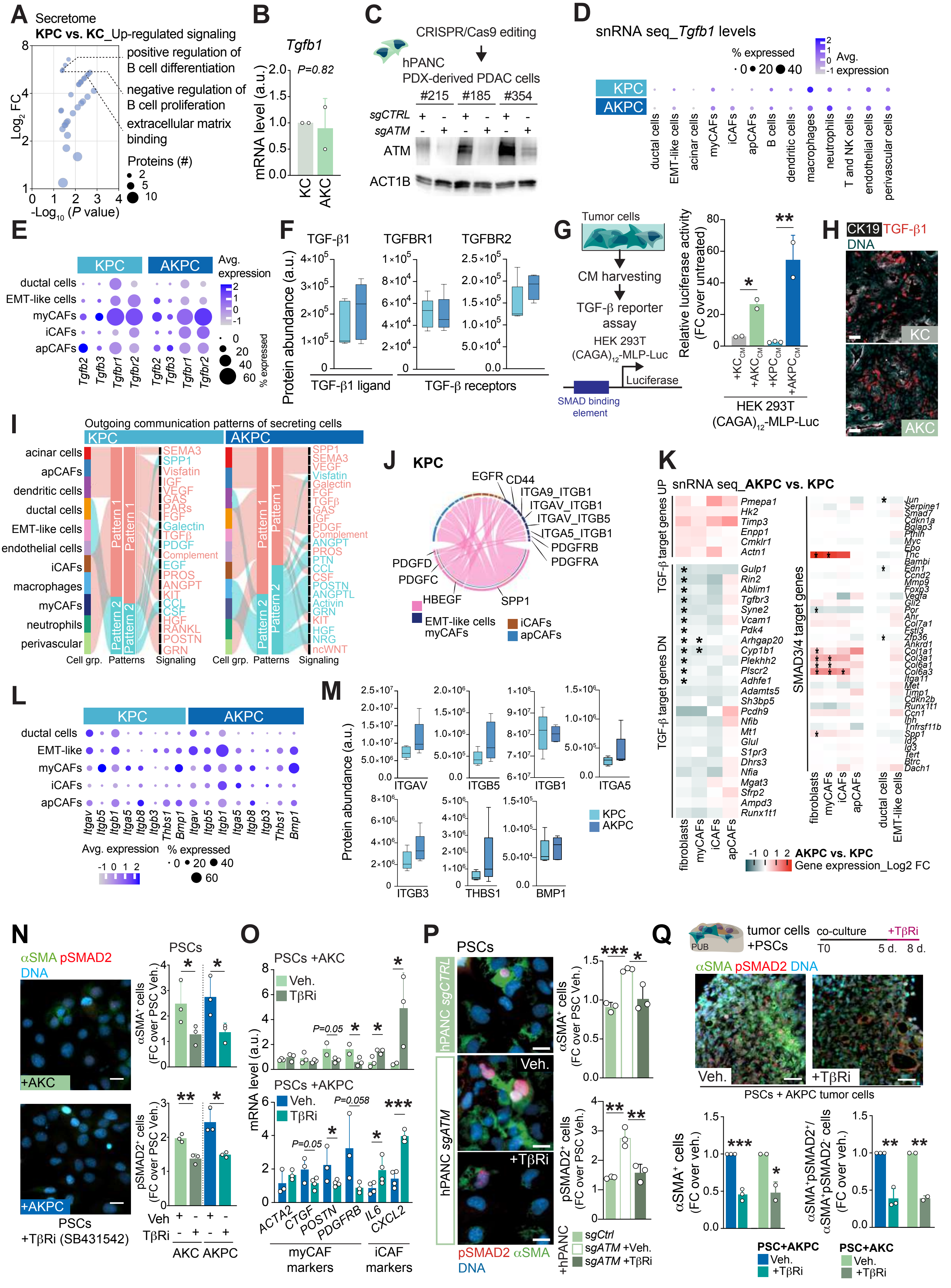

### Supplementary Fig. 8

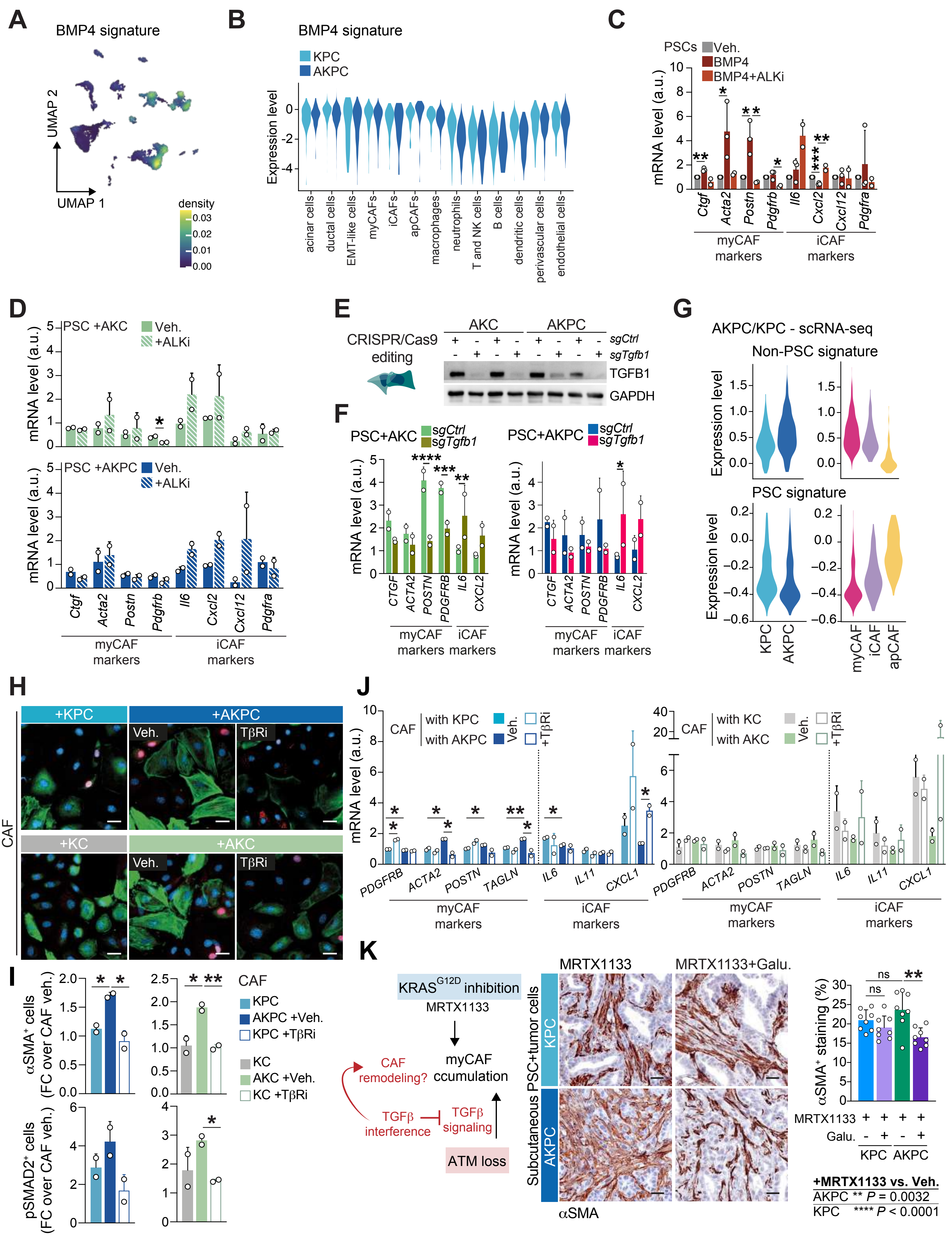

Supplementary Fig.9

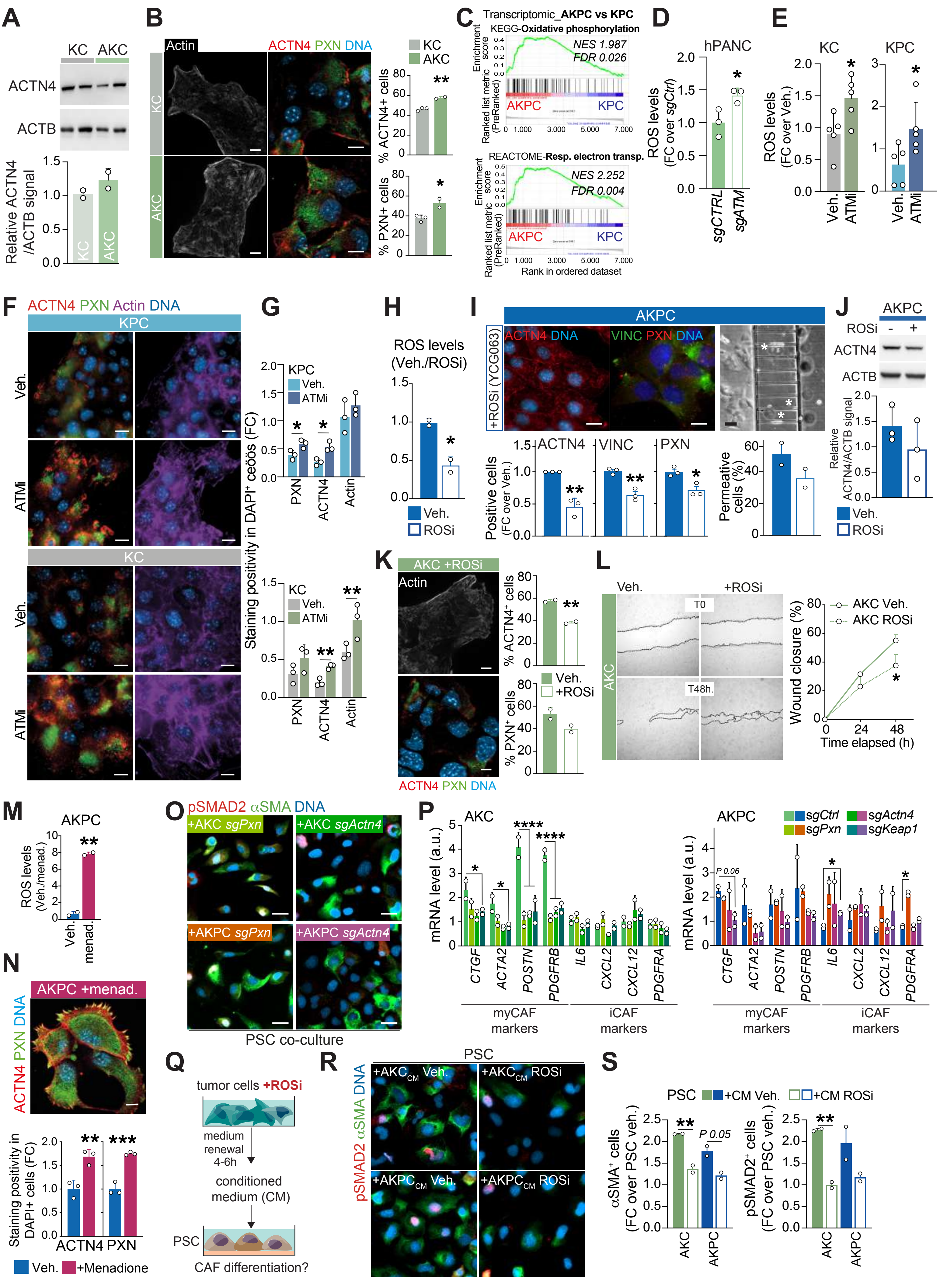

### Supplementary Fig. 10

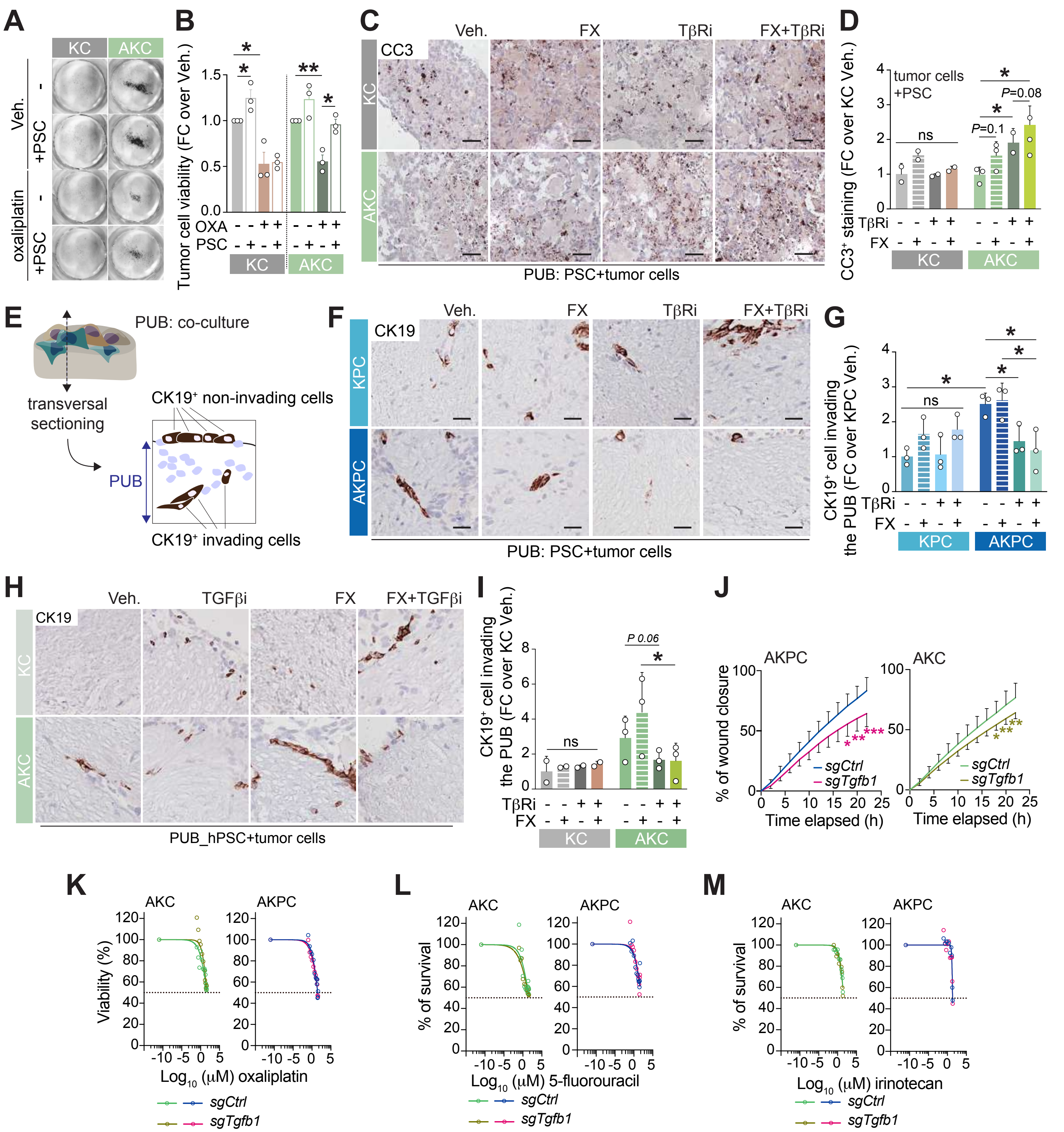

### Supplementary Fig. 11

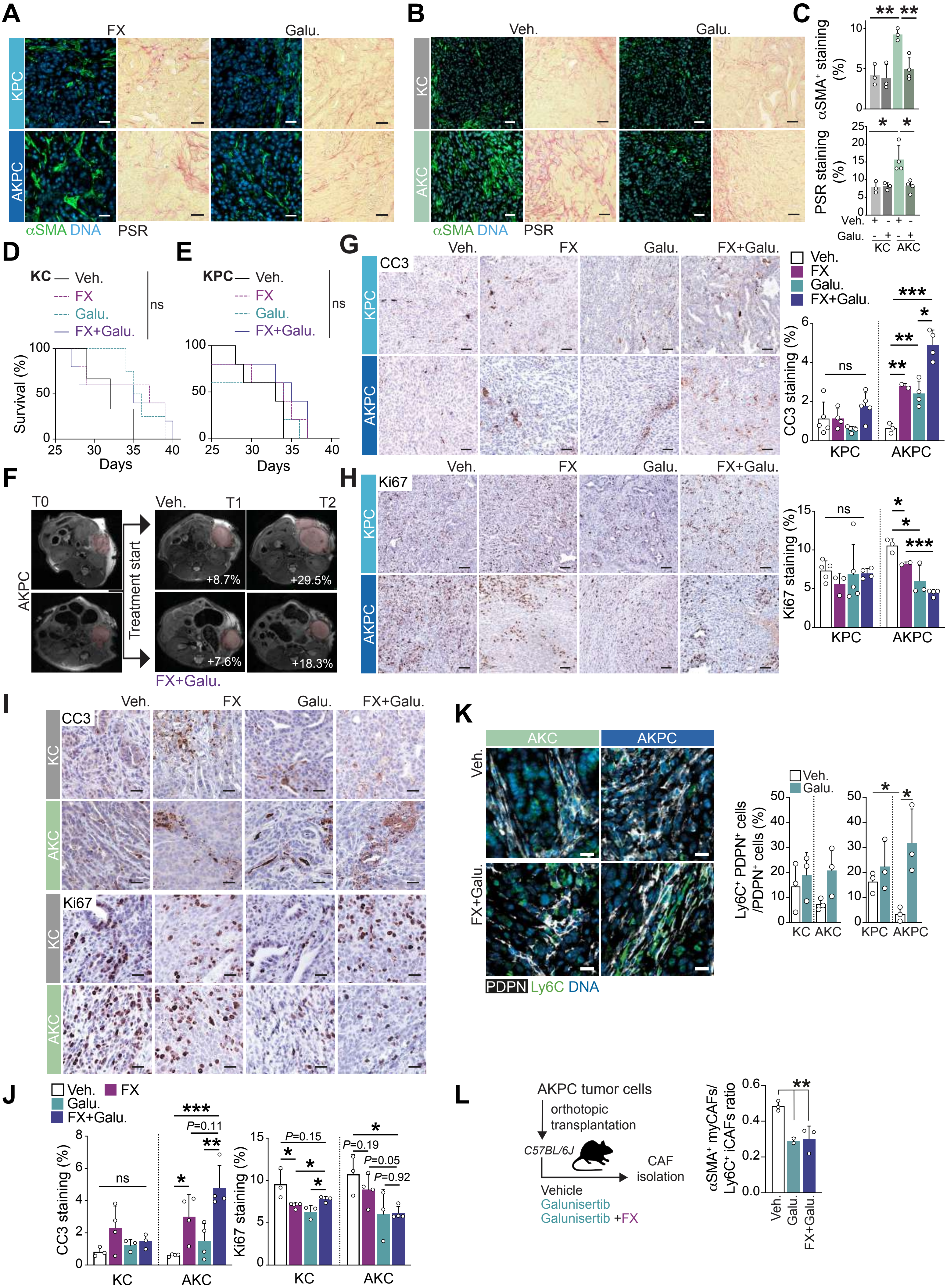

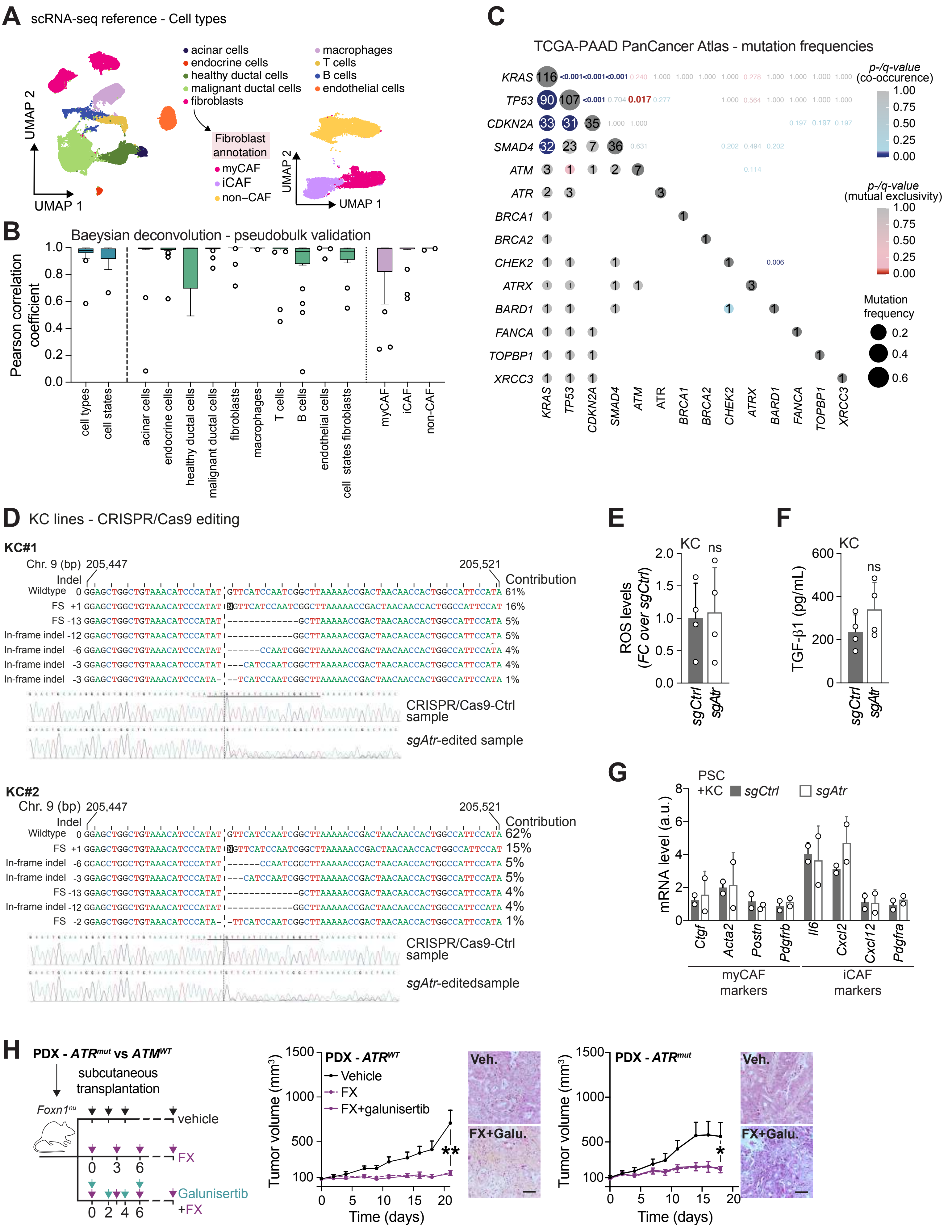
A
scRNA-seq reference - Cell types
acinar cells
endocrine cells
healthy ductal cells
malignant ductal cells
fibroblasts
macrophages
T cells
B cells
endothelial cells
Fibroblast annotation
myCAF
iCAF
non–CAF
UMAP 1
UMAP 2
B
Baeysian deconvolution - pseudobulk validation
Pearson correlation coefficient
cell types
cell states
acinar cells
endocrine cells
healthy ductal cells
malignant ductal cells
fibroblasts
macrophages
T cells
B cells
endothelial cells
cell states fibroblasts
myCAF
iCAF
non-CAF
C
TCGA-PAAD PanCancer Atlas - mutation frequencies
KRAS
TP53
CDKN2A
SMAD4
ATM
ATR
BRCA1
BRCA2
CHEK2
ATRX
BARD1
FANCA
TOPBP1
XRCC3
p-/q-value (co-occurence)
p-/q-value (mutual exclusivity)
Mutation frequency
D
KC lines - CRISPR/Cas9 editing
KC#1
Chr. 9 (bp) 205,447
205,521
Indel
Contribution
Wildtype 0 GGAGCTGGCTGTAAACATCCCATAT GTTCATCCAATCGGCTTAAAAACCGACTAACAACCACTGGCCATTCCATA 61%
FS +1 16%
FS -13 5%
In-frame indel -12 5%
In-frame indel -6 4%
In-frame indel -3 4%
In-frame indel -3 1%
CRISPR/Cas9-Ctrl sample
sgAtr-edited sample
KC#2
Chr. 9 (bp) 205,447
205,521
Indel
Contribution
Wildtype 0 62%
FS +1 15%
In-frame indel -6 5%
In-frame indel -3 5%
FS -13 4%
In-frame indel -12 4%
FS -2 1%
CRISPR/Cas9-Ctrl sample
sgAtr-editedsample
E
KC
ns
ROS levels (FC over sgCtrl)
sgCtrl
sgAtr
F
KC
ns
TGF-β1 (pg/mL)
sgCtrl
sgAtr
G
PSC +KC
sgCtrl
sgAtr
mRNA level (a.u.)
Ctgf
Acta2
Postn
Pdgfrb
Il6
Cxcl2
Cxcl12
Pdgfra
myCAF markers
iCAF markers
H
PDX - ATRmut vs ATMWT
subcutaneous transplantation
Foxn1nu
vehicle
FX
Galunisertib +FX
PDX - ATRWT
Vehicle
FX
FX+galunisertib
Tumor volume (mm3)
Time (days)
Veh.
FX+Galu.
PDX - ATRmut
